# Genome Position Does Not Impact Transgene Expression Efficiency in the Ancient Red Alga *Cyanidioschyzon merolae*

**DOI:** 10.64898/2026.01.31.703004

**Authors:** Kaian Teles, Gordon B. Wellman, Yuhan Zhang, Bárbara Bastos de Freitas, Viktor A. Slat, Martha R. Stark, Lecong Zhou, Perry J. Blackshear, Stephen D. Rader, Kyle J. Lauersen

**Author notes:** Corresponding Author: Kyle J. Lauersen.

## Abstract

The thermoacidophilic red alga *Cyanidioschyzon merolae* represents one of the simplest photosynthetic eukaryotes and an ancient divergent group in the primary endosymbiotic Viridiplantae. Because of its ∼16 Mbp genome, containing few introns, and capacity for transgene integration by homologous recombination, it is an emerging chassis for synthetic biology. However, genomic integration sites and scalable transformation methods have not been established to systematically investigate the effect of genome position on transgene expression. Here, we combined bioinformatic genome analysis, liquid-handling robotics, and assays of heterologous protein and metabolite production to establish a reproducible framework for nuclear genome engineering in *C. merolae*. We mapped and annotated 40 intergenic loci as candidate neutral sites across 16 out of 20 chromosomes and could validate 38 of them through robotic-assisted transformation. Reporter gene expression analysis revealed highly uniform expression at all integration sites across broad populations of transformants, indicating surprising minimal positional effects and transcriptional neutrality. The functional equivalence of these genomic ‘landing pads’ was determined by expression of a heterologous isoprene synthase, and coupling algal photobioreactors to headspace analysis to quantify isoprene production driven by transgene expression from different integration sites. Single copy transgene integrants, regardless of genome position, exhibited comparable reporter signals and consequent isoprene production. Together, these results provide the first experimentally validated set of neutral integration sites in *C. merolae* and establish a high-throughput transformation protocol for its genetic engineering in the context of synthetic genome biology.

## Introduction

Microalgae occupy an extraordinary range of ecological niches, thriving in environments that vary from polar oceans to acidic hot springs (Rappaport and Oliverio, 2023). Among them, members of the class *Cyanidiophyceae* represent one of the few photosynthetic eukaryotes adapted to life at both high temperature and low pH, often dominating microbial communities in geothermal and volcanic areas. Within this lineage, the unicellular red alga *Cyanidioschyzon merolae* 10D was originally isolated from the Phlegrean Fields near Naples, Italy (Toda et al., 1995). It is a thermoacidophilic photoautotroph with robust growth at 42–46 °C and pH 1–3, thriving under conditions that are extreme by mesophile standards. In addition, it has been shown adaptable to both freshwater and seawater (Hirooka et al., 2020; Villegas-Valencia et al., 2023). *C. merolae* is widely regarded as one of the simplest free-living eukaryotes, containing only a single copy of each major organelle (Miyagishima and Tanaka, 2021). Its ∼16 Mbp nuclear genome, the first photosynthetic eukaryotic genome completely assembled from telomere to telomere, encodes roughly 4,775 protein-coding genes with few introns and minimal genomic redundancy (Matsuzaki et al., 2004; Nozaki et al., 2007a; Schärfen et al., 2022). The fully sequenced genome, combined with its capacity for synchronized cell division and ease of cultivation in the lab, makes *C. merolae* uniquely suited for dissecting the molecular coordination of organelle replication, energy metabolism, and gene regulation in a minimal eukaryotic context (Imamura et al., 2015, 2010; Imoto and Yoshida, 2017; Misumi et al., 2005; Miyagishima et al., 2014; Mori et al., 2016; Moriyama et al., 2014a; Suzuki et al., 1994).

A defining feature of *C. merolae* as a model organism is its exceptional genetic tractability. Unlike model algae such as *Chlamydomonas reinhardtii*, *C. merolae* displays a high rate of homologous recombination (HR) integration events, enabling precise, marker-directed integration into the nuclear genome (Minoda et al., 2004). There is also no evidence yet reported of transgene silencing, a challenge in other algae (Watanabe et al., 2014; Zienkiewicz et al., 2019). These characteristics make *C. merolae* an excellent system for precise genome engineering and long-term heterologous pathway expression.

From a biotechnological perspective, *C. merolae* offers several advantages for large-scale cultivation and metabolic engineering. Its robust growth in defined mineral media, and tolerance to high light and heat, support consistent biomass production in the laboratory, outdoors, and in pilot-scale photobioreactors (Hirooka et al., 2020; Villegas-Valencia et al., 2023). The species naturally accumulates valuable bioproducts including thermostable phycocyanin, carotenoids (without alpha carotenes), and β-glucans (Lang et al., 2022; Rahman et al., 2017). Its lack of a rigid cell wall facilitates both DNA delivery and product recovery from biomass (Miyagishima and Tanaka, 2021). Recent studies have demonstrated its metabolic engineering potential, including triacylglycerol overproduction without growth penalty (Sumiya et al., 2015), heterotrophic growth enabled by a *Galdieria sulphuraria* sugar transporter (Fujiwara et al., 2019), the production of non-native ketocarotenoids (Seger et al., 2023), and heterologous isoprene production (Villegas-Valencia et al., 2025). Recently, specific genetic tools have been developed for *C. merolae*, including rapamycin-inducible knockdown systems (Fujiwara et al., 2024), multiplex organelle imaging platforms (Tanaka et al., 2021), and multiple classes of inducible promoters, encompassing chemical inducers (estradiol, IPTG) (Fujiwara, n.d.), nutrient-responsive regulation (nitrogen source) (Fujiwara et al., 2015), and stress-responsive heat-shock promoters (Sumiya et al., 2014).

Despite these advances, a major bottleneck in *C. merolae* engineering is the absence of standardized genomic integration sites and an understanding of their influence on transgene expression. In many eukaryotes, transgene performance is strongly influenced by genomic position, as local chromatin state, replication timing, and proximity to regulatory elements modulate expression efficiency (Brady et al., 2020; Cabrera et al., 2022). However, this has not been systematically characterized in *C. merolae*. Identifying transcriptionally neutral ‘safe-harbor’ loci, genomic regions that permit stable, uniform transgene expression without perturbing essential functions, is essential for establishing predictable engineering frameworks, especially in the context of emerging synthetic genomic efforts (Babaei et al., 2021; Bai Flagfeldt et al., 2009; Jensen et al., 2014; Juhas et al., 2014).

To enable a systematic evaluation of genomic position effects, we established a robotics-assisted transformation protocol. Using this platform, we performed a genome-wide identification and mapping of potential neutral integration loci in the nuclear genome of *C. merolae*. By integrating bioinformatic screening with reported expression assays and heterologous metabolite quantification, we determined that position effect does not influence the level of transgene expression in the nuclear genome of this alga. This approach provides the foundation for high-throughput engineering efforts in this host, which will be valuable for future synthetic genomics applications.

## Materials and Methods

### Neutral Site Mapping and Visualization

For RNA sequencing (RNA-seq), *Cyanidioschyzon merolae* 10D cells were cultured, total RNA was extracted and sequenced as previously described by (Schärfen et al., 2022), with a minor modification in which cells were grown in MA2G medium (MA2 supplemented with 50 mM glycerol, as described below). Raw RNA-seq reads were aligned to the *C. merolae* 10D reference genome (Matsuzaki et al., 2004; Nozaki et al., 2007b) obtained from Ensembl Plants release 46 (Howe et al., 2020) using STAR v2.7.3a (Dobin et al., 2013). STAR was configured to output stranded wiggle signal files containing unnormalized raw read counts for both uniquely mapping and multi-mapping reads. Wiggle files were subsequently converted to BigWig format using wigToBigWig v2.8 (Kent et al., 2010).

Genomic coordinates of putative neutral loci were identified using a custom Python script developed in-house (code available at https://github.com/gbwellman/Cm_NS_finder). The reference genome of *Cyanidioschyzon merolae* 10D was used as input with mapped RNA-seq data (Nozaki et al., 2007a). Intergenic spaces were filtered stepwise for flanking genes in a 3′ to 3′ orientation, both flanking genes had evidence of transcription based on RNA-seq and contained a ‘neutral region’ (>10 bp) with no mapped RNA-seq reads indicating no overlapping 3′UTRs. This resulted in 39 putative loci. Manual inspection was carried out to ensure identified loci were not located close to centromeric regions. Genomic sequences, 3000 bp around the ‘neutral region’ were exported for plasmid design. The previously used neutral site (CMD184C – CMD185C) was also included, although it did not meet the criteria for selection. *Bbs*I and *Zra*I restriction sites were removed by addition of one base change where needed from the 1100 bp surrounding the center each locus to form the 550 bp homology arms for integration. Partial upstream *Zra*I sites were identified to enable linearization by blunt-end restriction enzyme digest for transformation. Therefore, integrated HR regions in each plasmid are variable in length by several nucleotides. Sequences were synthesized by Genscript and subcloned into pUC57-KanR. Homology arm sequences were identified and mapped to their corresponding genomic loci for visualization (Supplementary Fig. 1A).

### Plasmid Construction

Plasmids were designed for integration of chloramphenicol acetyltransferase (CAT), *mVenus* and/or isoprene synthase (*IspS*) into the intergenic neutral loci identified above. A complete list of gene elements used *in silico* designs is provided in Supplementary File 1, and all plasmids are available upon request from the authors or with permission from Genscript.

Endogenous elements (promoters, terminators, homology arms) were retrieved from the *C. merolae* 10D reference genome (Fujiwara et al., 2019, 2017; Moriyama et al., 2014b). The *S. aureus* CAT (NCBI M58516.1 (Schwarz and Cardoso, 1991)) was used as a selectable marker, and the *Ipomoea batatas* isoprene synthase (*IbIspS*; NCBI AZW07551.1 (Ilmén et al., 2015)) was used for isoprene biosynthesis as previously described (Villegas-Valencia et al., 2025). Coding sequences were back-translated and codon-optimized according to *C. merolae* nuclear codon usage table found in the Kazusa database (https://www.kazusa.or.jp/codon/cgi-bin/showcodon.cgi?species=280699). All sequences were synthesized and subcloned by Genscript (Piscataway, USA). Plasmids were delivered lyophilized, transformed into *E. coli* DH5α using kanamycin (50 µg/mL) or ampicillin (100 µg/mL) selection, and maintained in lysogeny broth (LB).

### Robotics

All scripting for the Opentrons OT-2 liquid-handling (Opentrons, USA) robot was developed in Python and made available as open-source code (below). Custom protocols were executed through the Opentrons App, which was used to load user-defined scripts, labware definitions, and calibration parameters. The OT-2 hardware configuration included a multichannel P300 and a multichannel P20 Gen 2 pipette, Gen 1 Heater-Shaker Module, Gen 1 Temperature Module, Deep-Well Metal Adapter, and standard Opentrons P20 and P300 tip racks all from Opentrons, USA. For automated colony isolation, a PIXL robotic colony picker (Singer Instruments, USA) was used. Detailed operational parameters and settings for PIXL are described in Supplementary Table 1 and detailed protocols and code for OT-2 are available at https://github.com/KaianTeles/ot2-cmerolae-pipeline.git.

### Algal Culture

The wild-type *C. merolae* 10D strain (NIES-3377) was obtained from the National Institute for Environmental Studies (NIES, Japan). Cultures were maintained in MA2G medium, a modified version of Modified Allen’s medium (‘MA’, (Minoda et al., 2004)) doubling (‘2’) concentrations of several components. MA2G contains: 40 mM (NH₄)₂SO₄, 8.0 mM KH₂PO₄, 4.0 mM MgSO₄, 1.0 mM CaCl₂·2H₂O, 100 μM FeCl₃·6H₂O, 80 μM Na₂EDTA·2H₂O, 184.4 μM H₃BO₃, 36.4 μM MnCl₂·4H₂O, 3.08 μM ZnCl₂, 6.4 μM Na₂MoO₄·2H₂O, 0.68 μM CoCl₂·6H₂O, 1.28 μM CuCl₂·2H₂O, and 50 mM glycerol (‘G’). The pH was adjusted to 2.5 with H₂SO₄. Glycerol was used to promote faster growth as it enhances respiratory activity (Moriyama et al., 2015).

Cultures were either kept on MA2G + 250μg/mL chloramphenicol gellan gum plates with 10μL corn starch beds or liquid cultures with MA2G incubated under continuous illumination (90–130 μE) at 42 °C in a Percival incubator AL-30 (Geneva Scientific, USA) with 5 % CO₂ supplementation and agitation at 120 rpm.

### Transformation of *C. merolae* 10D

Transformation was performed as previously described with minor modifications (Fujiwara et al., 2017; Ohnuma et al., 2014; Villegas-Valencia et al., 2025; Zienkiewicz et al., 2019). Linear DNA (1 pmol) was generated by PCR using Platinum™ SuperFi™ DNA Polymerase (Invitrogen, USA) (Supplementary Table 2). PCR products were purified using ZR-96 DNA Clean & Concentrator columns (Zymo Research, USA), resuspended in nuclease-free water, and quantified using a NanoDrop One spectrophotometer (Thermo Fisher Scientific, UK).

Exponentially growing cultures were maintained in MA2G medium at 42 °C, 90–130 μE light, with 5% CO₂. One day before transformation, cells were diluted to OD₇₄₀ ∼ 0.2 and grown overnight to OD₇₄₀ ∼0.8–1.0. PEG4000 (Alfa Aesar) was prepared by dissolving 0.9 g PEG in 750 μL MA-I medium and incubated at 42 °C until fully solubilized. The medium ‘MA-I’ is composed of (20 mM (NH₄)₂SO₄, 2 mM MgSO₄, 92.2 μM H₃BO₃, 18.2 μM MnCl₂·4H₂O, 1.54 μM ZnCl₂, 3.2 μM Na₂MoO₄·2H₂O, 0.34 μM CoCl₂·6H₂O, 0.64 μM CuCl₂·2H₂O, pH 2.5).

Cells were concentrated ∼250-fold by centrifugation (2000×g, 10 min) and resuspended in warm MA-I. The DNA mixture consisted of 84 μL linear DNA solution, 10 μL 10X MA-I, and 6 μL 10 mg/mL denatured salmon sperm DNA (Invitrogen, USA). Twenty-five μL of concentrated cells were added, followed by 125 μL PEG solution, and mixed gently by flicking 8–10 times. The mixture was immediately diluted with 1 mL warm MA2G and transferred into 50 mL MA2G for recovery (48 h, 42 °C, 120 rpm). Transformations were also done using OT-2 liquid handler, detailed protocol are available in (https://github.com/KaianTeles/ot2-cmerolae-pipeline.git).

For starch-bed plate preparation, gellan gum plates were prepared as described in (Villegas-Valencia et al., 2025), with the following modifications. Gellan gum was poured (∼50 mL per plate) into PlusPlates (Singer Instruments, USA) and allowed to solidify inside a safety cabinet at room temperature. After solidification, plates were weighed. A 20% (w/v) starch solution in MA2G-C medium was prepared as described in (Villegas-Valencia et al., 2025), and 40 mL of this solution was transferred to an OT-2–compatible 90 mL reservoir. To adjust Z-axis offset, plate weights were provided as input to a custom OT-2 API, and the OT-2 protocol was executed to dispense starch onto the gellan gum plates, resulting in an array of 96 starch spots (10 µL per spot) per plate.

Recovered cells were pelleted and resuspended in 2.7 mL conditioned MA2G-C medium (pre-grown wild-type culture medium, OD₇₄₀ ≈ 1.0, filter-sterilized (Villegas-Valencia et al., 2025)). MA2G-C was used both for preparing starch plates and for diluting cells prior to colony formation. Cell suspensions were manually transferred into a 96-well plate using a 200 µL multichannel pipette. A custom OT-2 script was then used to generate serial dilutions, producing neat, 1:3, 1:9, and 1:27 dilutions. Following dilution, the OT-2 was used to plate cells onto starch-bed plates, applying the same weight-based Z-axis adjustment method described above. After cell plating, plates were sealed with grafting tape and incubated at 42 °C under 100 µE light intensity.

### Molecular Screening and Genotyping

After 9 –14 days, small circular green colonies were visible, and 22 colonies from each transformation were robotically picked using the PIXL (Singer Instruments, USA) colony-picker and inoculated into 200 μL MA2G medium with 250 μg/mL chloramphenicol in 96-well plates. After ∼5 d, liquid cultures were re-plated on starch beds with antibiotic selection and incubated ∼4 d.

All 22 colonies from the previous step were picked from starch beds and PCR was performed for left and right homology arms, using Platinum II Taq Hot-Start DNA Polymerase (Thermo Fisher Scientific, USA) according to the manufacturer’s instructions. Genotyping primers were designed to anneal ∼100 bp outside each pair of homology arms (Supplementary Table 3) and inside the transformed cassette.

For quantitative analysis, qPCR was conducted on a CFX96 Real-Time PCR System (Bio-Rad) using SsoAdvanced Universal SYBR Green Supermix (Bio-Rad, USA) according to the manufacturer’s instructions. Copy number of the *mVenus* gene within each plasmid was normalized to *C. merolae* 60S rDNA (NCBI: 16997147). DNA was extracted as described in (Hu and Lagarias, n.d.). Reactions (20 μL) included no-template controls. Primer sequences were validated in (Villegas-Valencia et al., 2025) and are provided in Supplementary Table 4.

### Flow Cytometry

For benchmarking of *mVenus* reporter expression, 10 colonies per construct confirmed by PCR (both left and right homology arm positive) were selected. Cultures were grown in 6-well plates to stationary phase, diluted to OD₇₄₀ ∼ 0.2, and incubated overnight under growth conditions described above. Cells (5 µL) were diluted in 0.9 % saline solution and analyzed using an Attune NxT flow cytometer (Thermo Fisher Scientific, UK). Data were processed in FlowJo software and gating was employed to exclude doublets and debris (Supplementary Fig. 2). Median fluorescence intensity (BL-1 channel) was used for quantification.

### Fluorescence Imaging

Transformants grown on starch-bed plates were screened using a ChemStudio PLUS imager (Analytik Jena, USA) equipped with appropriate filters for *mVenus* and chlorophyll channels as previously described (Gutiérrez et al., 2022). Images of all transformants were acquired for both channels (Supplementary Fig. 3A).

For in-gel fluorescence, 1 mL of mid-log cells were centrifuged (2000 × g, 10 min), and the pellet either snap-frozen (−80 °C) or resuspended in ∼200 μL sample buffer (0.2 M SDS, 0.3 M Tris, 30 % glycerol, 0.02 % bromophenol blue). Samples were incubated 5 min at 40 °C and cooled on ice prior to SDS-PAGE. Fifteen μL of sample and 4 μL PageRuler Plus Prestained Ladder (Thermo Fisher Scientific, UK) were loaded per lane. Gels were visualized in the ChemStudio PLUS using appropriate filters as previously described (Gutiérrez et al., 2022).

### Quantification of Heterologous Isoprene Biosynthesis

To assess *Ib*IspS activity and isoprene production, 2 transformants of each nuclear locus tested were cultured in 400 mL MA2G in Algem photobioreactors (Algenuity, UK). Copy numbers of transformed cassettes were quantified by qPCR in each transformant to ensure single-copy insertions (Supplementary Fig. 4). Cultures were grown at 42 °C and 120 rpm under constant light (400 μE). For each transformant tested, individual precultures were grown in 50 mL MA2G medium under the same cultivation conditions described above. Following preculture growth, cells were diluted into 400 mL of fresh MA2G medium to a starting optical density of OD₇₄₀ ≈ 0.4 for subsequent experiments.

Headspace isoprene was quantified as previously described (Villegas-Valencia et al., 2025), using a real-time Hiden Analytical HPR-20 R&D mass spectrometer equipped with a multi-port gas inlet. Cultures were sparged with 3 % CO₂ (25 mL/min), and off-gas was passed through CaCl₂ desiccant and analyzed sequentially (∼3.5 min per sample). Isoprene was monitored at m/z 67, 68, 53, 39, 40, 41, and 27, with background gases deconvoluted by QGA Software (v2.0.8). A detailed protocol can be found in reference (Villegas-Valencia et al., 2025).

### Statistical Analysis

Data analysis was conducted using biological and/or technical replicates, as specified in each figure legend. Mean values and standard errors of the mean (SEM) were calculated and illustrated in the corresponding graphs. Statistical analyses were performed using in-house Python scripts implementing standard procedures, including one-way or Welch’s ANOVA followed by Tukey’s HSD post-hoc test, with the NumPy, Pandas, SciPy, and StatsModels libraries. Differences were considered statistically significant at *p* < 0.05. Custom analysis scripts are available upon request.

## Results and Discussion

### Genome-wide identification and mapping of neutral integration sites

Neutral integration sites across the nuclear genome were identified and consequently investigated experimentally. Candidate loci were selected through bioinformatic analysis of a deep-sequenced transcriptomic data set (∼800 million reads for ∼5,000 genes, or ∼100 reads of each transcribed base pair) mapped to the *C. merolae* reference genome (Ensembl v46), allowing sensitive detection of even minimal transcription. Deep transcriptomic raw sequencing data of *Cyanidioschyzon merolae* strain 10D (WT) has been deposited with the NCBI under SRA accession SRR36442998. Along with transcribed regions, we excluded regions overlapping coding sequences, essential genes, or known regulatory elements (Fig. 1A). Centromeric regions were defined based on previously mapped centromere positions (Kanesaki et al., 2015), and adjacent zones were masked to prevent unwanted recombination events. Homology arms of ∼1 kb were extracted from the flanking regions of each candidate locus, and a custom Python script (https://github.com/KaianTeles/ot2-cmerolae-pipeline.git) was developed to automate mapping, visualization, and annotation of all selected sites relative to chromosomal features and centromere coordinates.

**Figure 1.**
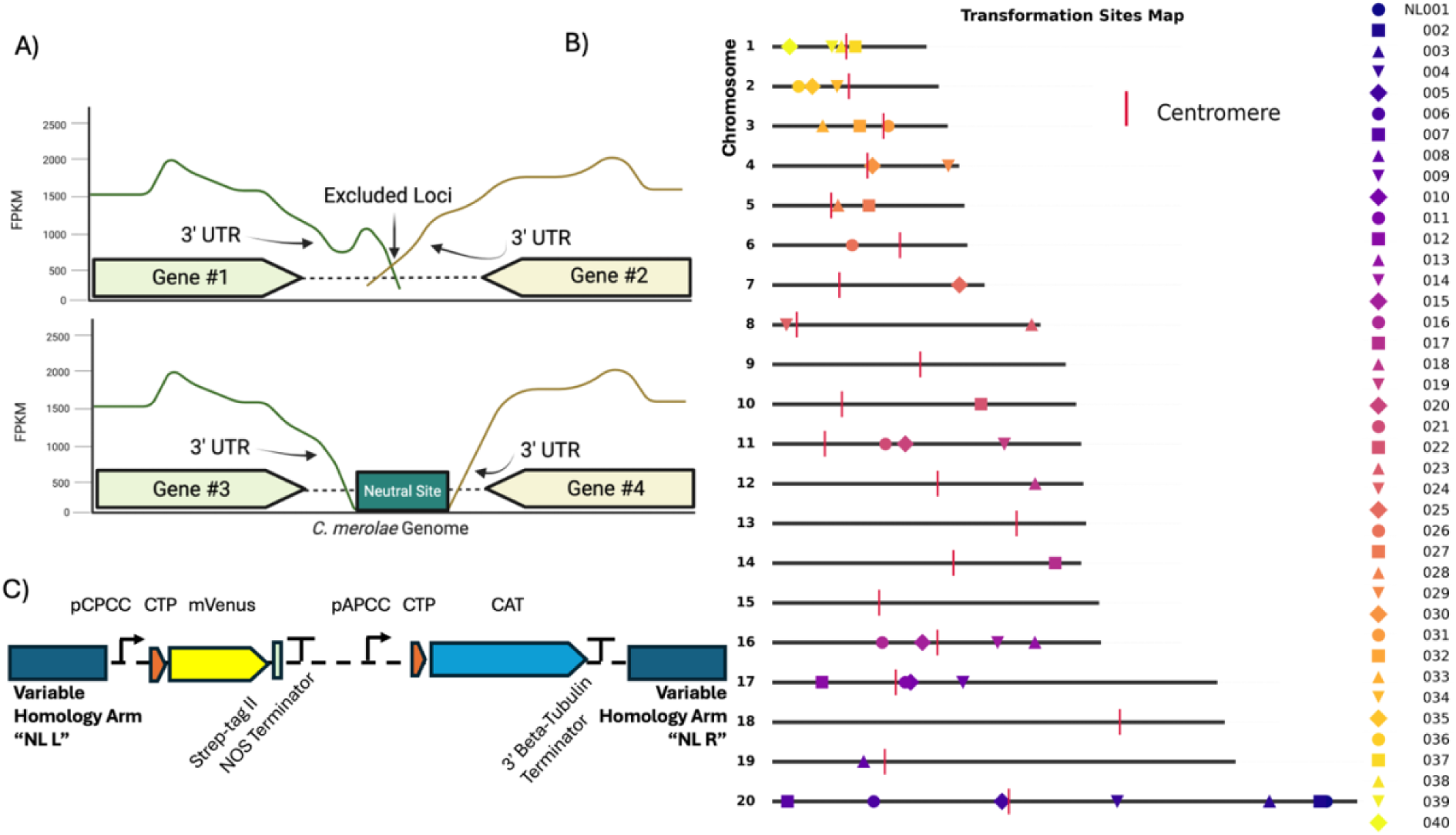
Identification of transcriptionally neutral genomic loci and design of integration constructs in *C. merolae*. (A) Schematic representation of the criteria used to identify putative neutral sites. Intergenic regions flanked by convergent 3′ UTRs with minimal or no transcriptional readthrough (bottom) were considered suitable for targeted integration, whereas regions showing detectable readthrough or overlapping transcriptional activity were excluded (top). RNA-seq coverage (FPKM) for each gene pair is shown to illustrate transcriptional boundaries. (B) Chromosomal map of the 40 candidate neutral loci identified across *C. merolae* chromosomes. Each symbol represents a mapped candidate locus (NL001–NL040), color-coded by site identifier. Centromeres are shown as red vertical markers. The relative genomic positions illustrate the broad and uniform distribution of neutral sites across the nuclear genome. (C) Schematic of the standardized reporter plasmid used for locus validation. Each construct contains a left and right homology arm matching a specific candidate locus, the constitutive *pCPCC* promoter driving *mVenus* fused to a Strep-tag II followed by the *NOS* terminator, and a chloramphenicol-resistance cassette (*CAT*) driven by the *pAPCC* promoter and terminated by the *3′ β-tubulin* terminator.

This analysis yielded 39 candidate ‘neutral’ loci (NL), without gene-dense or repetitive regions. NL were not found in chromosomes 9, 13, 15 and 18 (Fig. 1B, Supplementary Fig. 1A and Supplementary File 1). In addition, we included the previously used integration site on Chromosome 4 between CMD184C and CMD185C (hereafter referred to as NL029) as a benchmark control, given its widespread use in *C. merolae* transformation studies (Fujiwara et al., 2021, 2015, 2013; Seger et al., 2023; Villegas-Valencia et al., 2025). Although this locus did not meet our neutrality criteria due to detectable transcriptional readthrough (Supplementary Fig. 1B), it was retained for comparison to prior work. The mapped sites were validated with PCR (Supplementary Fig. 1C) and catalogued and used in subsequent integration experiments. For these experiments, we designed a set of reporter plasmids in which *mVenus* was placed under a constitutive promoter and flanked by the locus-specific homology arms corresponding to each mapped site (Fig.1C). The results of transformation of these plasmids are described in the following section.

### Development of a high-throughput automated transformation platform for *C. merolae*

Transformation of *C. merolae* has relied on fully manual – artisanal – workflows that are slow, labor-intensive, and prone to variability. Conventional transformation involves several consecutive steps, including PEG incubation, cell dilutions, preparation and plating of starch beds on gellan gum, and colony picking, each requiring manual handling (Villegas-Valencia et al., 2025). These manual operations limit scalability and complicate the comparison of constructs across experiments or individual operators.

To improve on this process, we developed a high-throughput transformation pipeline integrating an OT-2 liquid-handling robot and a PIXL robotic colony picker. The OT-2 was adapted to execute the PEG-mediated transformation steps and automate additional stages of the workflow, including preparation of starch beds, cell dilutions, plating of transformation, and subsequent colony genotyping (Fig. 2). Independent automation protocols were designed for each process. Custom scripts were developed for aspiration speeds, mixing cycles, and z-axis offsets, to enable gentle culture handling and uniform cell distribution during PEG exposure and plating.

**Figure 2.**
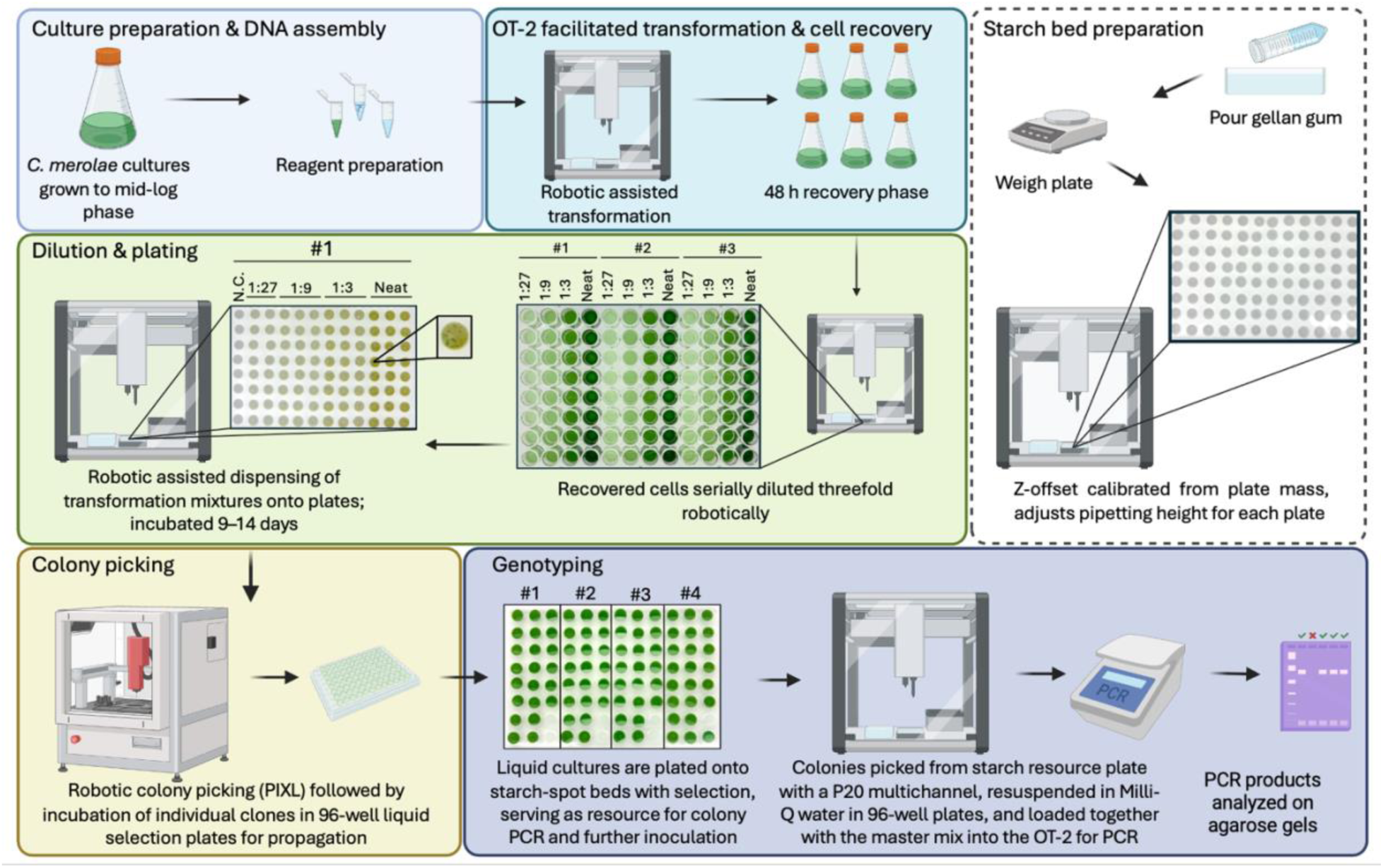
High-throughput automated transformation workflow for *Cyanidioschyzon merolae*. Overview of the fully automated transformation and screening pipeline integrating an OT-2 liquid-handling robot and a PIXL colony-picking system. Wild-type cultures are grown to mid-log phase, and DNA constructs are prepared for PEG-mediated transformation. The OT-2 performs transformation. After cell recovery, automated serial dilution and plating on starch-bed MA2G plates using Z-offset calibration based on gellan-gum weigh. After colony formation, the PIXL robot identifies and transfers individual colonies into 96-well plates for propagation. Colonies are subsequently arrayed in starch bed plates, ready for colony PCR setup on the OT-2.

For the development of starch-bed preparation and cell-plating routines, we adapted the strategy described in (Taguchi, 2023). Because gellan gum solidifies rapidly after preparation, precise volumetric dispensing of medium was not practical when preparing multiple plates in parallel. To address this limitation, plates were weighed after pouring, and the resulting gel height was calculated from the total plate mass using the density of gellan gum, applying the following formula:

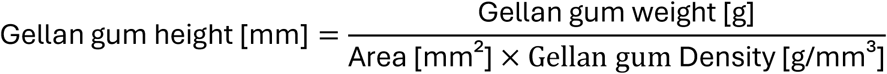

Each plate weight was then entered into a Python function within the Opentrons API, which calculates z-axis offset specific to that plate. This allowed automatic compensation for small variations in gel height between plates, positioning the pipette tip precisely 1 mm above the surface to accurately generate consistent starch spots and cell deposition.

Each automated transformation run was configured to execute multiple transformations in parallel, allowing simultaneous testing of different plasmid constructs or genomic loci under identical experimental conditions. Following transformation, the OT-2 was used to serially dilute the cells and, using the same z-axis correction logic, to plate the diluted cultures onto starch spots in an automated manner. Following transformant recovery, colonies are visually apparent as dark green spots on starch beds (Fig. 2). These can be isolated manually or using the colony picking robot system, and transferred to liquid medium in 96-well formats for downstream genotyping. This approach enables the entire transformation workflow to be standardized, from PEG incubation to colony isolation, minimizing manual intervention. To our knowledge, this constitutes the first robotics-assisted transformation platform developed for *C. merolae*.

### Benchmarking reporter expression across candidate genomic loci

To functionally evaluate the mapped loci, we employed the *mVenus* reporter plasmid library described above, in which each construct contains the homology arms for a specific candidate site across the genome. Integration was performed using the transformation pipeline described above. Transformants were plated on MA2G solid medium containing starch spots and recovered after ∼9 d of incubation at 42 °C under 5% CO₂. Colonies used for downstream analysis were first re-arrayed onto individual starch spots, and representative images of these plates capturing chlorophyll autofluorescence and mVenus fluorescence are shown in Fig. 3B and Supplementary Fig. 3. Successful integrations were confirmed by colony PCR using primers targeting the left and right junctions of each construct. Correct integration rates, approximately 94%, were comparable among all tested loci (Supplementary Table 5, Supplementary Fig. 5), indicating that non-homologous integration is rare and is not influenced by specific sequences at these loci. Of the 40 candidate loci, transformations were possible at 38 sites, as two loci (NL006 and NL011) could not be conclusively validated by genotyping (Supplementary Fig. 1C) or had challenges in DNA linearization by PCR. Prior to fluorescence quantification, flow cytometer parameters and gating were established by comparing wild-type cells with a representative *mVenus* transformant to define baseline fluorescence distributions (Fig. 3A). Quantification of mVenus fluorescence by flow cytometry across the 38 loci with confirmed integration revealed comparable expression (Fig. 3C and Supplementary Fig. 3). Although minor deviations could be seen in populations derived from each locus, this variation was not significant within the population of transformants analyzed. Welch’s ANOVA identified statistically significant variance between some NL, however, post hoc Tukey’s HSD testing showed that all transgenic loci (NL001–NL040) fell within overlapping significance groups (b–e) (Fig. 3C), indicating no statistically significant differences among them. The largest fold change among transformants was ∼1.4-fold, confirming that these minor numerical deviations are statistically detectable but not biologically meaningful. The results indicate that the selected genomic loci are effectively transcriptionally equivalent and do not impose positional bias on transgene expression.

**Figure 3.**
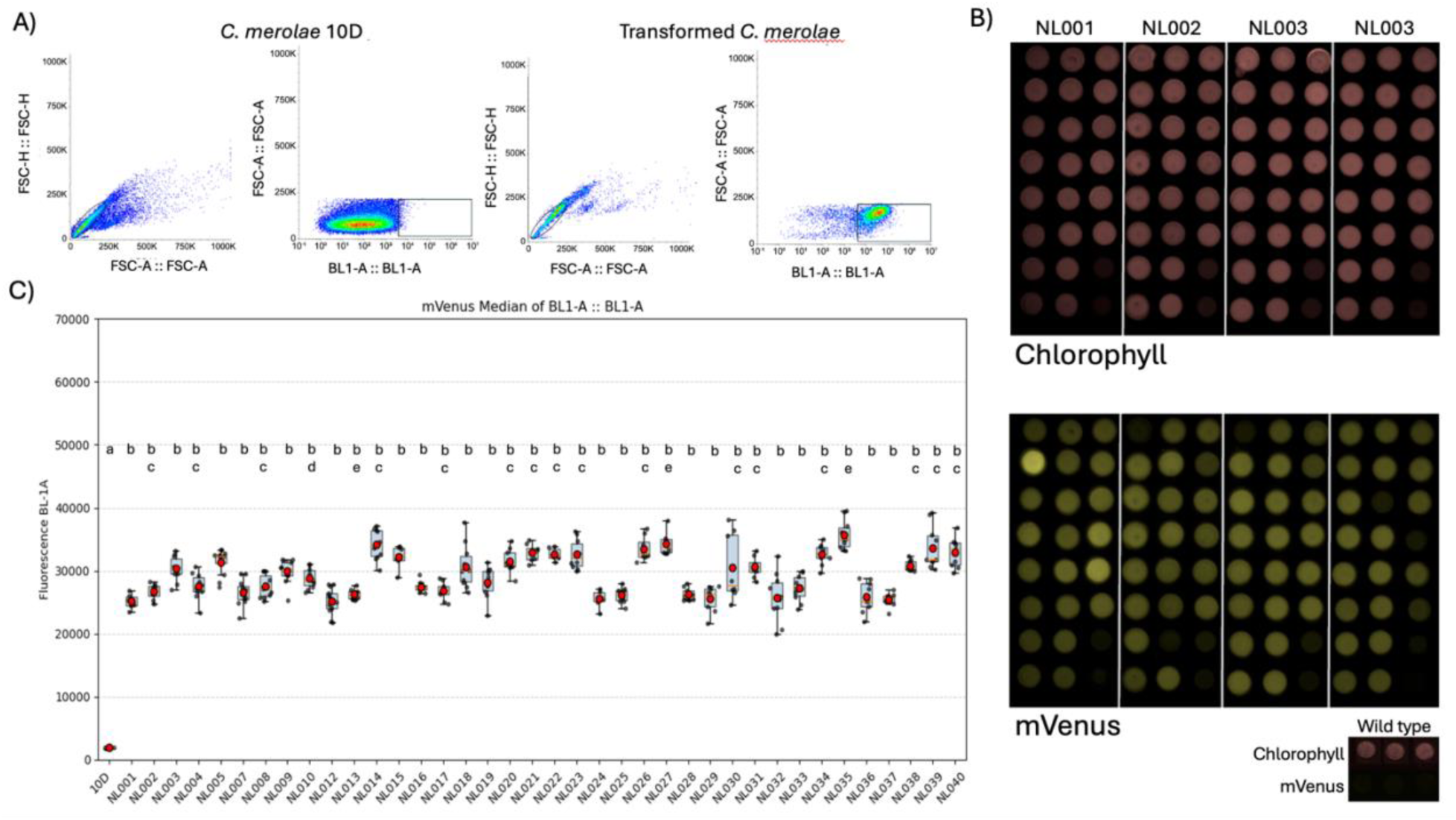
Reporter expression across validated genomic loci in *C. merolae*. (A) Flow cytometry profiles used to establish gating parameters for fluorescence quantification. Forward and side scatter distributions (FSC-H/FSC-A) used to exclude doublets and triplets are shown for wild-type (10D) and a representative *mVenus* transformant, followed by BL1-A fluorescence plots illustrating the fluorescence baseline of untransformed cells and the distinct fluorescence distribution of a positive transformant. (B) Representative plate images of colonies selected for downstream analysis. Each transformant was rearrayed onto individual starch spots and imaged for chlorophyll autofluorescence (top panels) and mVenus fluorescence (bottom panels), illustrating consistent reporter expression across biological replicates. (C) Flow-cytometry analysis of mVenus fluorescence from 10 independent transformants integrated at different loci. Each point represents a biological replicate; boxes show median ± SD in BL-1 channel. Letters above boxplots indicate statistically distinct groups based on one-way ANOVA followed by Tukey’s HSD post-hoc test (p < 0.05); identical letters denote no significant difference (p > 0.05).

The scarcity of repressive chromatin in *C. merolae* further supports its unusually permissive nuclear architecture. Silencing marks, such as H3K27me3, have been detected in only a small fraction of the genome, covering only 242 genes, and are largely confined to repetitive regions (Mikulski et al., 2017). This low abundance of epigenetic repression contrasts sharply with the extensive chromatin compartmentalization observed in metazoans. Consistent with this, the absence of strong hierarchy among genomic loci suggests that the *C. merolae* nuclear architecture tolerates exogenous insertions across multiple intergenic regions without major effects on gene expression or chromatin accessibility. This uniformity contrasts with patterns observed in more complex eukaryotes, where integration outcomes are often influenced by local chromatin state or epigenetic context (Policarpi et al., 2024; Zhang et al., 2020).

### Heterologous expression of an isoprene synthase in different loci

To evaluate the practical utility of the newly validated NL for metabolic engineering, we constructed a series of expression plasmids encoding a heterologous isoprene synthase (*IspS*) from sweet potato (*Ipomoaea batatas*) that was recently described to function in *C. merolae* (Villegas-Valencia et al., 2025). *C. merolae* does not naturally produce isoprene (Villegas-Valencia et al., 2025), however expression of the *Ib*IspS resulted in detectible isoprene levels that could be measured in its headspace (Villegas-Valencia et al., 2025; Yahya et al., 2025).The plasmids are identical in selection marker and gene of interest cassettes, varying only in target integration site in the genome. Six loci were randomly selected from NL sites described above (Fig. 4A). Transformant strains were generated as above and confirmed for both single copy insertion of the transgene and full-length expression of the *Ib*IspS-mVenus fusion by in-gel fluorescence (Supplementary Fig.4A & B).

**Figure 4.**
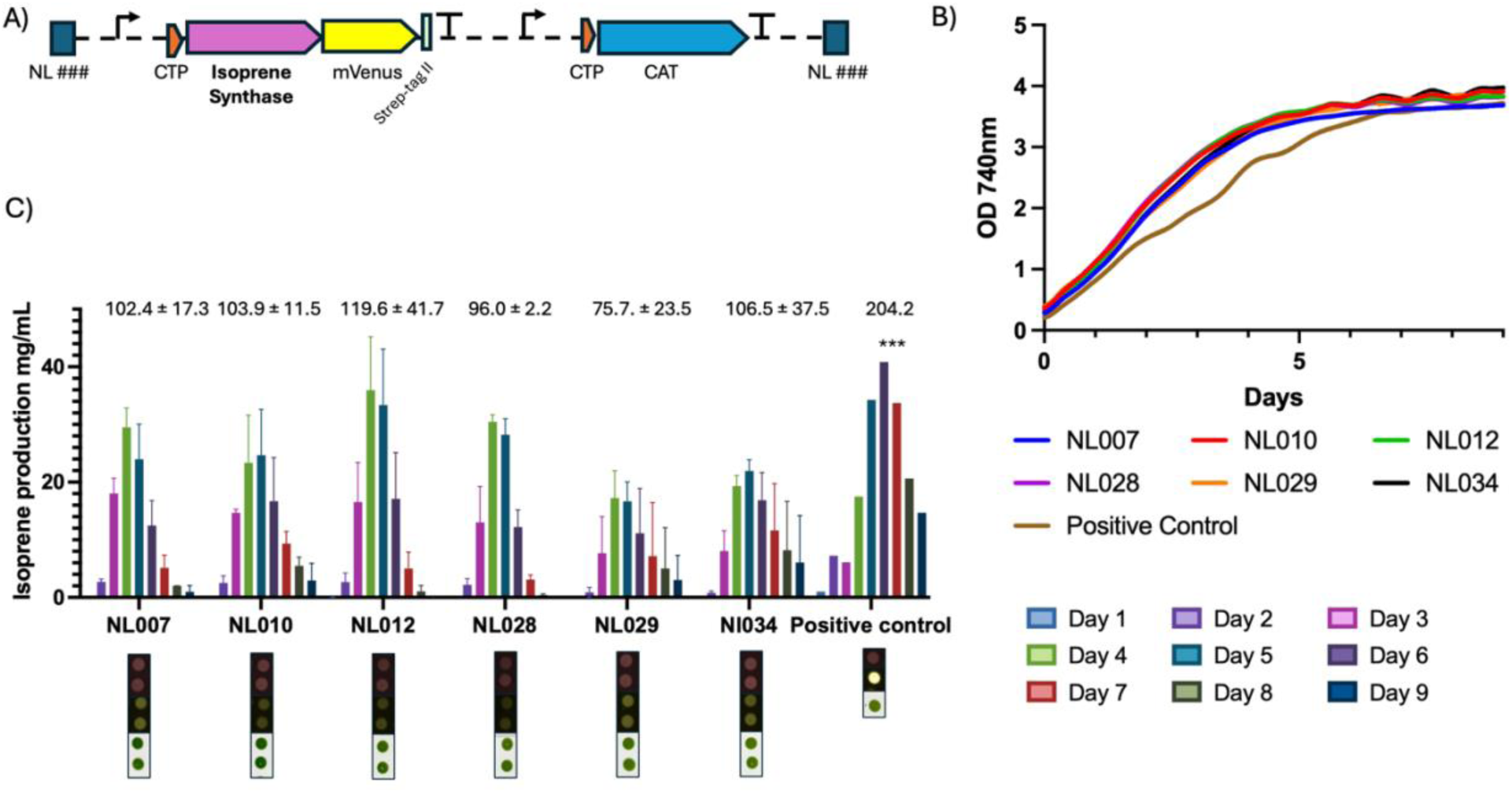
Benchmarking isoprene synthase expression across selected genomic loci in *C. merolae*. (A) Schematic of the isoprene-producing construct used for locus benchmarking. Each plasmid contains a left and right homology arm specific to the target locus, the *pCPCC* promoter driving a chloroplast-targeted isoprene synthase (CTP–Isoprene Synthase), followed by an *mVenus–Strep-tag II* reporter and *NOS* terminator. A chloramphenicol-resistance cassette (*CAT*) driven by the *pAPCC* promoter and terminated by the *3′ β-tubulin* terminator is included for selection. (B) Median optical density values from biological replicates of independently isolated transformants grown under identical cultivation conditions. (C) Isoprene production across six selected genomic loci compared to the positive control. Bar plots show daily isoprene titers (Days 1–9) as well as total accumulated isoprene for two independently isolated colonies per locus. Numerical values above the bars indicate total production whereas the positive-control strain produced significantly higher isoprene levels (***p < 0.001).

To measure isoprene production, transformed *C. merolae* strains were cultivated under standard phototrophic conditions, and headspace samples were continuously analyzed using gas chromatography–mass spectrometry (GC–MS) directly coupled to the algal culture vessel as recently described (Villegas-Valencia et al., 2025; Yahya et al., 2025). As expected, all six genomic integration sites supported isoprene synthesis which could be detected in the headspace. No statistically significant differences in isoprene production rates among them were observed and each strain exhibited similar growth behavior (Fig. 4B and C). A positive-control strain, developed with the CMD184-185C locus using a previously published plasmid was used (Villegas-Valencia et al., 2025). The generated strain was selected for high mVenus fluorescence and contained 6 copies of the *Ib*IspS-mVenus cassette. The positive control showed a marked increase in headspace isoprene levels (Fig. 4C and Supplementary Fig. 4), confirming that the analytical setup was sensitive and that increased abundance in synthase titer improves isoprene yields.

These results demonstrate that the NL are functionally interchangeable for metabolic engineering applications. The uniformity of isoprene production across independent genomic sites is consistent with the expression patterns observed for the mVenus reporter constructs, further validating the neutrality of the identified loci.

### Conclusions

This work establishes a practical and scalable framework for predictable nuclear genome engineering in *Cyanidioschyzon merolae*. By integrating bioinformatically-defined and experimentally-validated neutral loci with an automated transformation pipeline, we provide a standardized set of genomic landing pads that support uniform transgene expression and similar rates of locus-specific integration. The functional equivalence of these sites is demonstrated through consistent heterologous metabolite production across loci. The systematic identification and benchmarking of neutral sites provides a critical step for more complex engineering in *C. merolae*. These loci define predictable genomic coordinates that can be repeatedly targeted without confounding positional effects on expression. As a result, they enable stepwise genome rewriting strategies, including the distributed assembly of multi-gene pathways, iterative testing of regulatory architectures, and controlled expansion of engineered regions prior to chromosome-level consolidation. Importantly, benchmarking expression from defined genomic contexts establishes a reference framework against which emergent regulatory phenomena can be detected during chromosome reconstruction. Deviations from baseline behavior can thus be attributed to architectural features such as cryptic transcription, regulatory interference, or long-range chromosomal effects, rather than to unknown insertion-site variability. In this way, neutral-site mapping functions not merely as a convenience for transgene insertion, but as an essential measurement scaffold for incremental chromosome refactoring, analogous to approaches underpinning large-scale synthetic genome projects in other systems. In the era of synthetic genomics, the features determined here position *C. merolae* as a promising model to develop the first synthetic algal nuclear genome (Goold et al., 2024).

## Supporting information

Supplemental File 1 - All Plasmids

## Acknowledgements

This research was supported in part by the Intramural Research Program of the National Institutes of Health (NIH). The contributions of the NIH authors were made as part of their official duties as NIH federal employees, are in compliance with agency policy requirements, and are considered Works of the United States Government. However, the findings and conclusions presented in this paper are those of the authors and do not necessarily reflect the views of the NIH or the U.S. Department of Health and Human Services. SDR and MRS acknowledge funding from the Natural Sciences and Engineering Research Council of Canada Discovery Grant program and University of Northern British Columbia (UNBC)’s Office of Research and Innovation. KJL acknowledges baseline research funding from King Abdullah University of Science and Technology (KAUST) as well as funding from Office of Naval Research Global (ONRG) and United States Army Combat Capabilities Development Command (DEVCOM) Award N62909-25-1-2005. During the preparation of this manuscript, AI-assisted tools (ChatGPT version 4.o, OpenAI) were used to support the development of custom Python scripts for Opentrons OT-2 automation and figure generation. All code was reviewed, validated, and executed by the authors, who take full responsibility for the correctness, integrity, and interpretation of the results.

## Author Contributions

KT – Responsible for experimental work with *C. merolae* robotics, transformation and validation, supervision, method development, coding development, manuscript writing GBW – responsible for generating code and discovery of neutral loci, YZ – experimental workflow and data collection, BBF – experimental workflow for photobioreactors and supervision, VAS – bioinformatics analysis and curation of transcriptomic data, MRS, LZ, PJB, SDR – experimental planning, funding acquisition and workflow of transcriptomics data generation, KJL – funding acquisition, DNA construct design, supervision and resource acquisition.

## Figure Legends

**Supplementary Figure 1.** Genome-wide mapping and validation of candidate neutral integration sites in *Cyanidioschyzon merolae*. (A) Distribution of predicted “neutral” intergenic loci across all 20 nuclear chromosomes. Each plot shows the density of coding sequences (CDS; black line) along the chromosome and the position of selected neutral loci (colored vertical lines). Candidate sites were chosen to avoid gene-dense regions, repetitive sequences, and transcriptional readthrough. (B) Genomic context of the previously used CMD184C–CMD185C locus (orange), here designated NL029. RNA-seq coverage tracks on the plus (green) and minus (magenta) strands show detectable transcriptional readthrough across the intergenic region, indicating that this site does not meet neutrality criteria despite its widespread historical use in *C. merolae* transformation. (C) PCR-based validation of genomic integration for 40 candidate loci. Diagnostic amplification from genomic DNA confirmed correct insertion at 38 out of 39 mapped sites, producing the expected band size (∼1 kb) for each corresponding locus.

**Supplementary Figure 2:** Flow cytometry gating workflow used to define mVenus fluorescence in *C. merolae*. Representative gating strategy showing the sequential exclusion of debris (FSC-A vs FSC-H) and doublets or triplets (FSC-A vs BL1-A) prior to quantifying BL1-A fluorescence. The top panels show the non-transformed control (10D), used to establish the autofluorescence threshold. The same gates were applied to all transformants, illustrated by the representative mVenus-expressing strain NL002 (bottom panels).

**Supplementary Figure 3:** Colony-level fluorescence screening of *C. merolae* transformants. Following transformation, 22 colonies from each integration locus were re-plated onto starch beds and imaged for chlorophyll autofluorescence (red channel) and mVenus fluorescence (yellow channel). Each panel corresponds to an independent genomic locus (NL001–NL040). Chlorophyll signal indicates colony integrity and uniform growth, while mVenus fluorescence reflects successful transgene expression at the respective integration site.

**Supplementary Figure 4:** Quantitative PCR analysis of *mVenus* genomic copy number normalized to the 60S ribosomal subunit gene and full-length expression of the *IbIspS-mVenus* fusion validation. (A) Genomic DNA from strains carrying *mVenus* integrated at selected neutral loci (NL007, NL010, NL012, NL028, NL029, and NL034) was analyzed by qPCR. Bars represent mean ± SD of technical replicates (n = 2). *mVenus* copy number was normalized to the single-copy 60S ribosomal gene. (B) In-gel fluorescence analysis of protein extracts from the same strains, used to visualize mVenus and confirm its expected molecular weight across all integration sites.

