## Supplemental Figures 1-5 for "Genome Position Does Not Impact Transgene Expression Efficiency in the Ancient Red Alga *Cyanidioschyzon merolae*"

### Slide 1
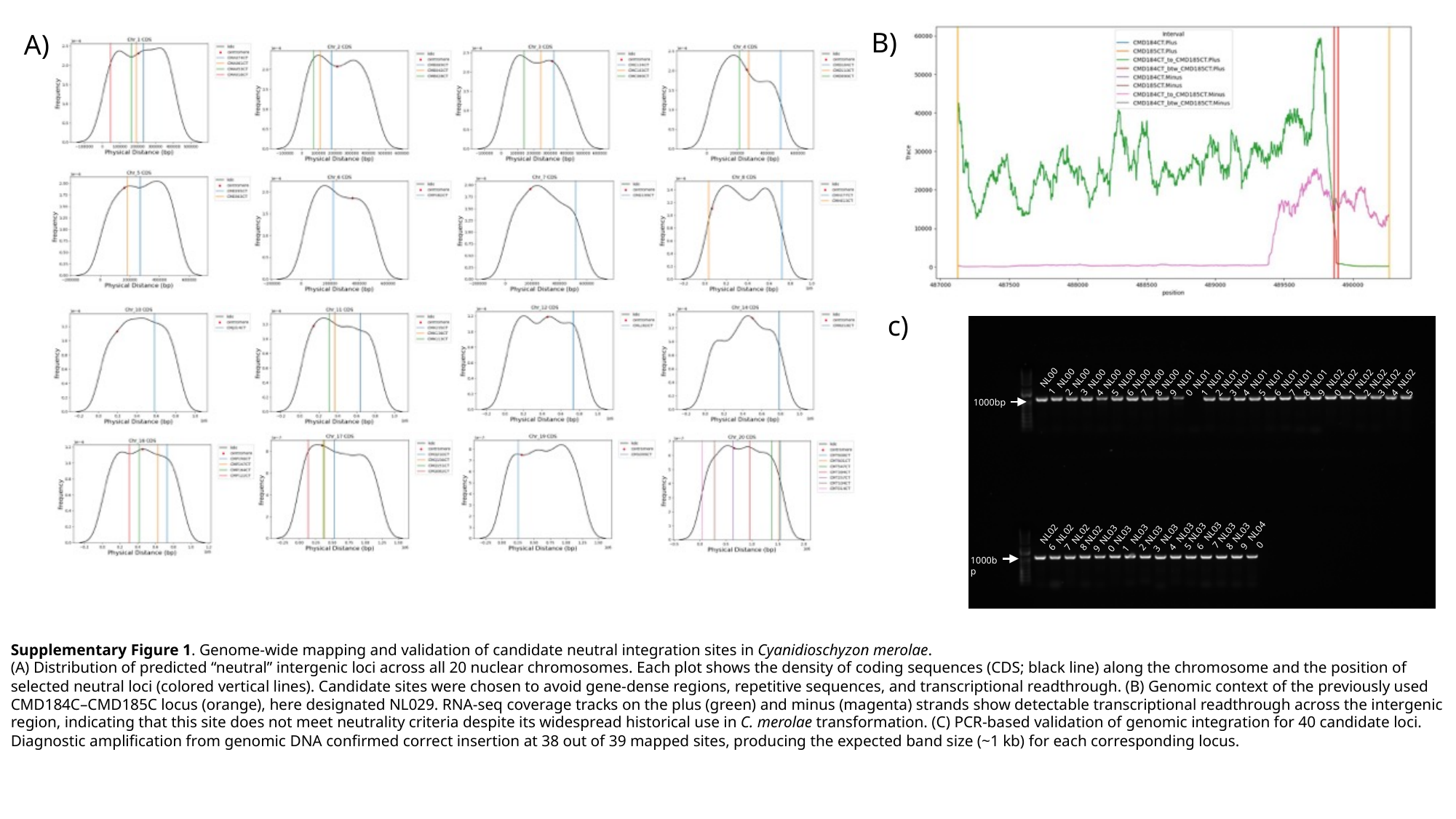

B)
A)
c)
NL001
NL002
NL003
NL025
NL024
NL020
NL022
NL021
NL023
NL015
NL016
NL017
NL018
NL019
NL008
NL009
NL010
NL012
NL011
NL004
NL006
NL007
NL013
NL014
NL005
1000bp
NL037
NL040
NL035
NL038
NL039
NL036
NL034
NL032
NL026
NL028
NL027
NL033
NL030
NL031
NL029
1000bp
Supplementary Figure 1. Genome-wide mapping and validation of candidate neutral integration sites in Cyanidioschyzon merolae.
(A) Distribution of predicted “neutral” intergenic loci across all 20 nuclear chromosomes. Each plot shows the density of coding sequences (CDS; black line) along the chromosome and the position of selected neutral loci (colored vertical lines). Candidate sites were chosen to avoid gene-dense regions, repetitive sequences, and transcriptional readthrough. (B) Genomic context of the previously used CMD184C–CMD185C locus (orange), here designated NL029. RNA-seq coverage tracks on the plus (green) and minus (magenta) strands show detectable transcriptional readthrough across the intergenic region, indicating that this site does not meet neutrality criteria despite its widespread historical use in C. merolae transformation. (C) PCR-based validation of genomic integration for 40 candidate loci. Diagnostic amplification from genomic DNA confirmed correct insertion at 38 out of 39 mapped sites, producing the expected band size (~1 kb) for each corresponding locus.

### Slide 2
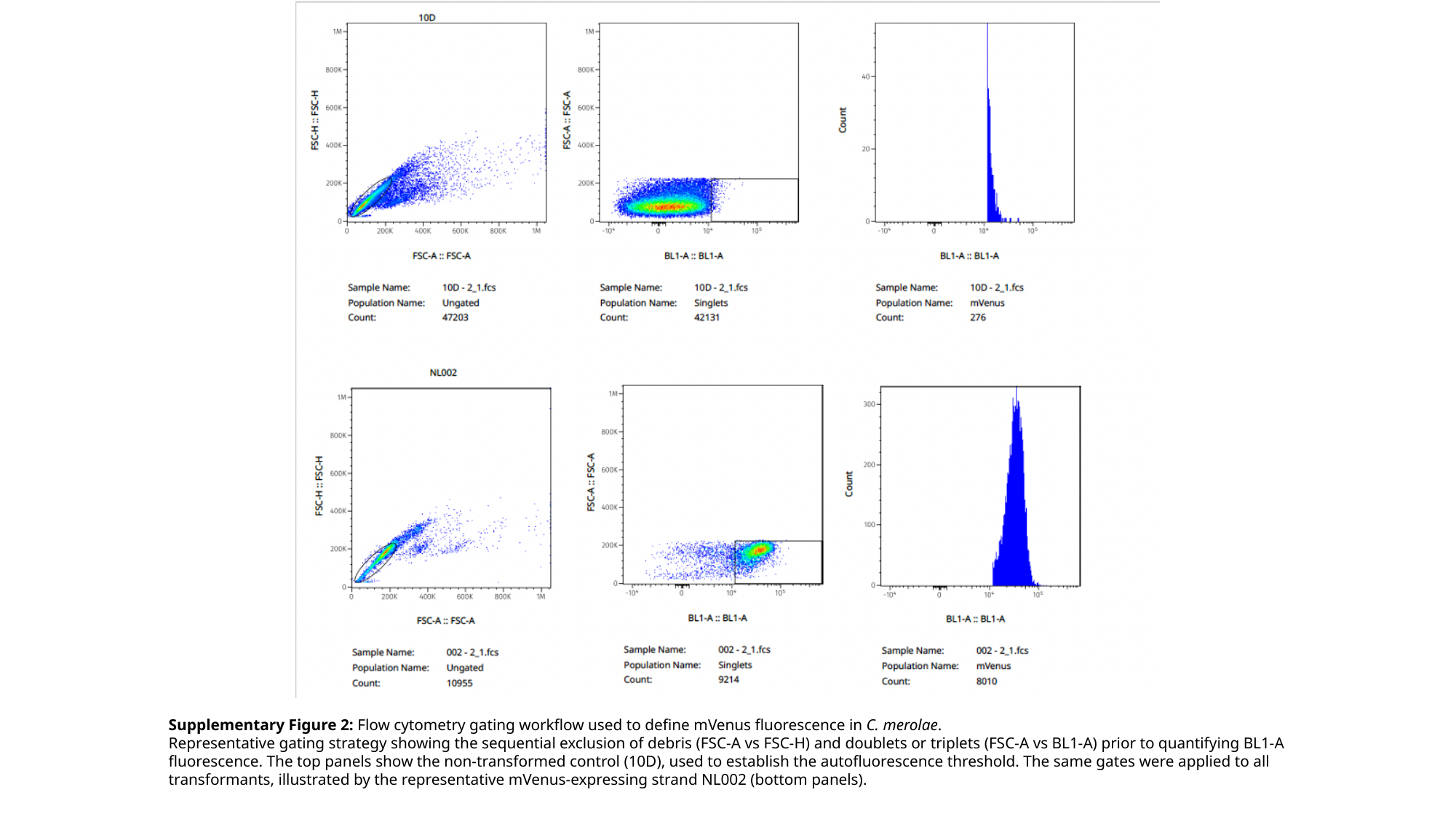

Supplementary Figure 2: Flow cytometry gating workflow used to define mVenus fluorescence in C. merolae.
Representative gating strategy showing the sequential exclusion of debris (FSC-A vs FSC-H) and doublets or triplets (FSC-A vs BL1-A) prior to quantifying BL1-A fluorescence. The top panels show the non-transformed control (10D), used to establish the autofluorescence threshold. The same gates were applied to all transformants, illustrated by the representative mVenus-expressing strand NL002 (bottom panels).

### Slide 3
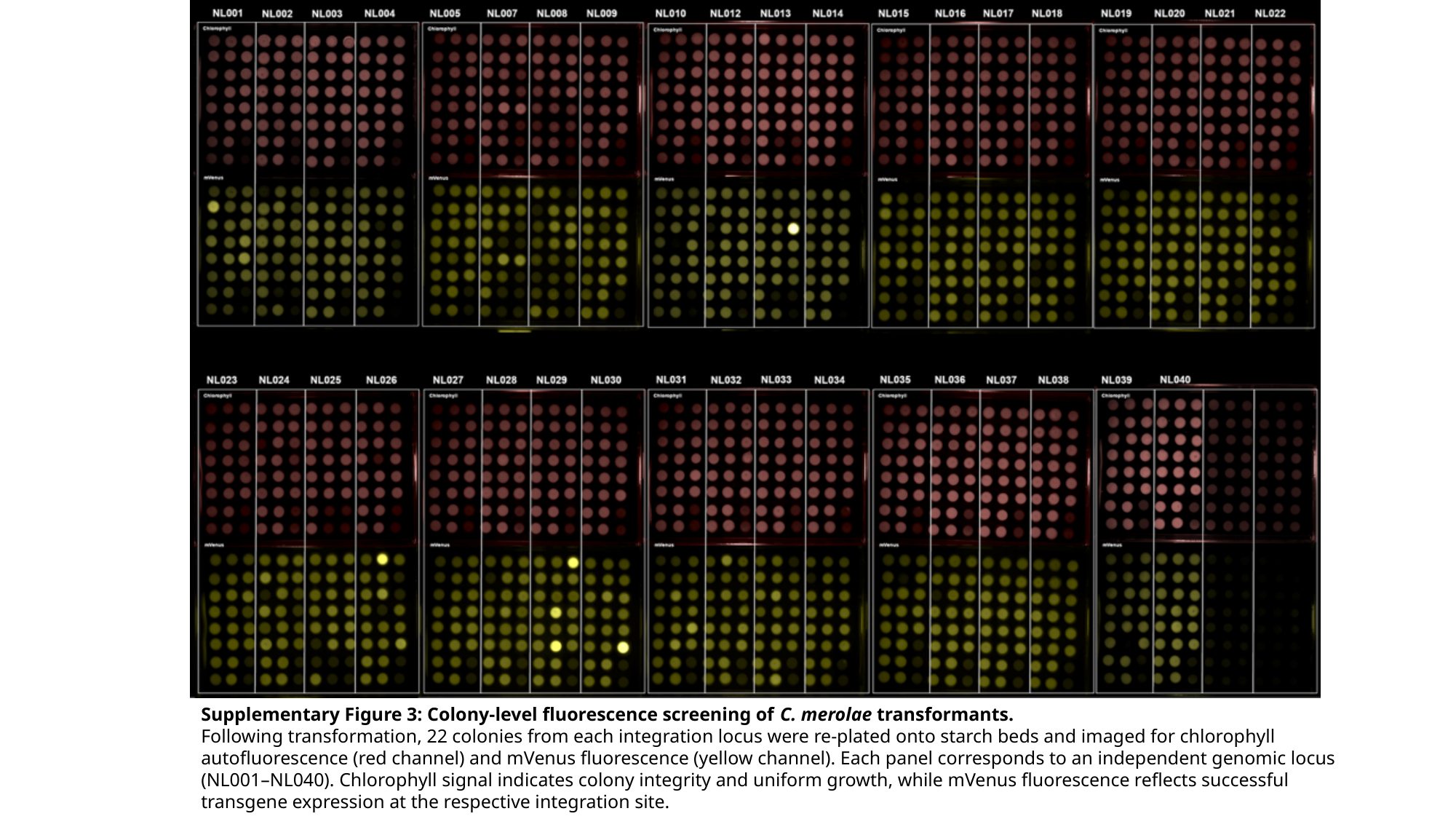

Supplementary Figure 3: Colony-level fluorescence screening of C. merolae transformants.
Following transformation, 22 colonies from each integration locus were re-plated onto starch beds and imaged for chlorophyll autofluorescence (red channel) and mVenus fluorescence (yellow channel). Each panel corresponds to an independent genomic locus (NL001–NL040). Chlorophyll signal indicates colony integrity and uniform growth, while mVenus fluorescence reflects successful transgene expression at the respective integration site.

### Slide 4
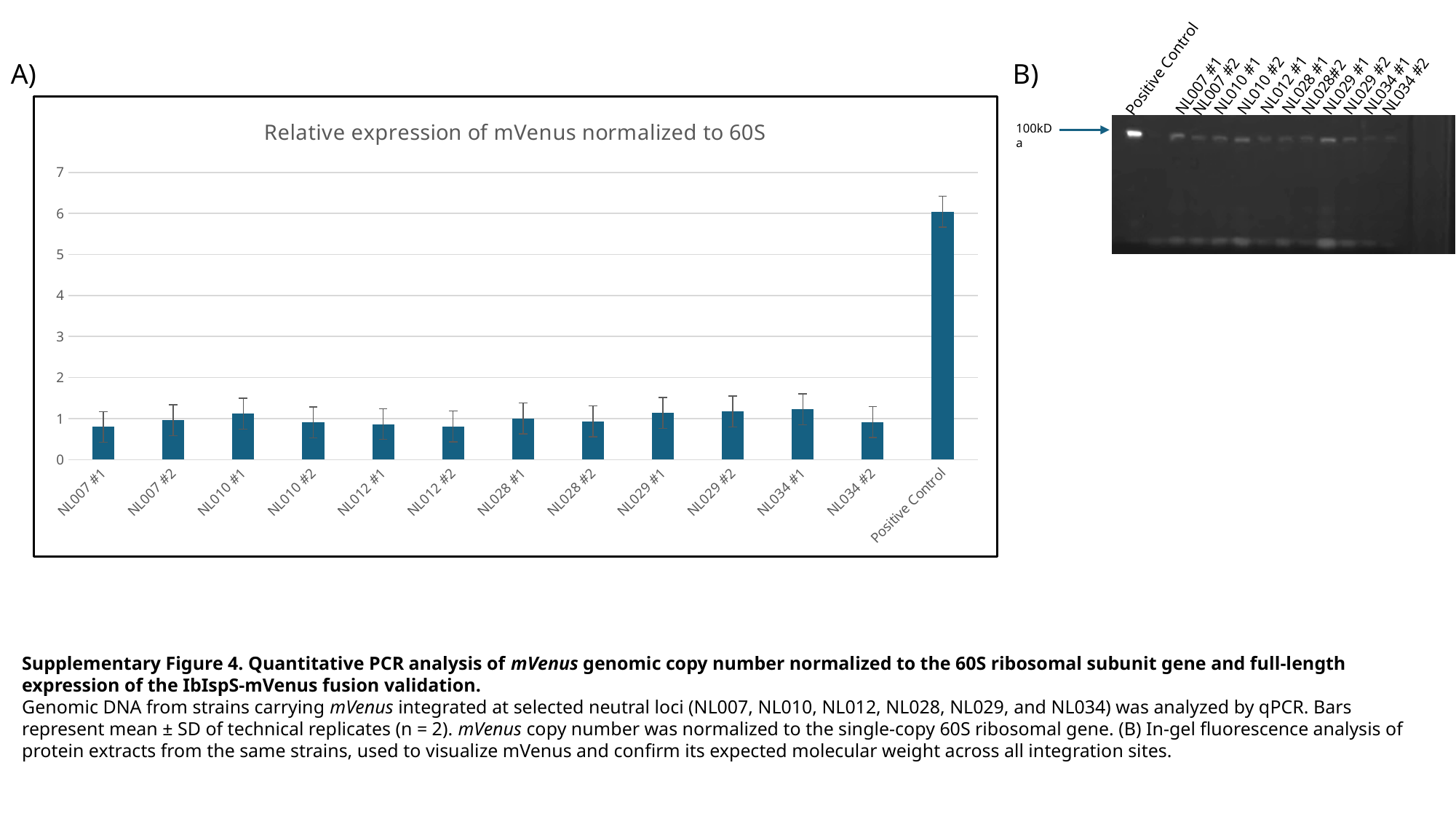

NL012 #1
NL028 #1
NL029 #2
NL034 #1
NL029 #1
NL028#2
NL010 #1
NL010 #2
NL007 #1
Positive Control
NL034 #2
NL007 #2
A)
B)
###### Chart: Relative expression of mVenus normalized to 60S
| Category | |
|---|---|
| NL007 #1 | 0.7963124491629298 |
| NL007 #2 | 0.9589140912959782 |
| NL010 #1 | 1.1175722997952011 |
| NL010 #2 | 0.9047138870333029 |
| NL012 #1 | 0.8639485908960043 |
| NL012 #2 | 0.8080629946482307 |
| NL028 #1 | 1.004119646608893 |
| NL028 #2 | 0.9337426175176387 |
| NL029 #1 | 1.1379594723317707 |
| NL029 #2 | 1.174087612851584 |
| NL034 #1 | 1.2236410593669944 |
| NL034 #2 | 0.9150230726366967 |
| Positive Control | 6.042865133298771 |100kDa
Supplementary Figure 4. Quantitative PCR analysis of mVenus genomic copy number normalized to the 60S ribosomal subunit gene and full-length expression of the IbIspS-mVenus fusion validation.
Genomic DNA from strains carrying mVenus integrated at selected neutral loci (NL007, NL010, NL012, NL028, NL029, and NL034) was analyzed by qPCR. Bars represent mean ± SD of technical replicates (n = 2). mVenus copy number was normalized to the single-copy 60S ribosomal gene. (B) In-gel fluorescence analysis of protein extracts from the same strains, used to visualize mVenus and confirm its expected molecular weight across all integration sites.

### Slide 5
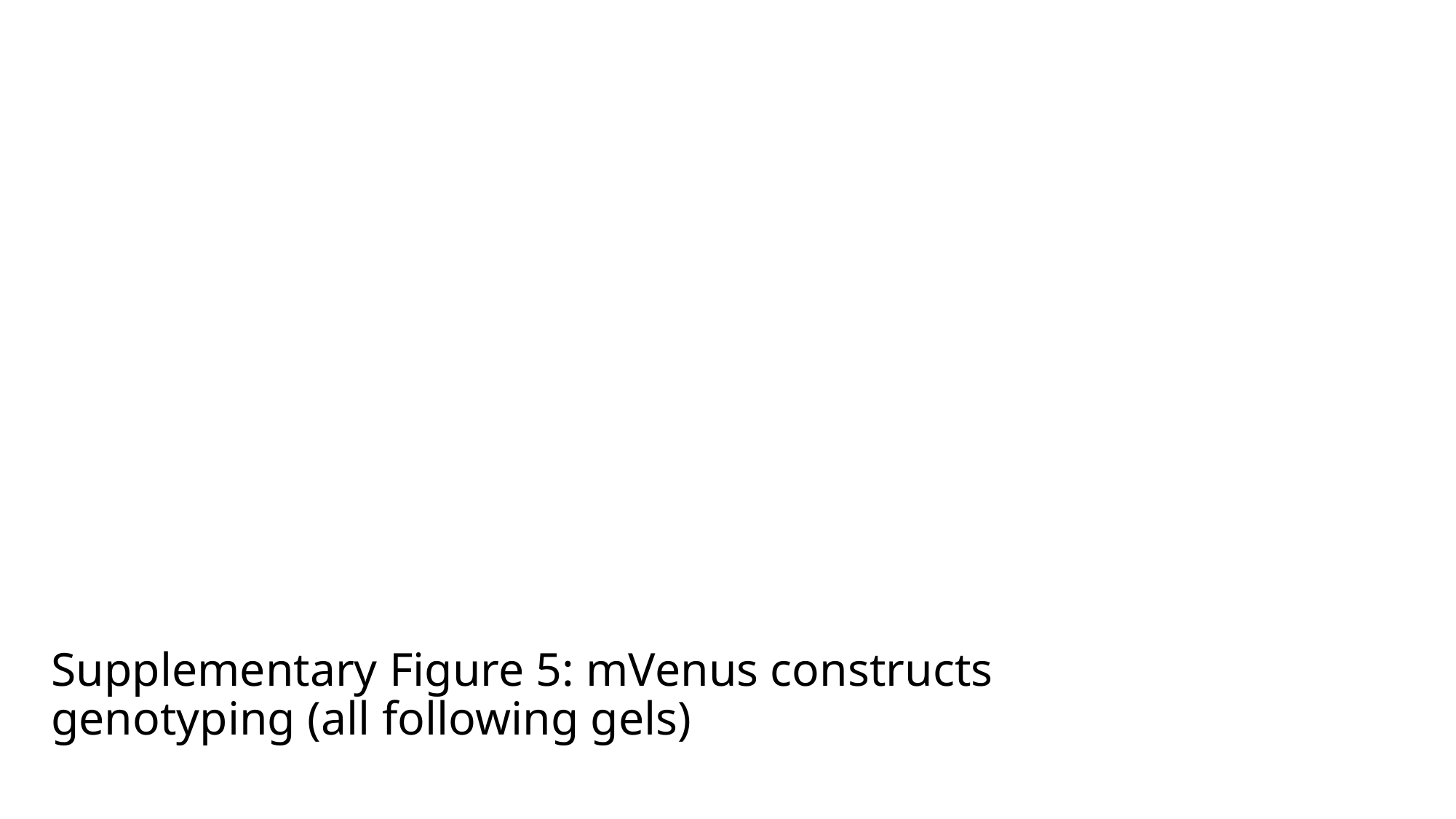

#### Supplementary Figure 5: mVenus constructs genotyping (all following gels)

### Slide 6
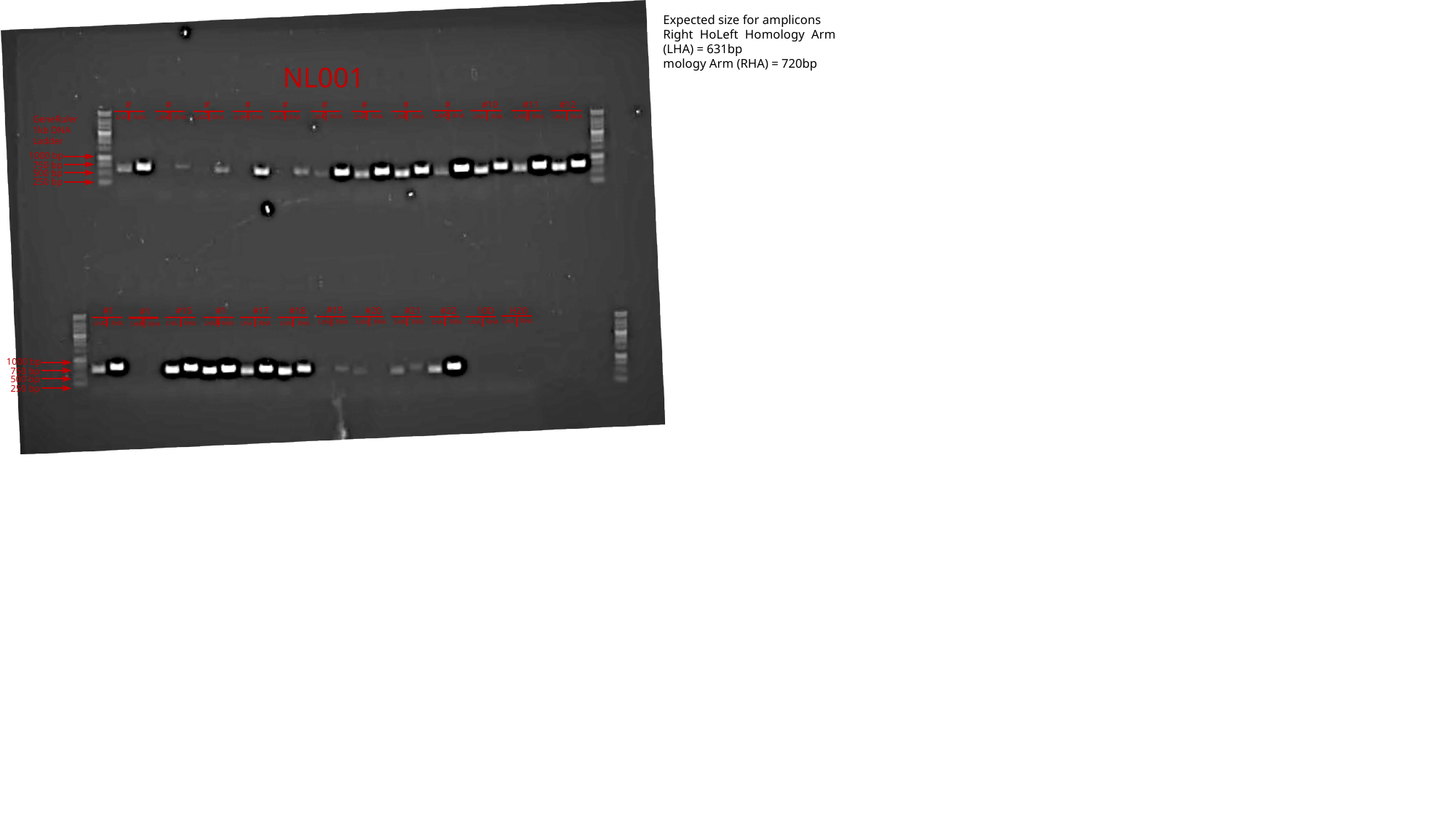

Expected size for amplicons
Right HoLeft Homology Arm (LHA) = 631bp
mology Arm (RHA) = 720bp
NL001
#3
#4
#5
#6
#7
#8
#9
#10
#11
#12
#1
#2
LHA RHA
LHA RHA
LHA RHA
LHA RHA
LHA RHA
LHA RHA
GeneRuler 1kb DNA Ladder
LHA RHA
LHA RHA
LHA RHA
LHA RHA
LHA RHA
LHA RHA
1000 bp
750 bp
500 bp
250 bp
#19
#20
#21
#22
10D
H20
#18
#16
#17
#13
#15
#14
LHA RHA
LHA RHA
LHA RHA
LHA RHA
LHA RHA
LHA RHA
LHA RHA
LHA RHA
LHA RHA
LHA RHA
LHA RHA
LHA RHA
1000 bp
750 bp
500 bp
250 bp

### Slide 7
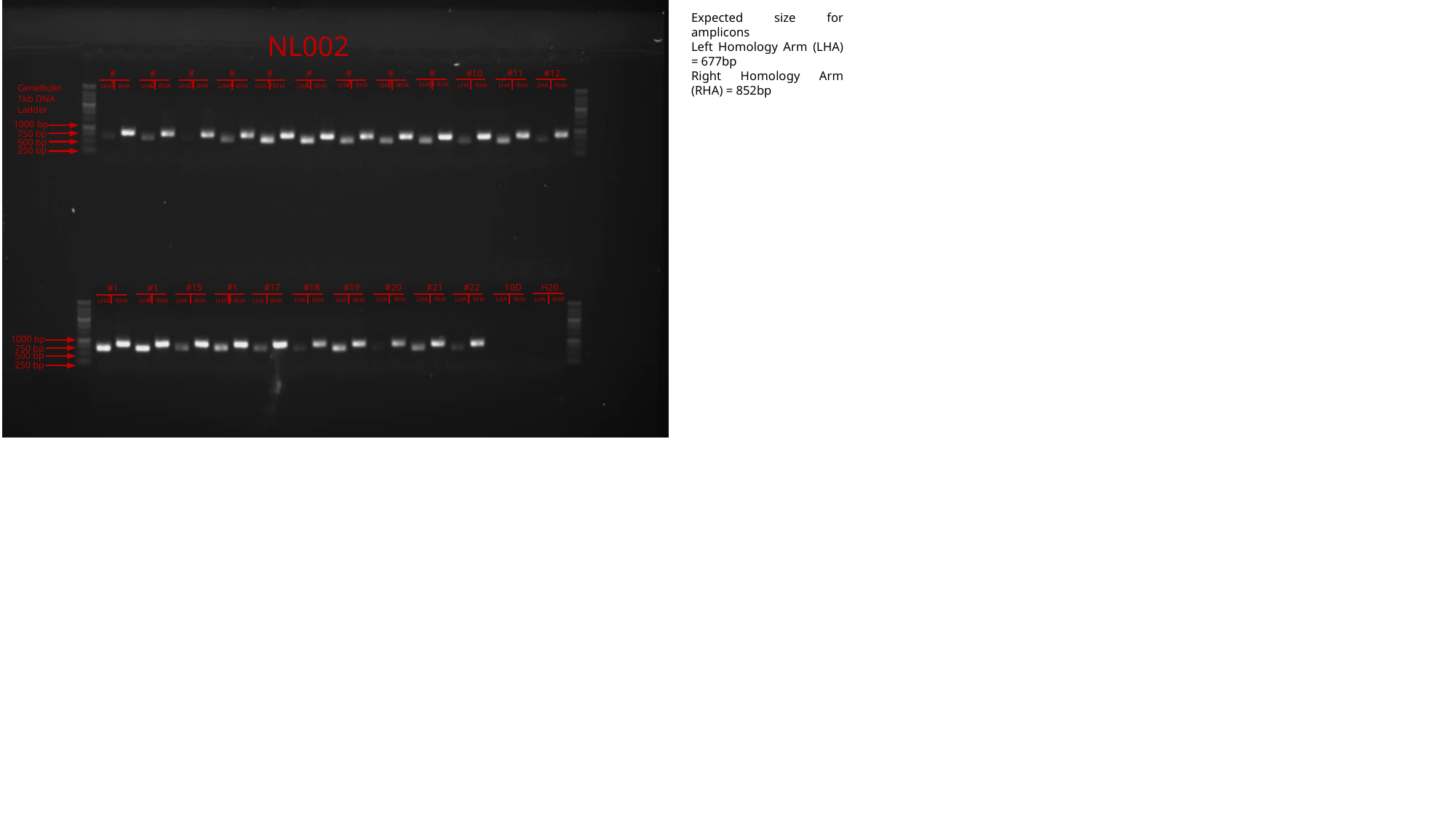

Expected size for amplicons
Left Homology Arm (LHA) = 677bp
Right Homology Arm (RHA) = 852bp
NL002
#3
#4
#5
#6
#7
#8
#9
#10
#11
#12
#1
#2
LHA RHA
LHA RHA
LHA RHA
LHA RHA
LHA RHA
LHA RHA
GeneRuler 1kb DNA Ladder
LHA RHA
LHA RHA
LHA RHA
LHA RHA
LHA RHA
LHA RHA
1000 bp
750 bp
500 bp
250 bp
#15
#16
#17
#18
#19
#20
#21
#22
10D
H20
#13
#14
LHA RHA
LHA RHA
LHA RHA
LHA RHA
LHA RHA
LHA RHA
LHA RHA
LHA RHA
LHA RHA
LHA RHA
LHA RHA
LHA RHA
1000 bp
750 bp
500 bp
250 bp

### Slide 8
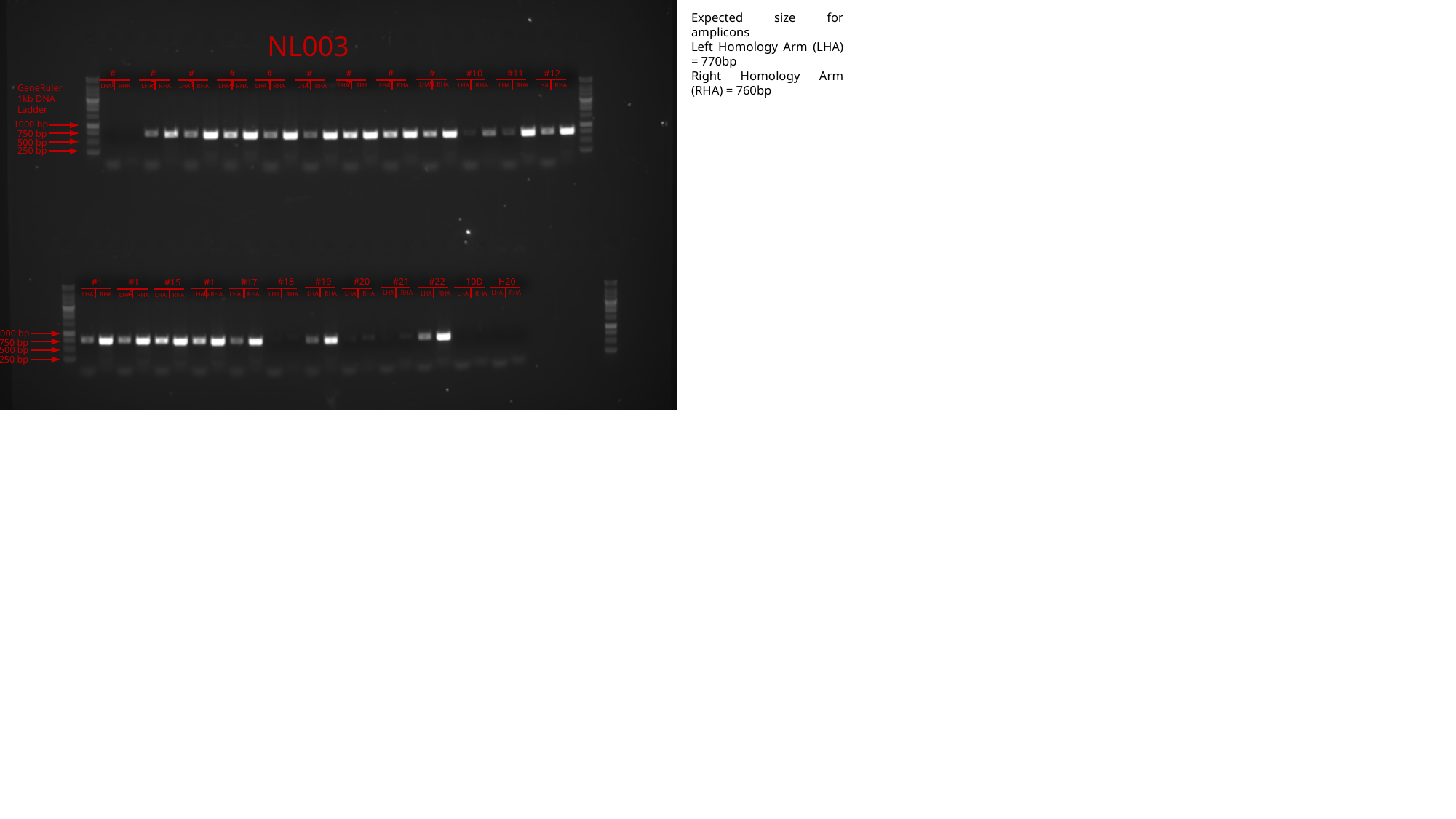

Expected size for amplicons
Left Homology Arm (LHA) = 770bp
Right Homology Arm (RHA) = 760bp
NL003
#3
#4
#5
#6
#7
#8
#9
#10
#11
#12
#1
#2
LHA RHA
LHA RHA
LHA RHA
LHA RHA
LHA RHA
LHA RHA
GeneRuler 1kb DNA Ladder
LHA RHA
LHA RHA
LHA RHA
LHA RHA
LHA RHA
LHA RHA
1000 bp
750 bp
500 bp
250 bp
#19
#20
#21
#22
10D
H20
#18
#16
#17
#13
#15
#14
LHA RHA
LHA RHA
LHA RHA
LHA RHA
LHA RHA
LHA RHA
LHA RHA
LHA RHA
LHA RHA
LHA RHA
LHA RHA
LHA RHA
1000 bp
750 bp
500 bp
250 bp

### Slide 9
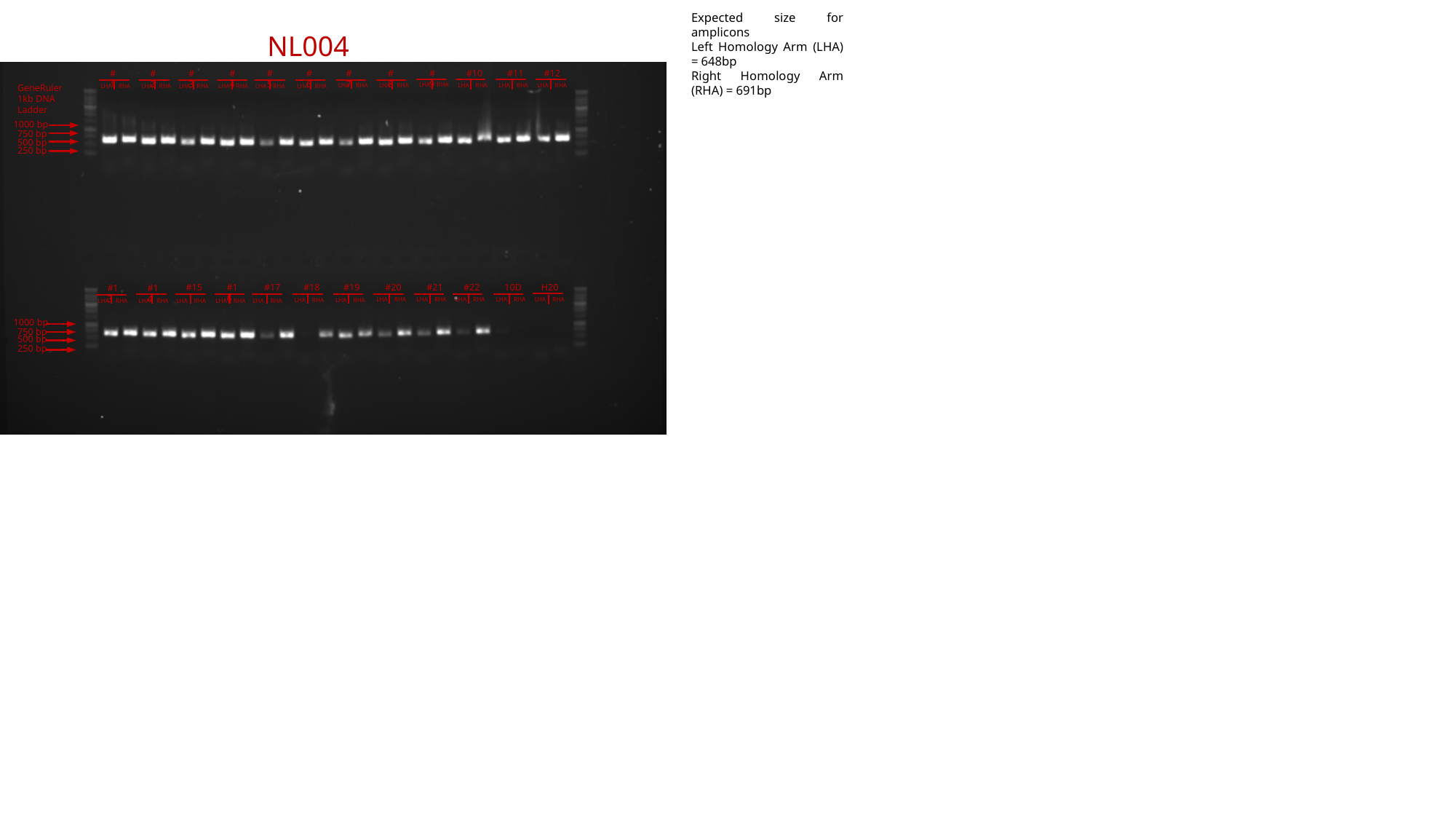

Expected size for amplicons
Left Homology Arm (LHA) = 648bp
Right Homology Arm (RHA) = 691bp
NL004
#3
#4
#5
#6
#7
#8
#9
#10
#11
#12
#1
#2
LHA RHA
LHA RHA
LHA RHA
LHA RHA
LHA RHA
LHA RHA
GeneRuler 1kb DNA Ladder
LHA RHA
LHA RHA
LHA RHA
LHA RHA
LHA RHA
LHA RHA
1000 bp
750 bp
500 bp
250 bp
#15
#16
#17
#18
#19
#20
#21
#22
10D
H20
#13
#14
LHA RHA
LHA RHA
LHA RHA
LHA RHA
LHA RHA
LHA RHA
LHA RHA
LHA RHA
LHA RHA
LHA RHA
LHA RHA
LHA RHA
1000 bp
750 bp
500 bp
250 bp

### Slide 10
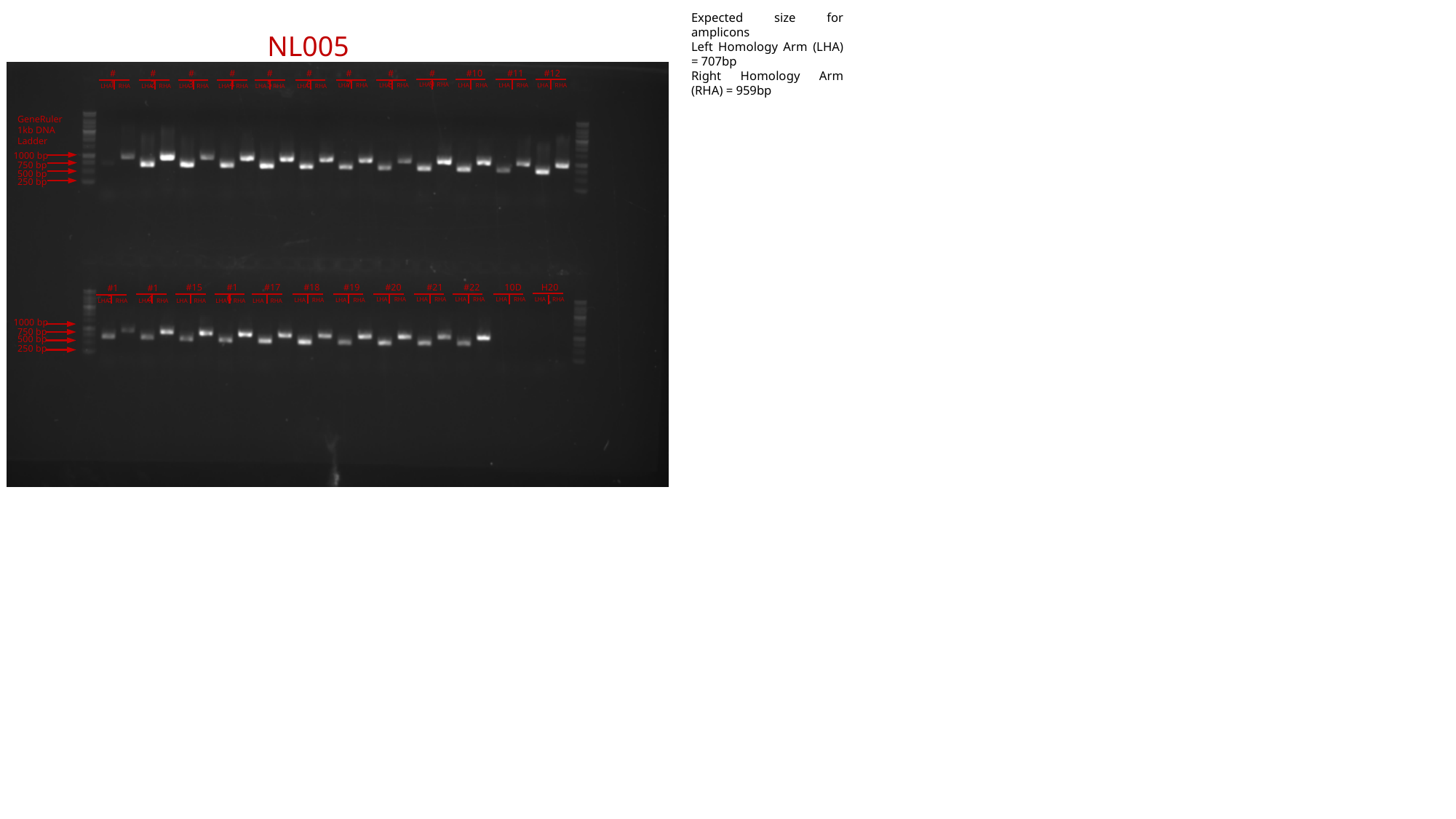

Expected size for amplicons
Left Homology Arm (LHA) = 707bp
Right Homology Arm (RHA) = 959bp
NL005
#3
#4
#5
#6
#7
#8
#9
#10
#11
#12
#1
#2
LHA RHA
LHA RHA
LHA RHA
LHA RHA
LHA RHA
LHA RHA
LHA RHA
LHA RHA
LHA RHA
LHA RHA
LHA RHA
LHA RHA
GeneRuler 1kb DNA Ladder
1000 bp
750 bp
500 bp
250 bp
#15
#16
#17
#18
#19
#20
#21
#22
10D
H20
#13
#14
LHA RHA
LHA RHA
LHA RHA
LHA RHA
LHA RHA
LHA RHA
LHA RHA
LHA RHA
LHA RHA
LHA RHA
LHA RHA
LHA RHA
1000 bp
750 bp
500 bp
250 bp

### Slide 11
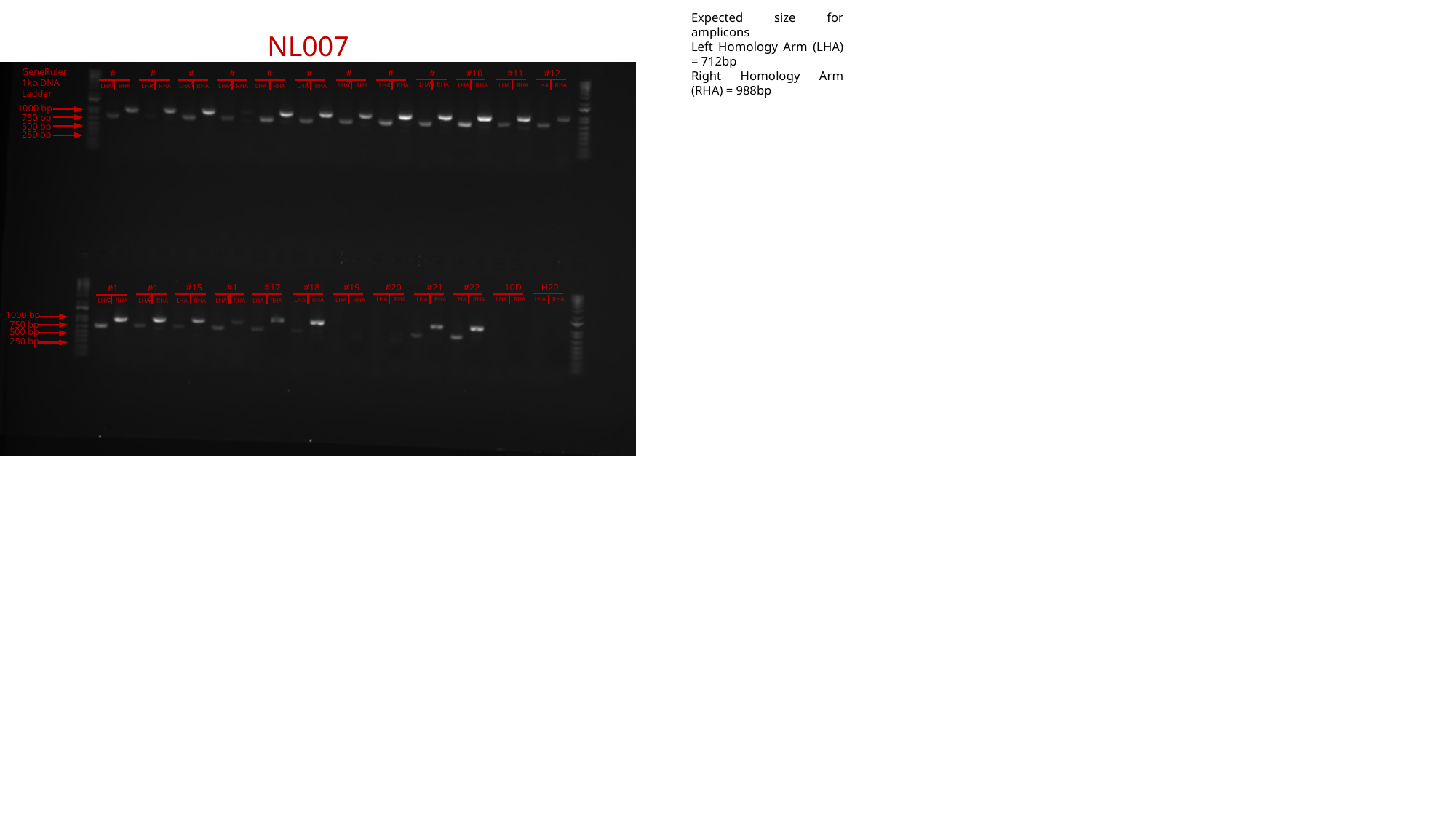

Expected size for amplicons
Left Homology Arm (LHA) = 712bp
Right Homology Arm (RHA) = 988bp
NL007
GeneRuler 1kb DNA Ladder
#3
#4
#5
#6
#7
#8
#9
#10
#11
#12
#1
#2
LHA RHA
LHA RHA
LHA RHA
LHA RHA
LHA RHA
LHA RHA
LHA RHA
LHA RHA
LHA RHA
LHA RHA
LHA RHA
LHA RHA
1000 bp
750 bp
500 bp
250 bp
#15
#16
#17
#18
#19
#20
#21
#22
10D
H20
#13
#14
LHA RHA
LHA RHA
LHA RHA
LHA RHA
LHA RHA
LHA RHA
LHA RHA
LHA RHA
LHA RHA
LHA RHA
LHA RHA
LHA RHA
1000 bp
750 bp
500 bp
250 bp

### Slide 12
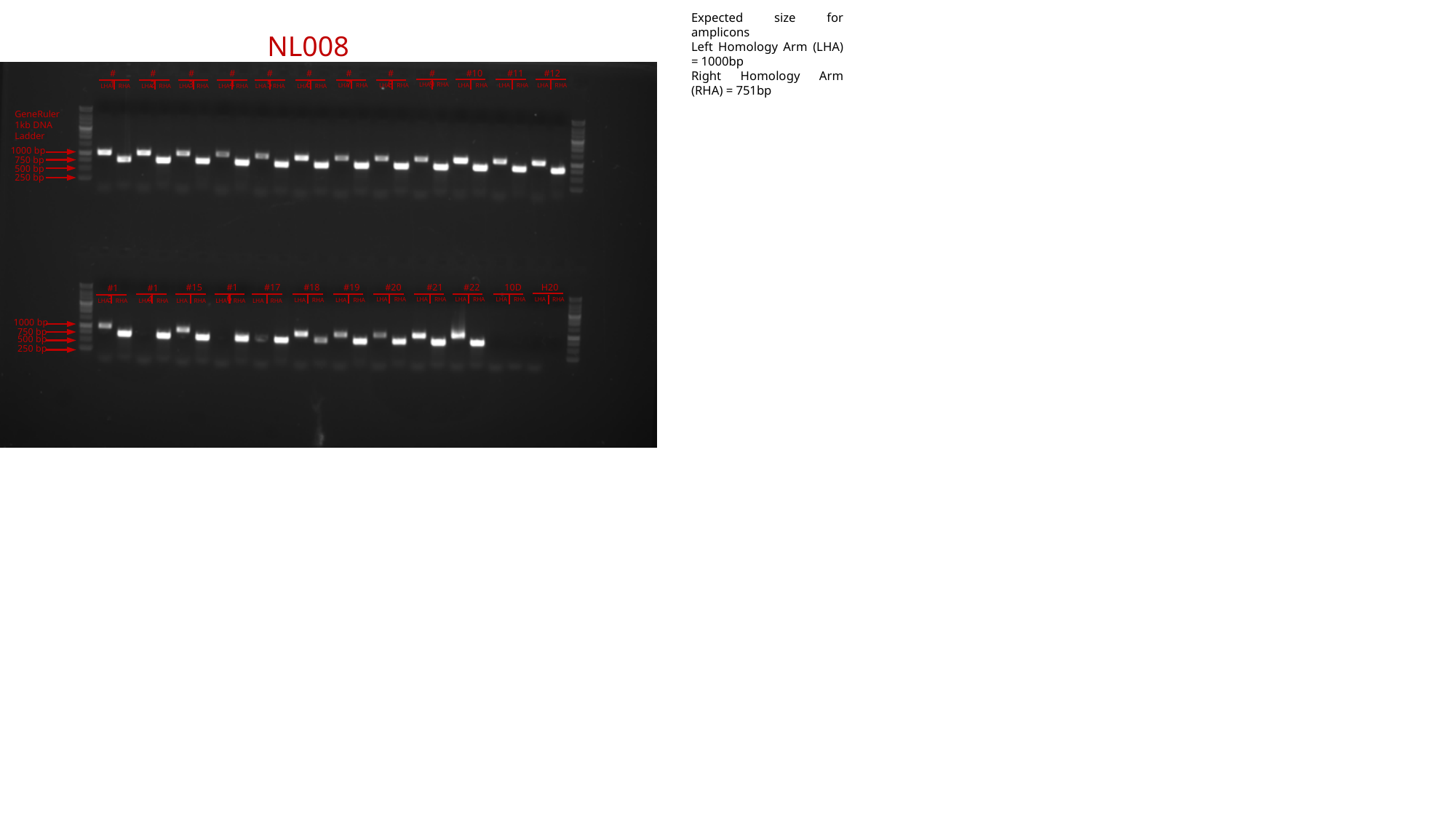

Expected size for amplicons
Left Homology Arm (LHA) = 1000bp
Right Homology Arm (RHA) = 751bp
NL008
#3
#4
#5
#6
#7
#8
#9
#10
#11
#12
#1
#2
LHA RHA
LHA RHA
LHA RHA
LHA RHA
LHA RHA
LHA RHA
LHA RHA
LHA RHA
LHA RHA
LHA RHA
LHA RHA
LHA RHA
GeneRuler 1kb DNA Ladder
1000 bp
750 bp
500 bp
250 bp
#15
#16
#17
#18
#19
#20
#21
#22
10D
H20
#13
#14
LHA RHA
LHA RHA
LHA RHA
LHA RHA
LHA RHA
LHA RHA
LHA RHA
LHA RHA
LHA RHA
LHA RHA
LHA RHA
LHA RHA
1000 bp
750 bp
500 bp
250 bp

### Slide 13
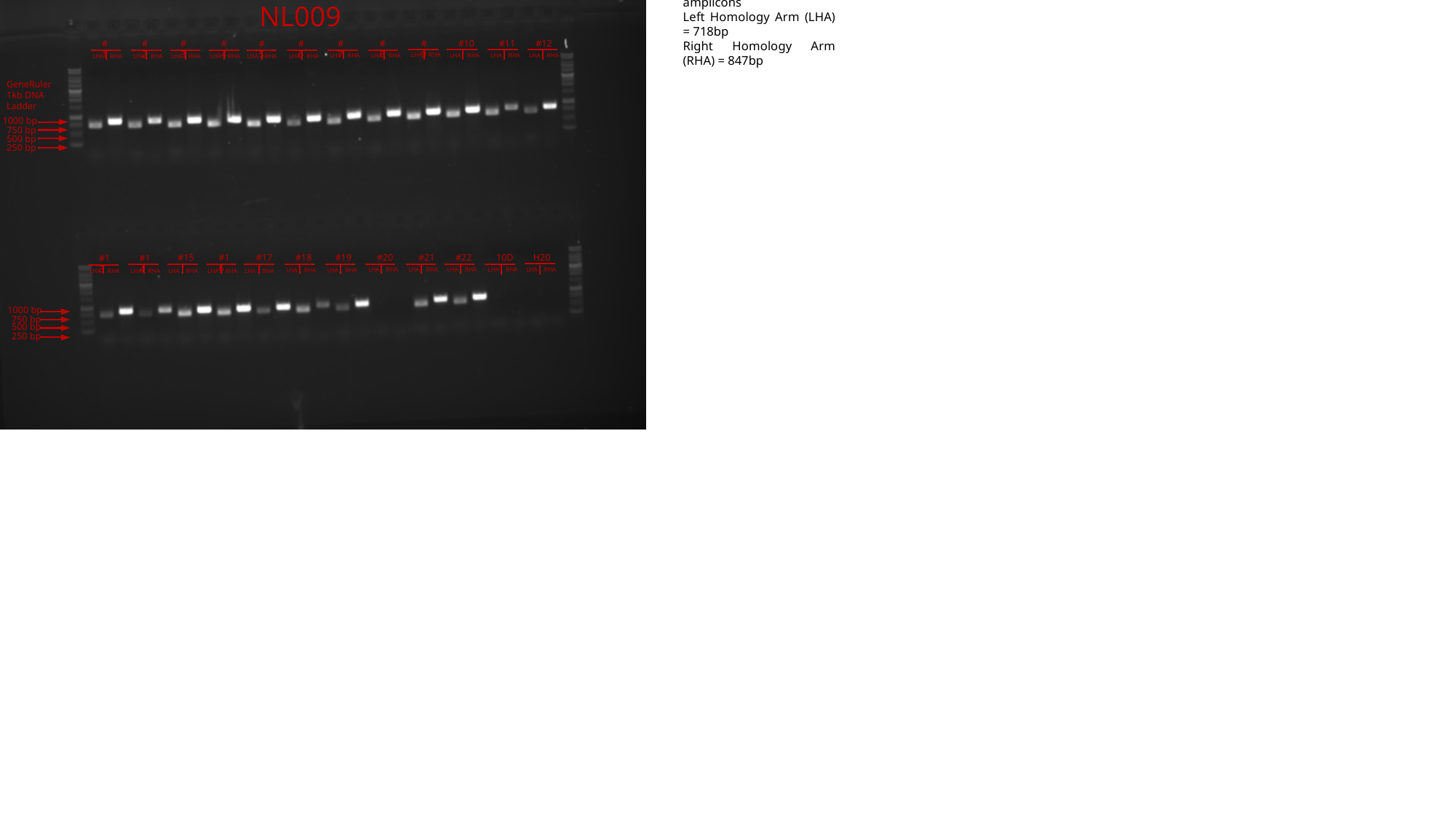

Expected size for amplicons
Left Homology Arm (LHA) = 718bp
Right Homology Arm (RHA) = 847bp
NL009
#3
#4
#5
#6
#7
#8
#9
#10
#11
#12
#1
#2
LHA RHA
LHA RHA
LHA RHA
LHA RHA
LHA RHA
LHA RHA
LHA RHA
LHA RHA
LHA RHA
LHA RHA
LHA RHA
LHA RHA
GeneRuler 1kb DNA Ladder
1000 bp
750 bp
500 bp
250 bp
#15
#16
#17
#18
#19
#20
#21
#22
10D
H20
#13
#14
LHA RHA
LHA RHA
LHA RHA
LHA RHA
LHA RHA
LHA RHA
LHA RHA
LHA RHA
LHA RHA
LHA RHA
LHA RHA
LHA RHA
1000 bp
750 bp
500 bp
250 bp

### Slide 14
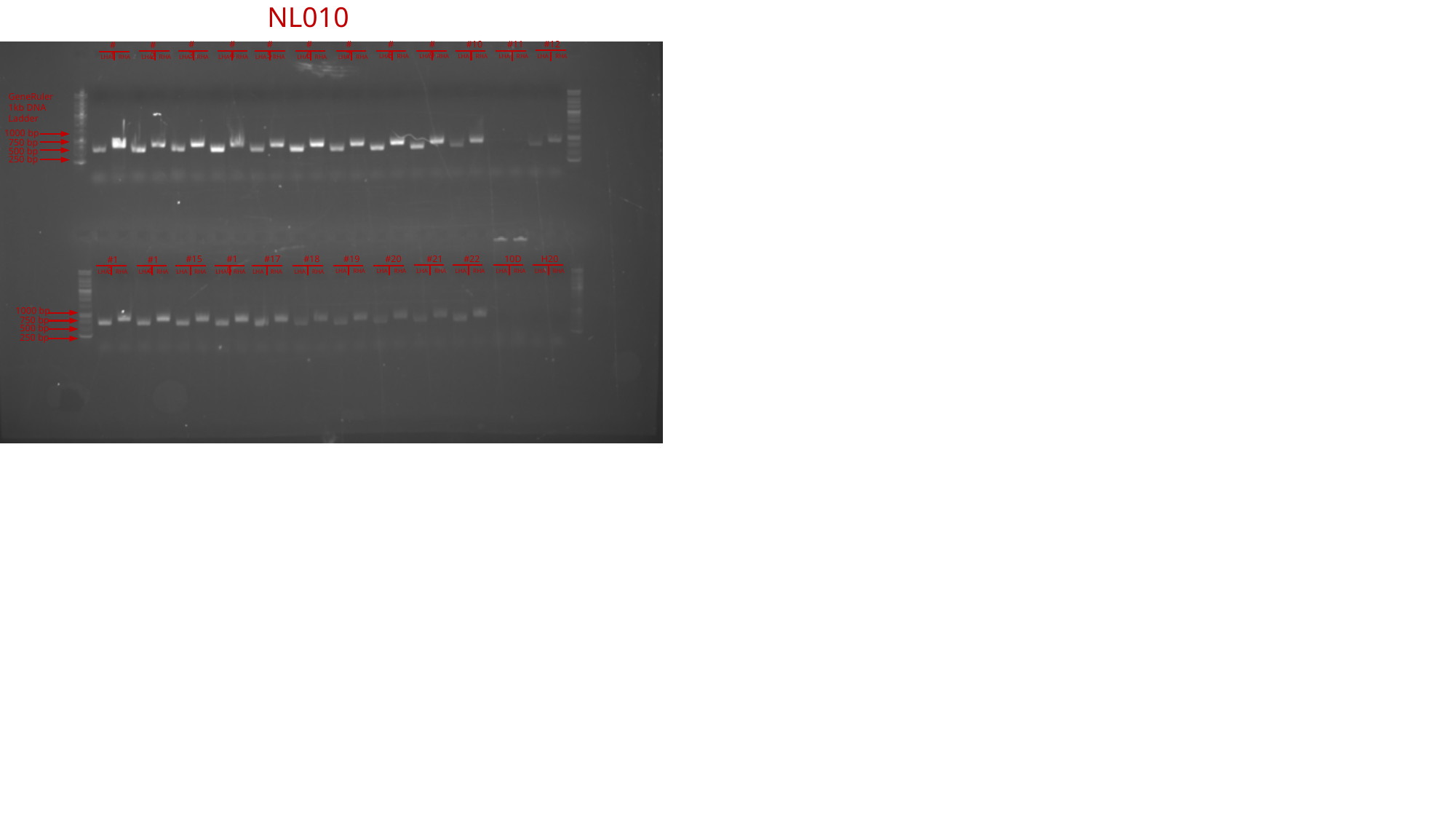

NL010
#3
#4
#5
#6
#7
#8
#9
#10
#11
#12
#1
#2
LHA RHA
LHA RHA
LHA RHA
LHA RHA
LHA RHA
LHA RHA
LHA RHA
LHA RHA
LHA RHA
LHA RHA
LHA RHA
LHA RHA
GeneRuler 1kb DNA Ladder
1000 bp
750 bp
500 bp
250 bp
#15
#16
#17
#18
#19
#20
#21
#22
10D
H20
#13
#14
LHA RHA
LHA RHA
LHA RHA
LHA RHA
LHA RHA
LHA RHA
LHA RHA
LHA RHA
LHA RHA
LHA RHA
LHA RHA
LHA RHA
1000 bp
750 bp
500 bp
250 bp

### Slide 15
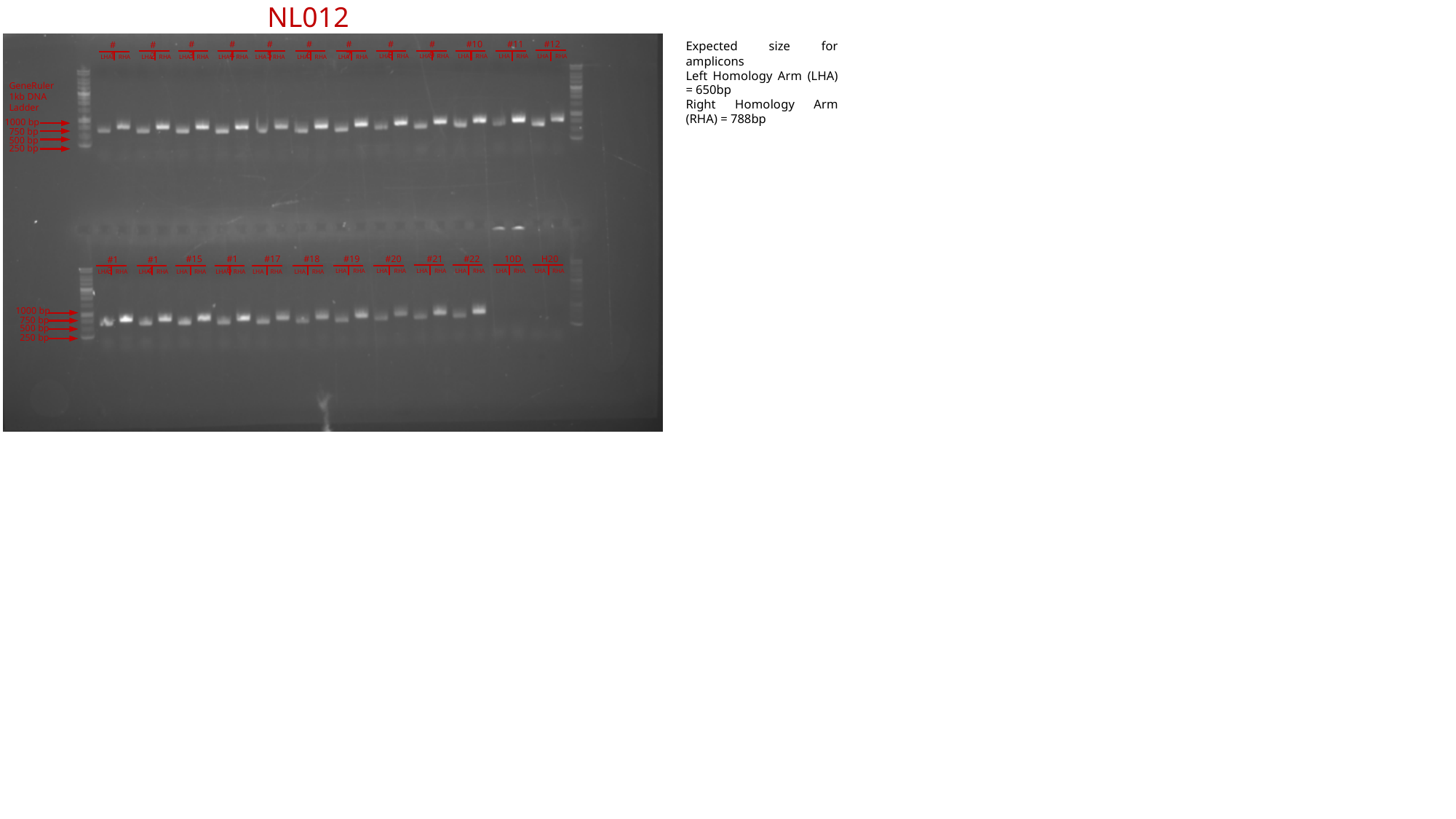

NL012
#3
#4
#5
#6
#7
#8
#9
#10
#11
#12
Expected size for amplicons
Left Homology Arm (LHA) = 650bp
Right Homology Arm (RHA) = 788bp
#1
#2
LHA RHA
LHA RHA
LHA RHA
LHA RHA
LHA RHA
LHA RHA
LHA RHA
LHA RHA
LHA RHA
LHA RHA
LHA RHA
LHA RHA
GeneRuler 1kb DNA Ladder
1000 bp
750 bp
500 bp
250 bp
#15
#16
#17
#18
#19
#20
#21
#22
10D
H20
#13
#14
LHA RHA
LHA RHA
LHA RHA
LHA RHA
LHA RHA
LHA RHA
LHA RHA
LHA RHA
LHA RHA
LHA RHA
LHA RHA
LHA RHA
1000 bp
750 bp
500 bp
250 bp

### Slide 16
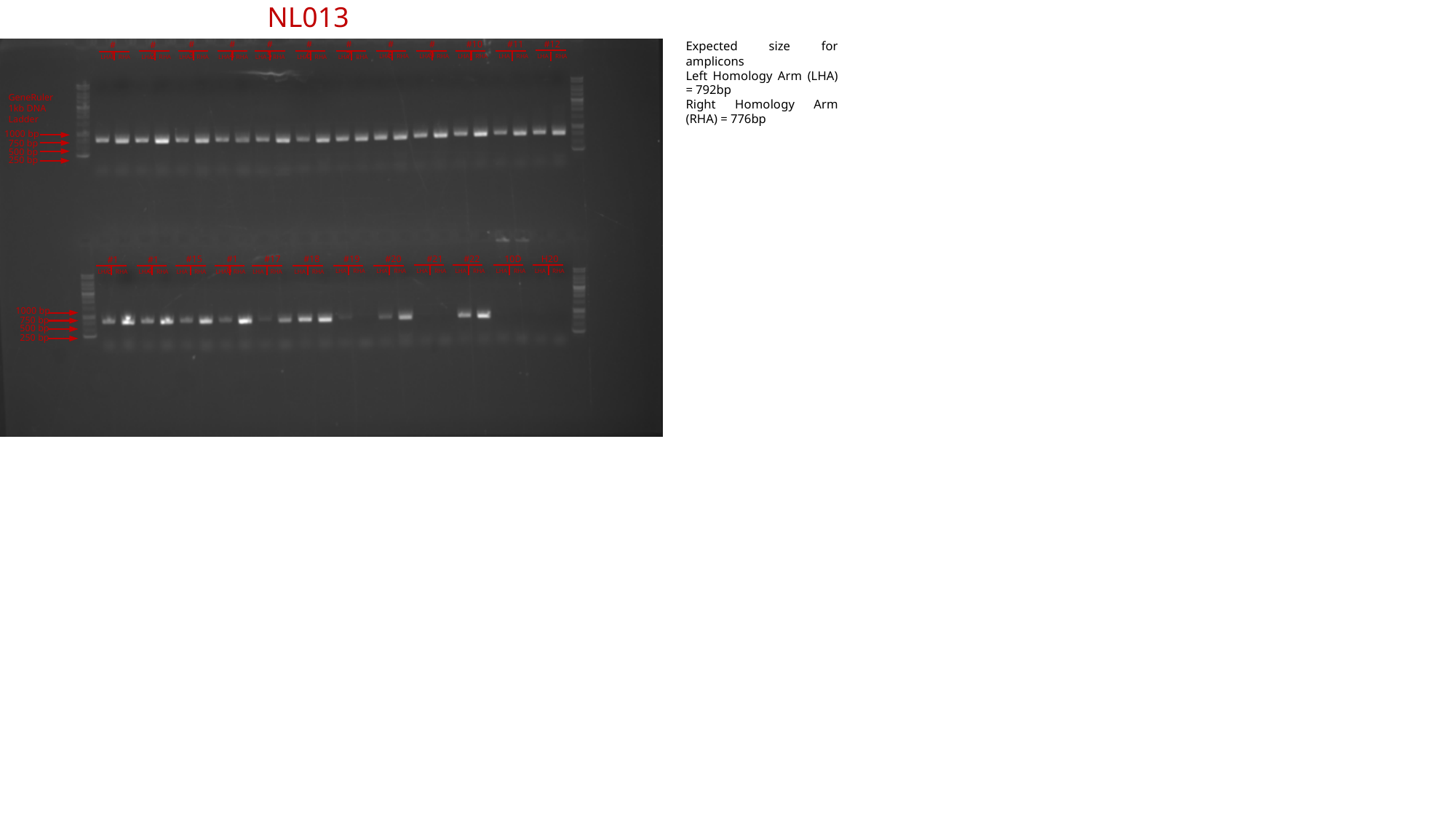

NL013
#3
#4
#5
#6
#7
#8
#9
#10
#11
#12
Expected size for amplicons
Left Homology Arm (LHA) = 792bp
Right Homology Arm (RHA) = 776bp
#1
#2
LHA RHA
LHA RHA
LHA RHA
LHA RHA
LHA RHA
LHA RHA
LHA RHA
LHA RHA
LHA RHA
LHA RHA
LHA RHA
LHA RHA
GeneRuler 1kb DNA Ladder
1000 bp
750 bp
500 bp
250 bp
#15
#16
#17
#18
#19
#20
#21
#22
10D
H20
#13
#14
LHA RHA
LHA RHA
LHA RHA
LHA RHA
LHA RHA
LHA RHA
LHA RHA
LHA RHA
LHA RHA
LHA RHA
LHA RHA
LHA RHA
1000 bp
750 bp
500 bp
250 bp

### Slide 17
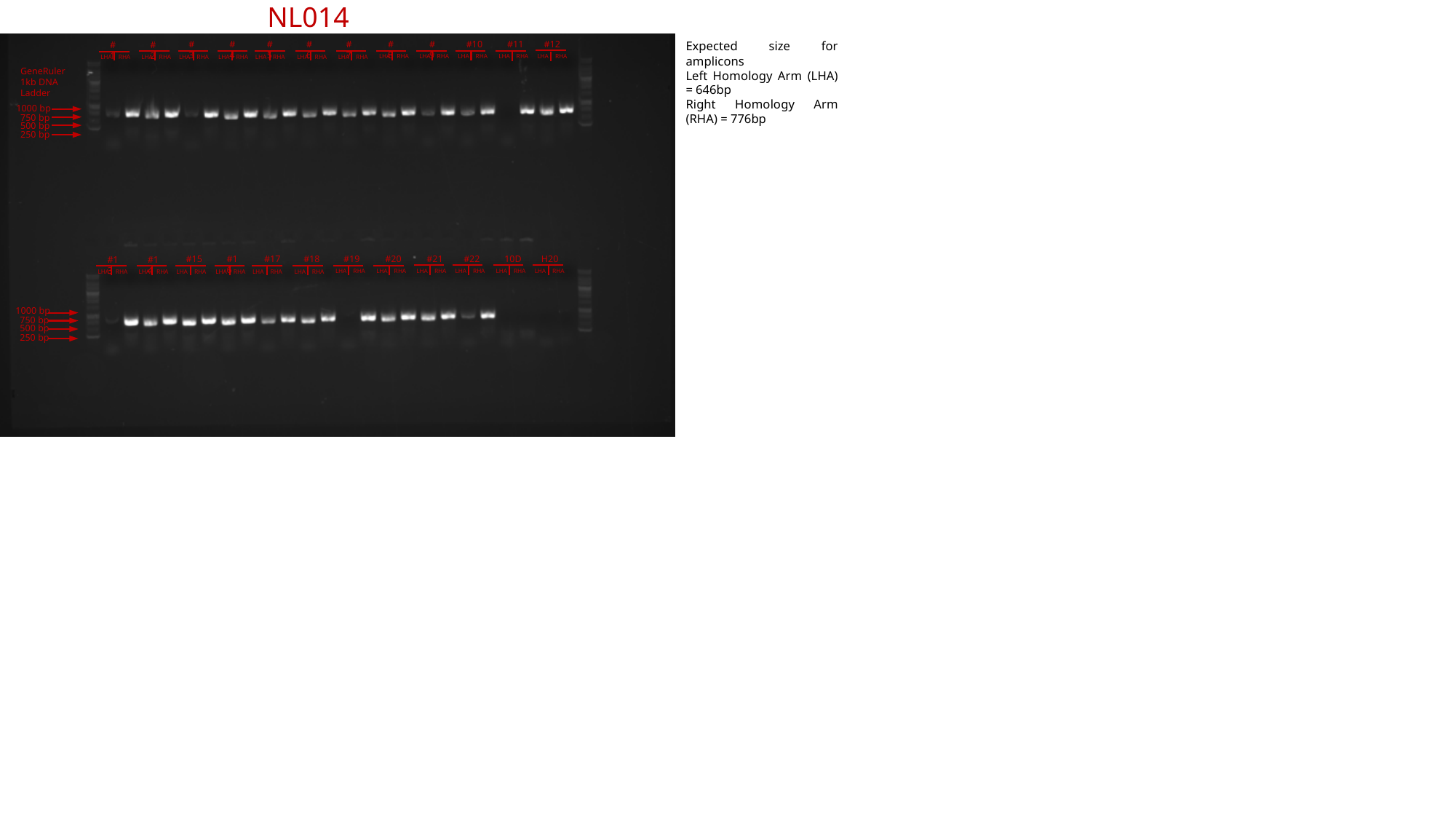

NL014
#3
#4
#5
#6
#7
#8
#9
#10
#11
#12
Expected size for amplicons
Left Homology Arm (LHA) = 646bp
Right Homology Arm (RHA) = 776bp
#1
#2
LHA RHA
LHA RHA
LHA RHA
LHA RHA
LHA RHA
LHA RHA
LHA RHA
LHA RHA
LHA RHA
LHA RHA
LHA RHA
LHA RHA
GeneRuler 1kb DNA Ladder
1000 bp
750 bp
500 bp
250 bp
#15
#16
#17
#18
#19
#20
#21
#22
10D
H20
#13
#14
LHA RHA
LHA RHA
LHA RHA
LHA RHA
LHA RHA
LHA RHA
LHA RHA
LHA RHA
LHA RHA
LHA RHA
LHA RHA
LHA RHA
1000 bp
750 bp
500 bp
250 bp

### Slide 18
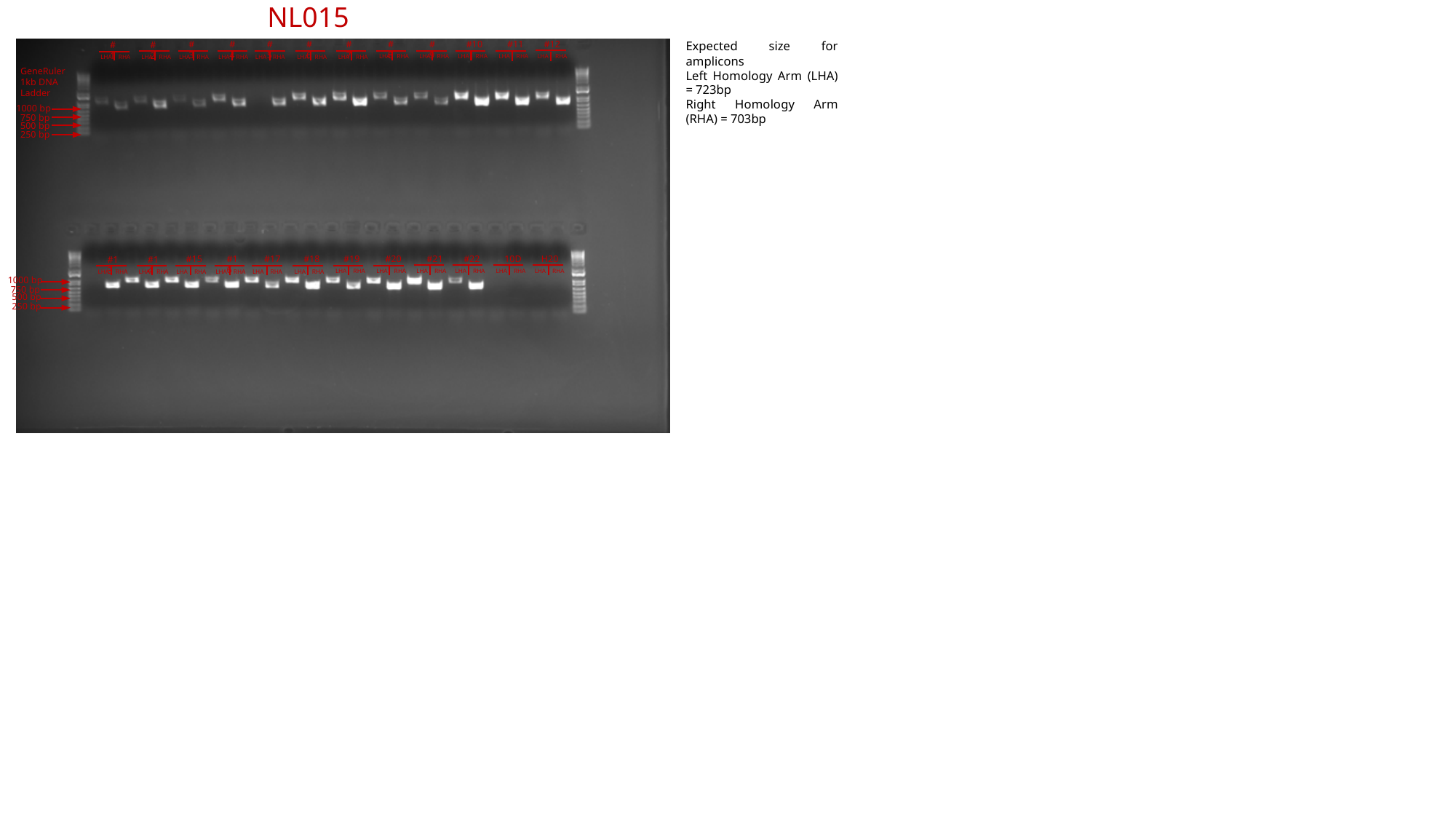

NL015
#3
#4
#5
#6
#7
#8
#9
#10
#11
#12
Expected size for amplicons
Left Homology Arm (LHA) = 723bp
Right Homology Arm (RHA) = 703bp
#1
#2
LHA RHA
LHA RHA
LHA RHA
LHA RHA
LHA RHA
LHA RHA
LHA RHA
LHA RHA
LHA RHA
LHA RHA
LHA RHA
LHA RHA
GeneRuler 1kb DNA Ladder
1000 bp
750 bp
500 bp
250 bp
#15
#16
#17
#18
#19
#20
#21
#22
10D
H20
#13
#14
LHA RHA
LHA RHA
LHA RHA
LHA RHA
LHA RHA
LHA RHA
LHA RHA
LHA RHA
LHA RHA
LHA RHA
LHA RHA
LHA RHA
1000 bp
750 bp
500 bp
250 bp

### Slide 19
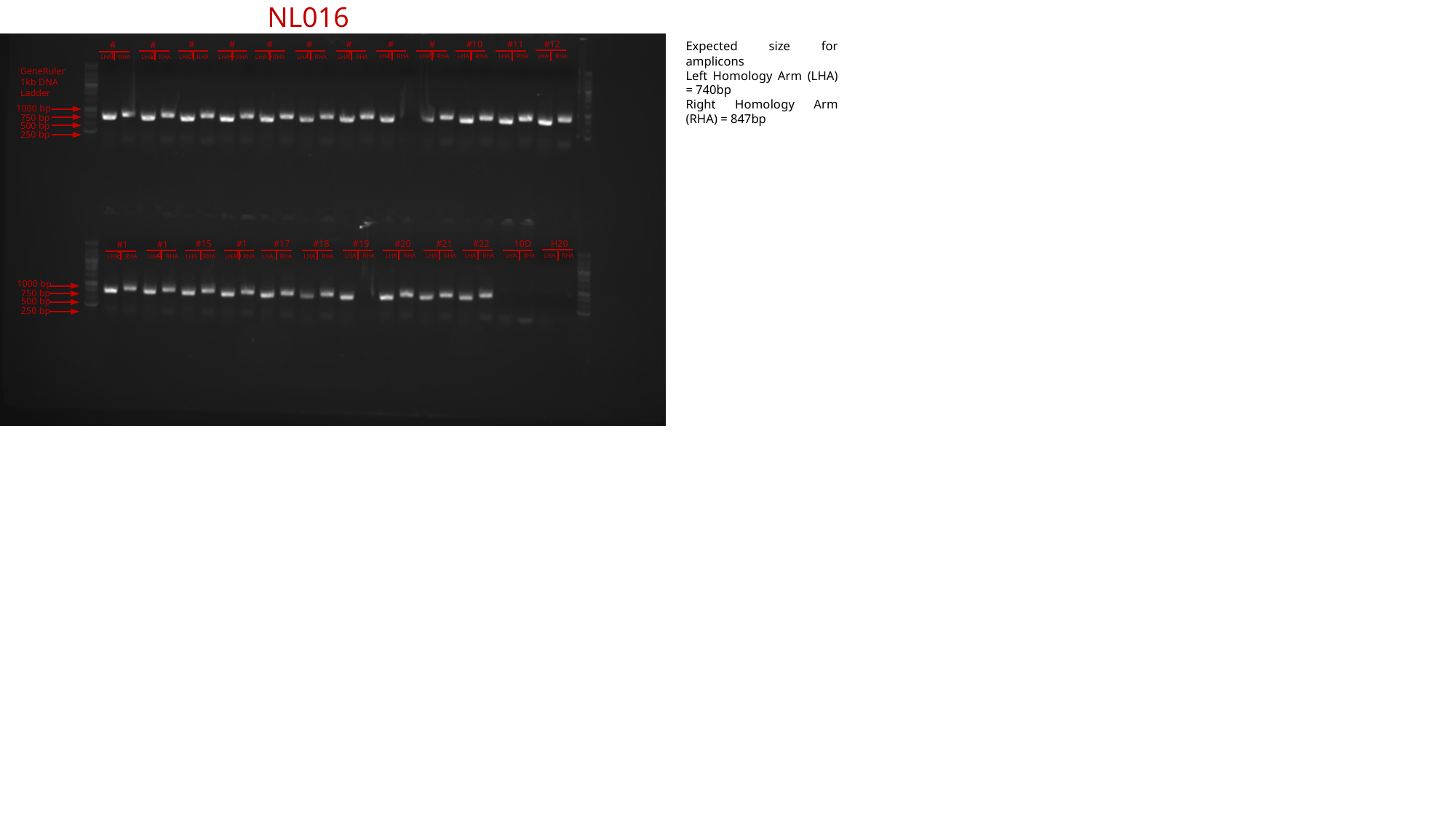

NL016
#3
#4
#5
#6
#7
#8
#9
#10
#11
#12
Expected size for amplicons
Left Homology Arm (LHA) = 740bp
Right Homology Arm (RHA) = 847bp
#1
#2
LHA RHA
LHA RHA
LHA RHA
LHA RHA
LHA RHA
LHA RHA
LHA RHA
LHA RHA
LHA RHA
LHA RHA
LHA RHA
LHA RHA
GeneRuler 1kb DNA Ladder
1000 bp
750 bp
500 bp
250 bp
#15
#16
#17
#18
#19
#20
#21
#22
10D
H20
#13
#14
LHA RHA
LHA RHA
LHA RHA
LHA RHA
LHA RHA
LHA RHA
LHA RHA
LHA RHA
LHA RHA
LHA RHA
LHA RHA
LHA RHA
1000 bp
750 bp
500 bp
250 bp

### Slide 20
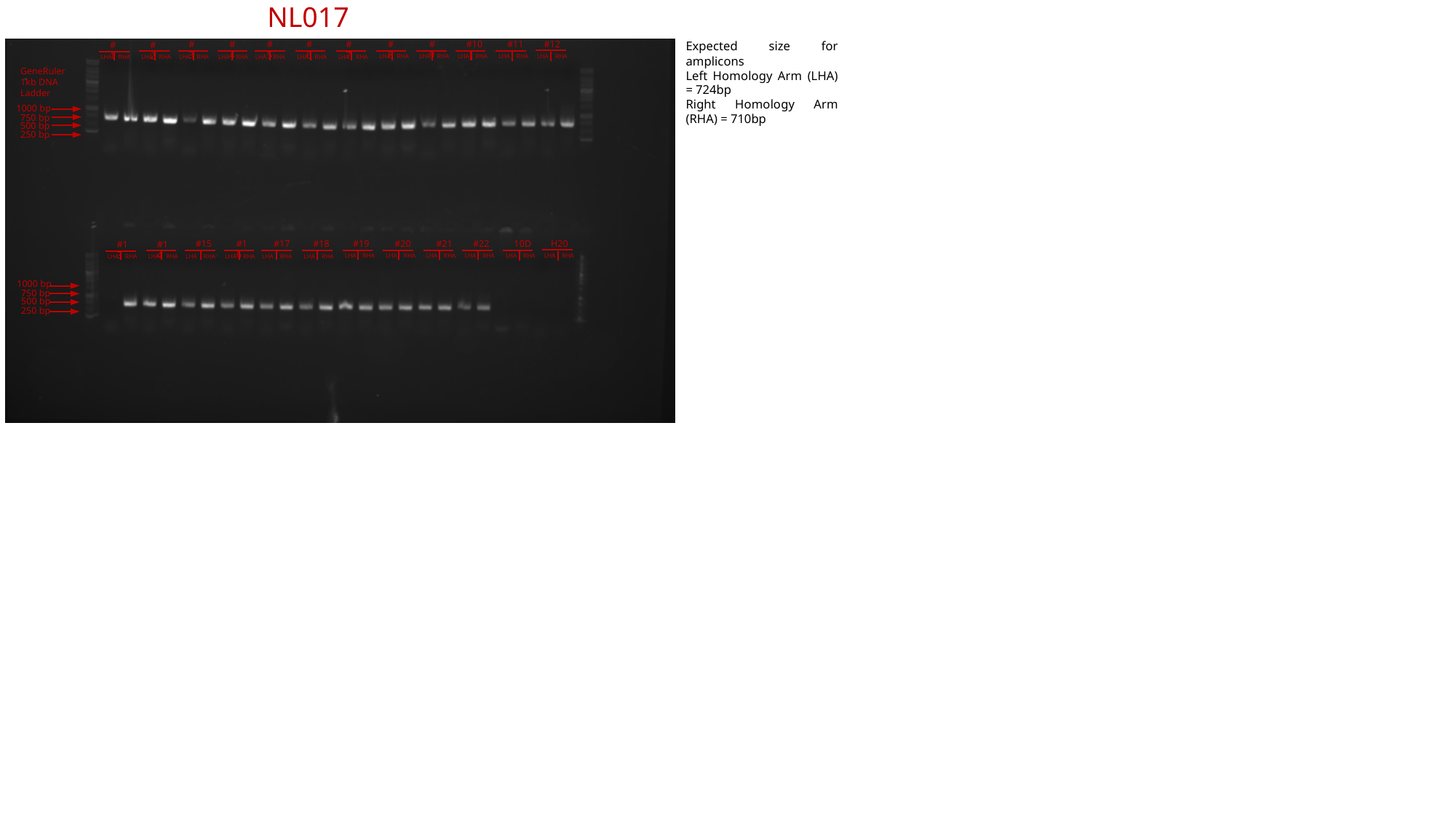

NL017
#3
#4
#5
#6
#7
#8
#9
#10
#11
#12
Expected size for amplicons
Left Homology Arm (LHA) = 724bp
Right Homology Arm (RHA) = 710bp
#1
#2
LHA RHA
LHA RHA
LHA RHA
LHA RHA
LHA RHA
LHA RHA
LHA RHA
LHA RHA
LHA RHA
LHA RHA
LHA RHA
LHA RHA
GeneRuler 1kb DNA Ladder
1000 bp
750 bp
500 bp
250 bp
#15
#16
#17
#18
#19
#20
#21
#22
10D
H20
#13
#14
LHA RHA
LHA RHA
LHA RHA
LHA RHA
LHA RHA
LHA RHA
LHA RHA
LHA RHA
LHA RHA
LHA RHA
LHA RHA
LHA RHA
1000 bp
750 bp
500 bp
250 bp

### Slide 21
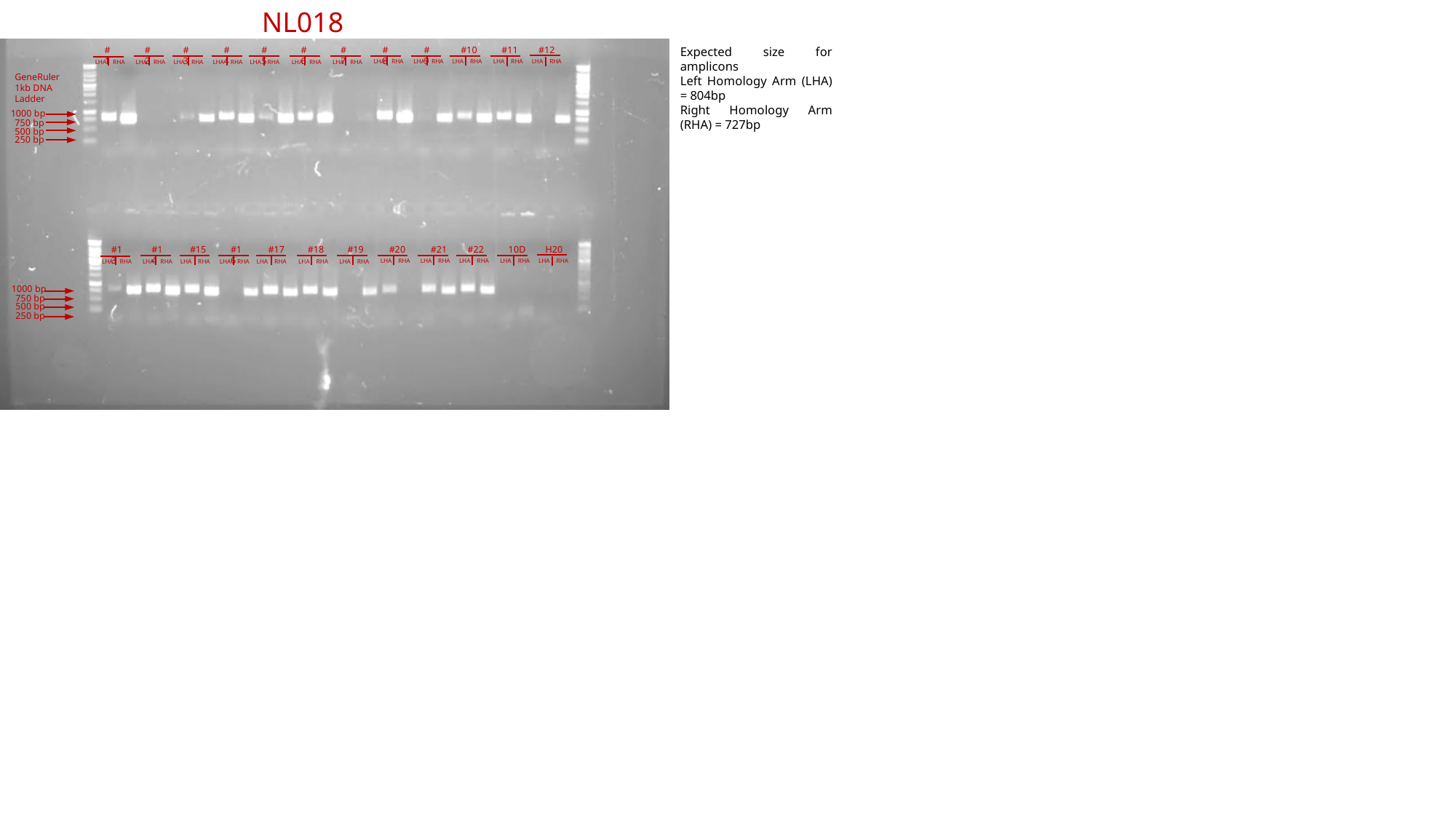

NL018
#3
#4
#5
#6
#7
#8
#9
#10
#11
#12
Expected size for amplicons
Left Homology Arm (LHA) = 804bp
Right Homology Arm (RHA) = 727bp
#1
#2
LHA RHA
LHA RHA
LHA RHA
LHA RHA
LHA RHA
LHA RHA
LHA RHA
LHA RHA
LHA RHA
LHA RHA
LHA RHA
LHA RHA
GeneRuler 1kb DNA Ladder
1000 bp
750 bp
500 bp
250 bp
#15
#16
#17
#18
#19
#20
#21
#22
10D
H20
#13
#14
LHA RHA
LHA RHA
LHA RHA
LHA RHA
LHA RHA
LHA RHA
LHA RHA
LHA RHA
LHA RHA
LHA RHA
LHA RHA
LHA RHA
1000 bp
750 bp
500 bp
250 bp

### Slide 22
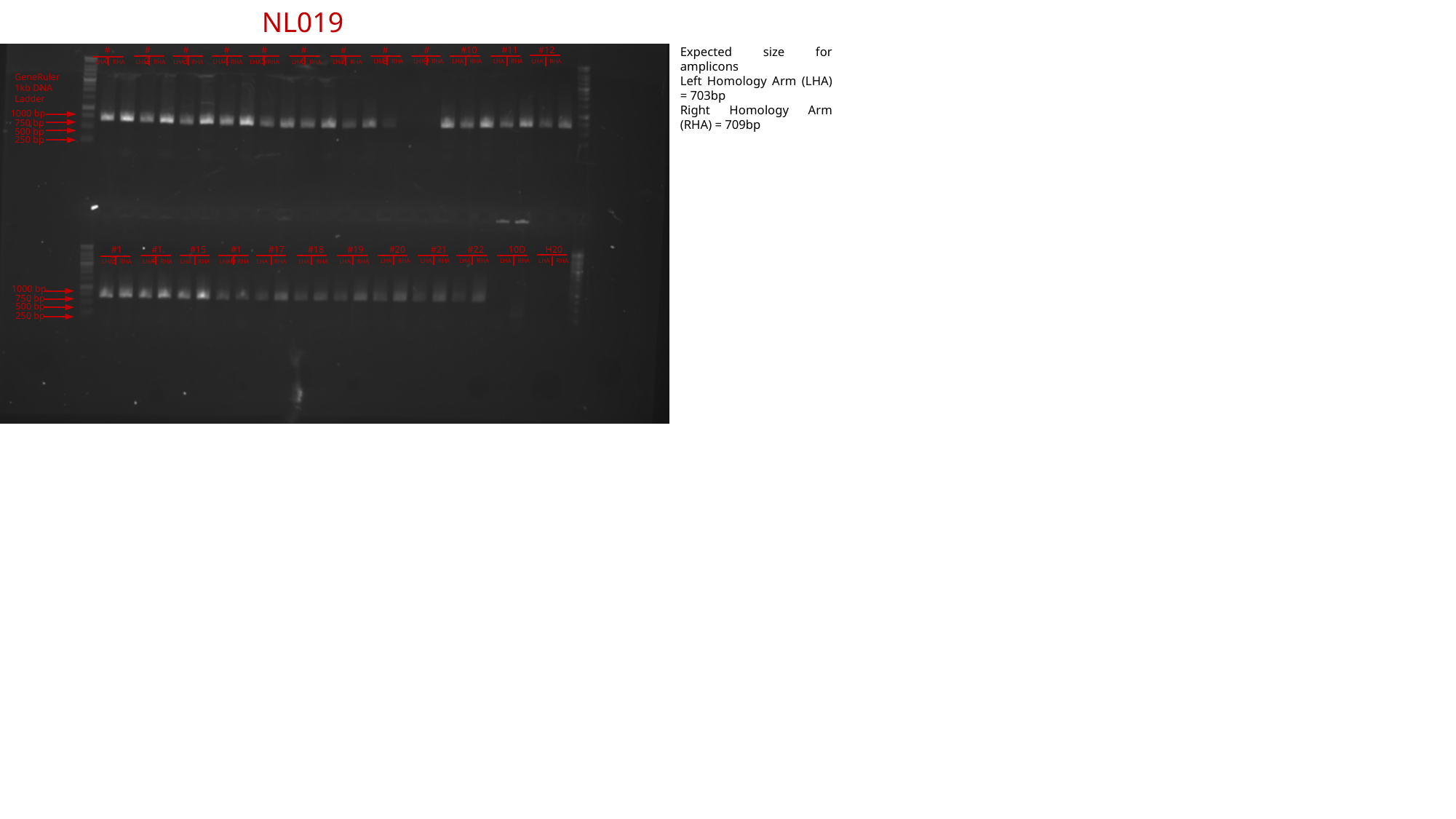

NL019
#3
#4
#5
#6
#7
#8
#9
#10
#11
#12
Expected size for amplicons
Left Homology Arm (LHA) = 703bp
Right Homology Arm (RHA) = 709bp
#1
#2
LHA RHA
LHA RHA
LHA RHA
LHA RHA
LHA RHA
LHA RHA
LHA RHA
LHA RHA
LHA RHA
LHA RHA
LHA RHA
LHA RHA
GeneRuler 1kb DNA Ladder
1000 bp
750 bp
500 bp
250 bp
#15
#16
#17
#18
#19
#20
#21
#22
10D
H20
#13
#14
LHA RHA
LHA RHA
LHA RHA
LHA RHA
LHA RHA
LHA RHA
LHA RHA
LHA RHA
LHA RHA
LHA RHA
LHA RHA
LHA RHA
1000 bp
750 bp
500 bp
250 bp

### Slide 23
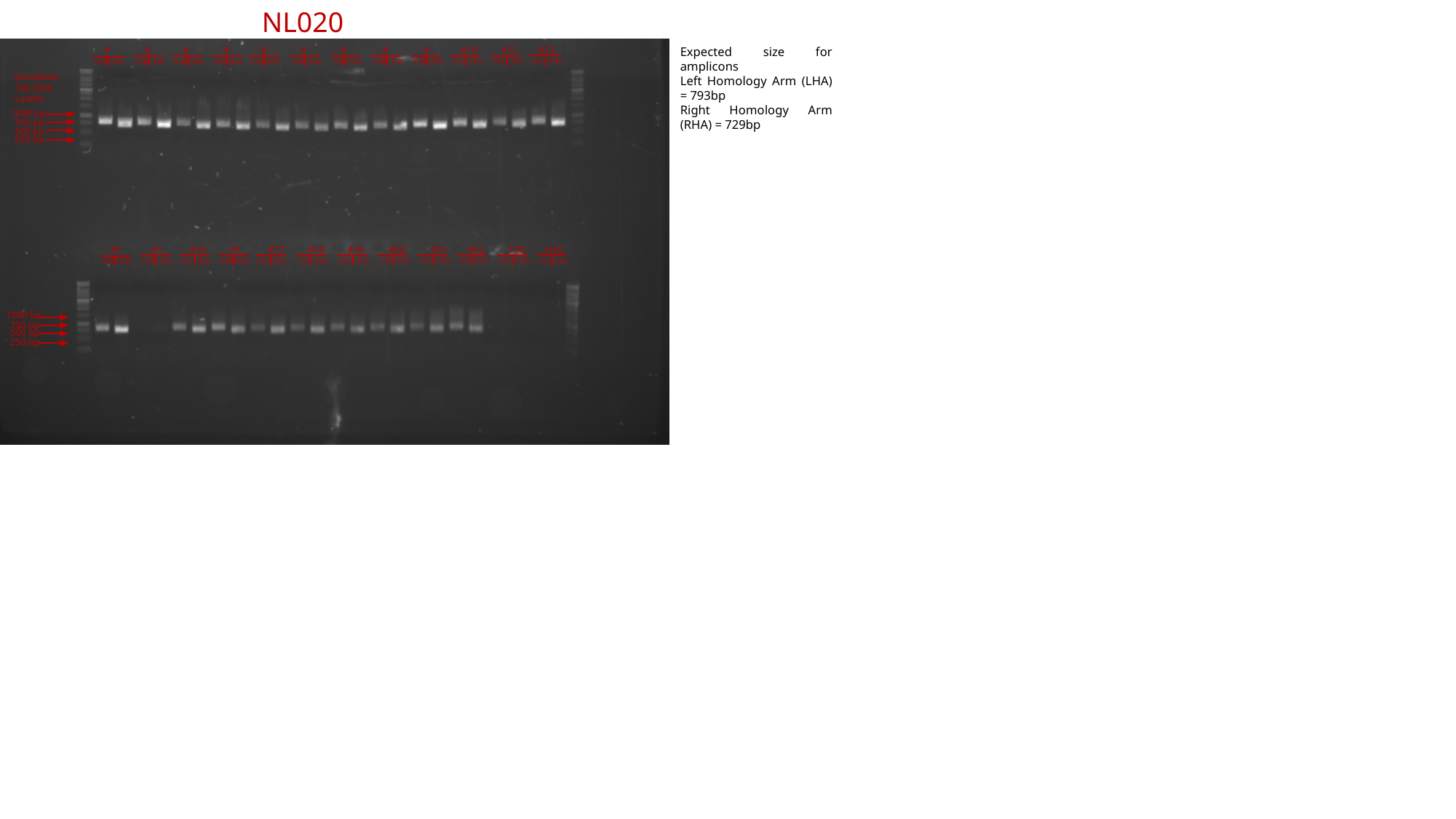

NL020
#3
#4
#5
#6
#7
#8
#9
#10
#11
#12
Expected size for amplicons
Left Homology Arm (LHA) = 793bp
Right Homology Arm (RHA) = 729bp
#1
#2
LHA RHA
LHA RHA
LHA RHA
LHA RHA
LHA RHA
LHA RHA
LHA RHA
LHA RHA
LHA RHA
LHA RHA
LHA RHA
LHA RHA
GeneRuler 1kb DNA Ladder
1000 bp
750 bp
500 bp
250 bp
#15
#16
#17
#18
#19
#20
#21
#22
10D
H20
#13
#14
LHA RHA
LHA RHA
LHA RHA
LHA RHA
LHA RHA
LHA RHA
LHA RHA
LHA RHA
LHA RHA
LHA RHA
LHA RHA
LHA RHA
1000 bp
750 bp
500 bp
250 bp

### Slide 24
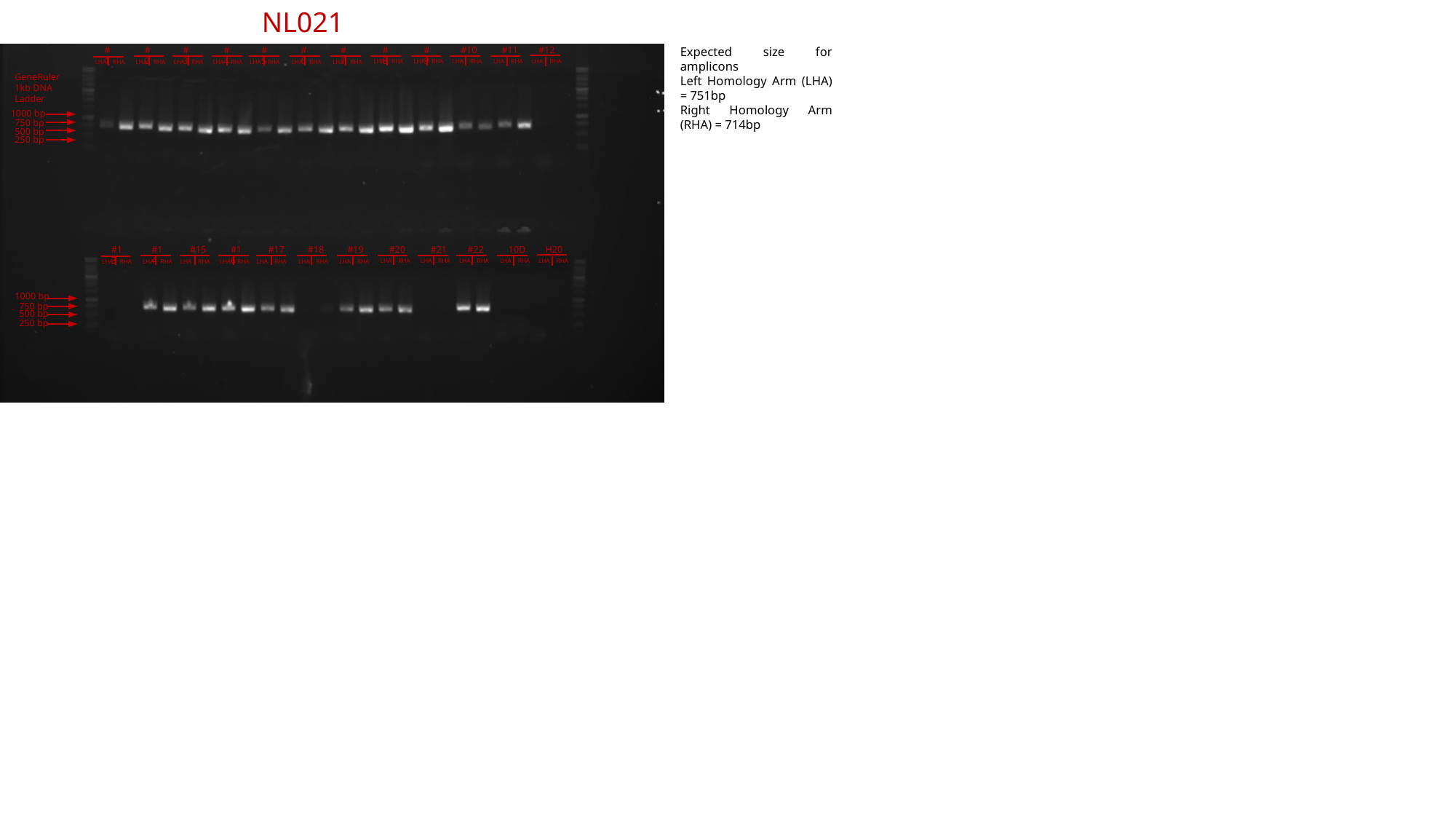

NL021
#3
#4
#5
#6
#7
#8
#9
#10
#11
#12
Expected size for amplicons
Left Homology Arm (LHA) = 751bp
Right Homology Arm (RHA) = 714bp
#1
#2
LHA RHA
LHA RHA
LHA RHA
LHA RHA
LHA RHA
LHA RHA
LHA RHA
LHA RHA
LHA RHA
LHA RHA
LHA RHA
LHA RHA
GeneRuler 1kb DNA Ladder
1000 bp
750 bp
500 bp
250 bp
#15
#16
#17
#18
#19
#20
#21
#22
10D
H20
#13
#14
LHA RHA
LHA RHA
LHA RHA
LHA RHA
LHA RHA
LHA RHA
LHA RHA
LHA RHA
LHA RHA
LHA RHA
LHA RHA
LHA RHA
1000 bp
750 bp
500 bp
250 bp

### Slide 25
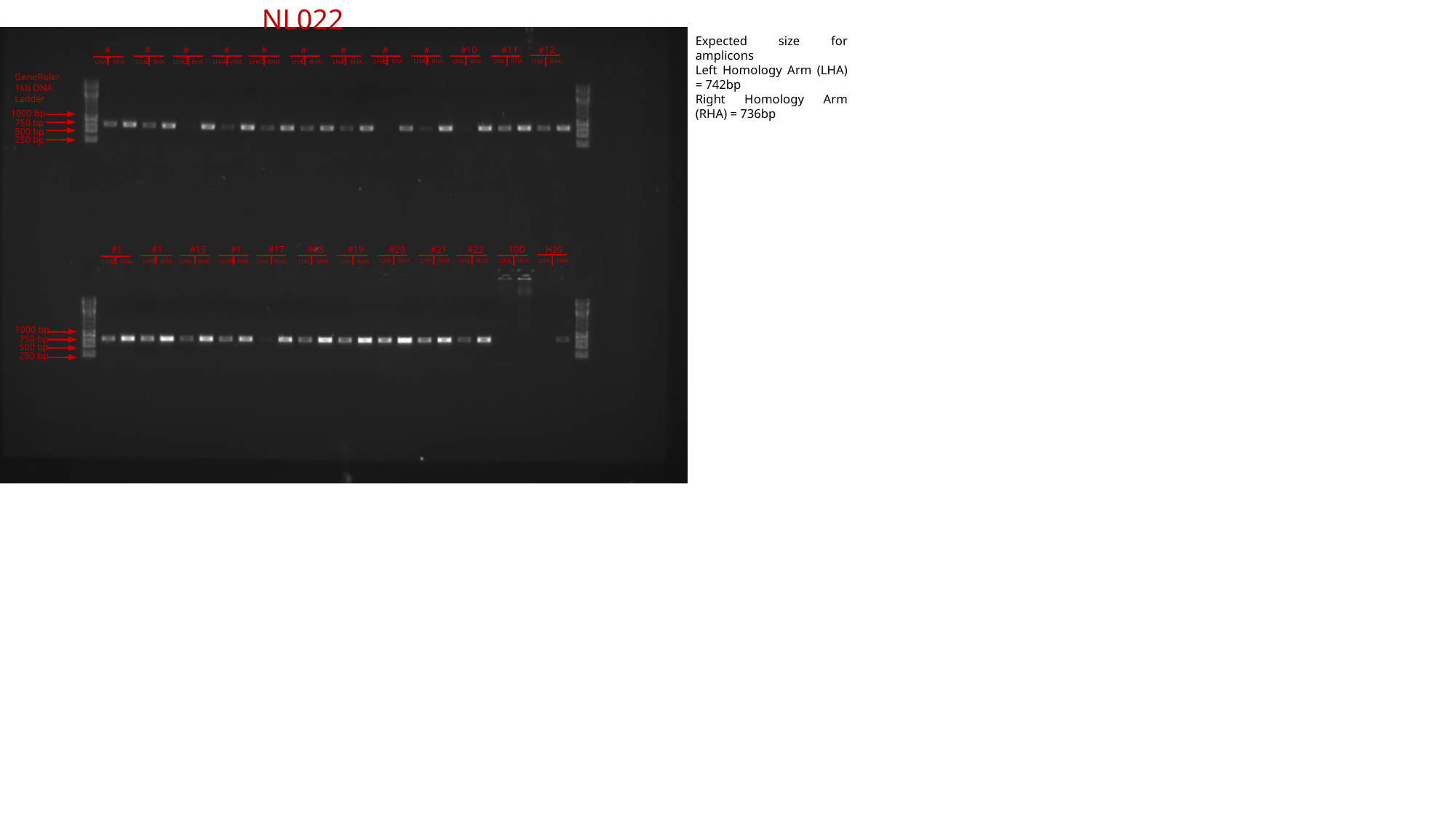

NL022
Expected size for amplicons
Left Homology Arm (LHA) = 742bp
Right Homology Arm (RHA) = 736bp
#3
#4
#5
#6
#7
#8
#9
#10
#11
#12
#1
#2
LHA RHA
LHA RHA
LHA RHA
LHA RHA
LHA RHA
LHA RHA
LHA RHA
LHA RHA
LHA RHA
LHA RHA
LHA RHA
LHA RHA
GeneRuler 1kb DNA Ladder
1000 bp
750 bp
500 bp
250 bp
#15
#16
#17
#18
#19
#20
#21
#22
10D
H20
#13
#14
LHA RHA
LHA RHA
LHA RHA
LHA RHA
LHA RHA
LHA RHA
LHA RHA
LHA RHA
LHA RHA
LHA RHA
LHA RHA
LHA RHA
1000 bp
750 bp
500 bp
250 bp

### Slide 26
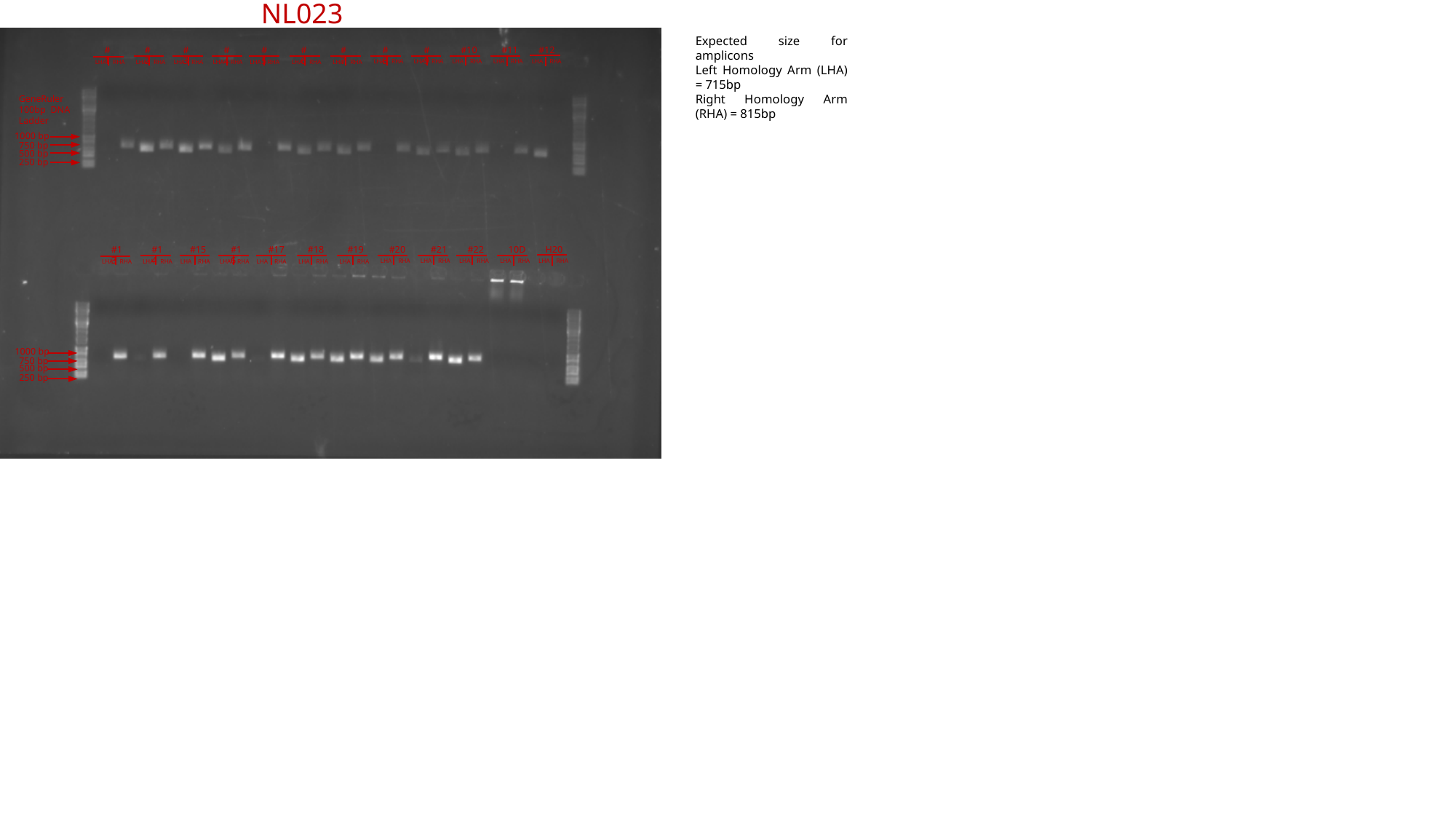

NL023
Expected size for amplicons
Left Homology Arm (LHA) = 715bp
Right Homology Arm (RHA) = 815bp
#3
#4
#5
#6
#7
#8
#9
#10
#11
#12
#1
#2
LHA RHA
LHA RHA
LHA RHA
LHA RHA
LHA RHA
LHA RHA
LHA RHA
LHA RHA
LHA RHA
LHA RHA
LHA RHA
LHA RHA
GeneRuler 100bp DNA Ladder
1000 bp
750 bp
500 bp
250 bp
#15
#16
#17
#18
#19
#20
#21
#22
10D
H20
#13
#14
LHA RHA
LHA RHA
LHA RHA
LHA RHA
LHA RHA
LHA RHA
LHA RHA
LHA RHA
LHA RHA
LHA RHA
LHA RHA
LHA RHA
1000 bp
750 bp
500 bp
250 bp

### Slide 27
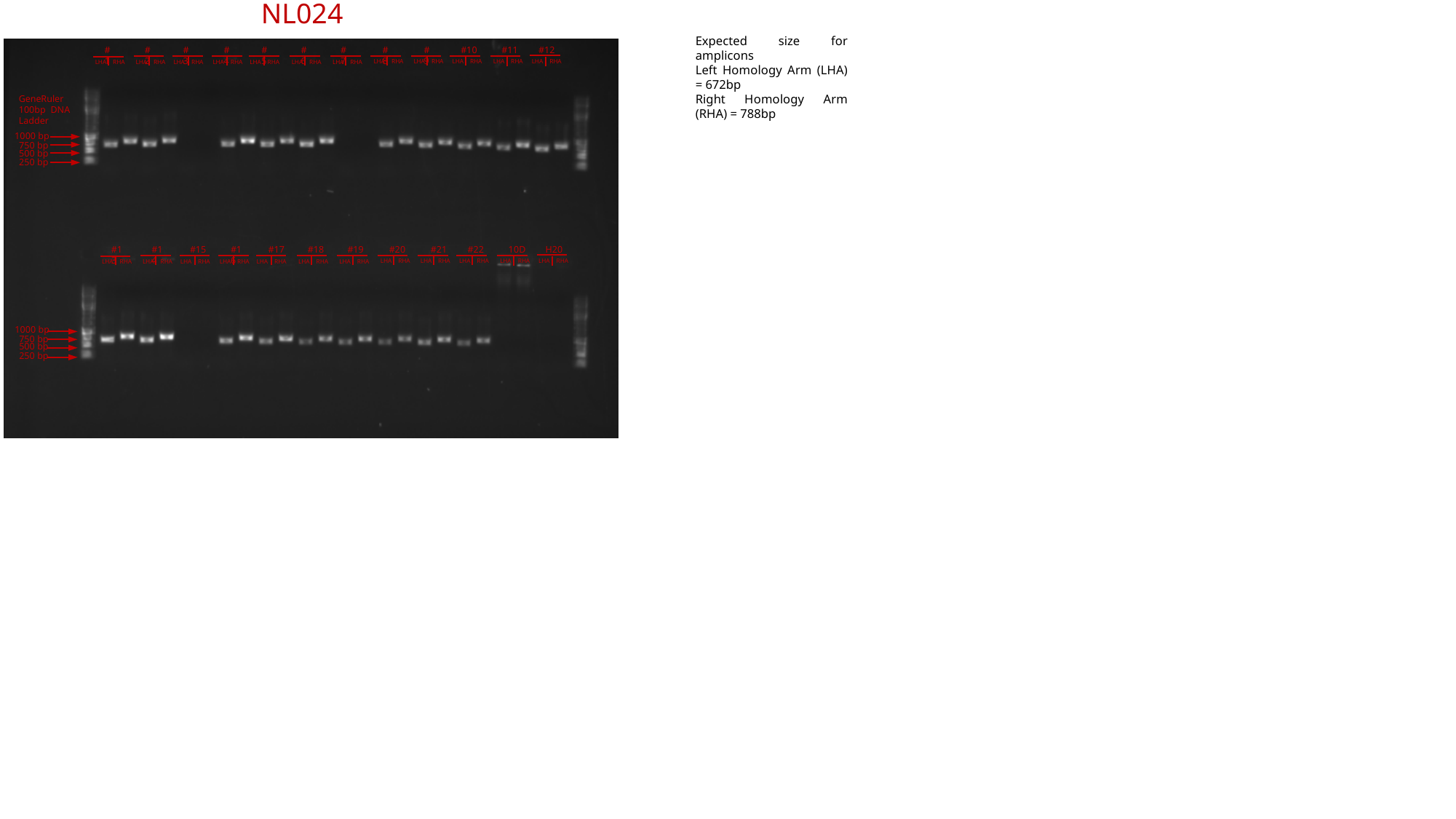

NL024
Expected size for amplicons
Left Homology Arm (LHA) = 672bp
Right Homology Arm (RHA) = 788bp
#3
#4
#5
#6
#7
#8
#9
#10
#11
#12
#1
#2
LHA RHA
LHA RHA
LHA RHA
LHA RHA
LHA RHA
LHA RHA
LHA RHA
LHA RHA
LHA RHA
LHA RHA
LHA RHA
LHA RHA
GeneRuler 100bp DNA Ladder
1000 bp
750 bp
500 bp
250 bp
#15
#16
#17
#18
#19
#20
#21
#22
10D
H20
#13
#14
LHA RHA
LHA RHA
LHA RHA
LHA RHA
LHA RHA
LHA RHA
LHA RHA
LHA RHA
LHA RHA
LHA RHA
LHA RHA
LHA RHA
1000 bp
750 bp
500 bp
250 bp

### Slide 28
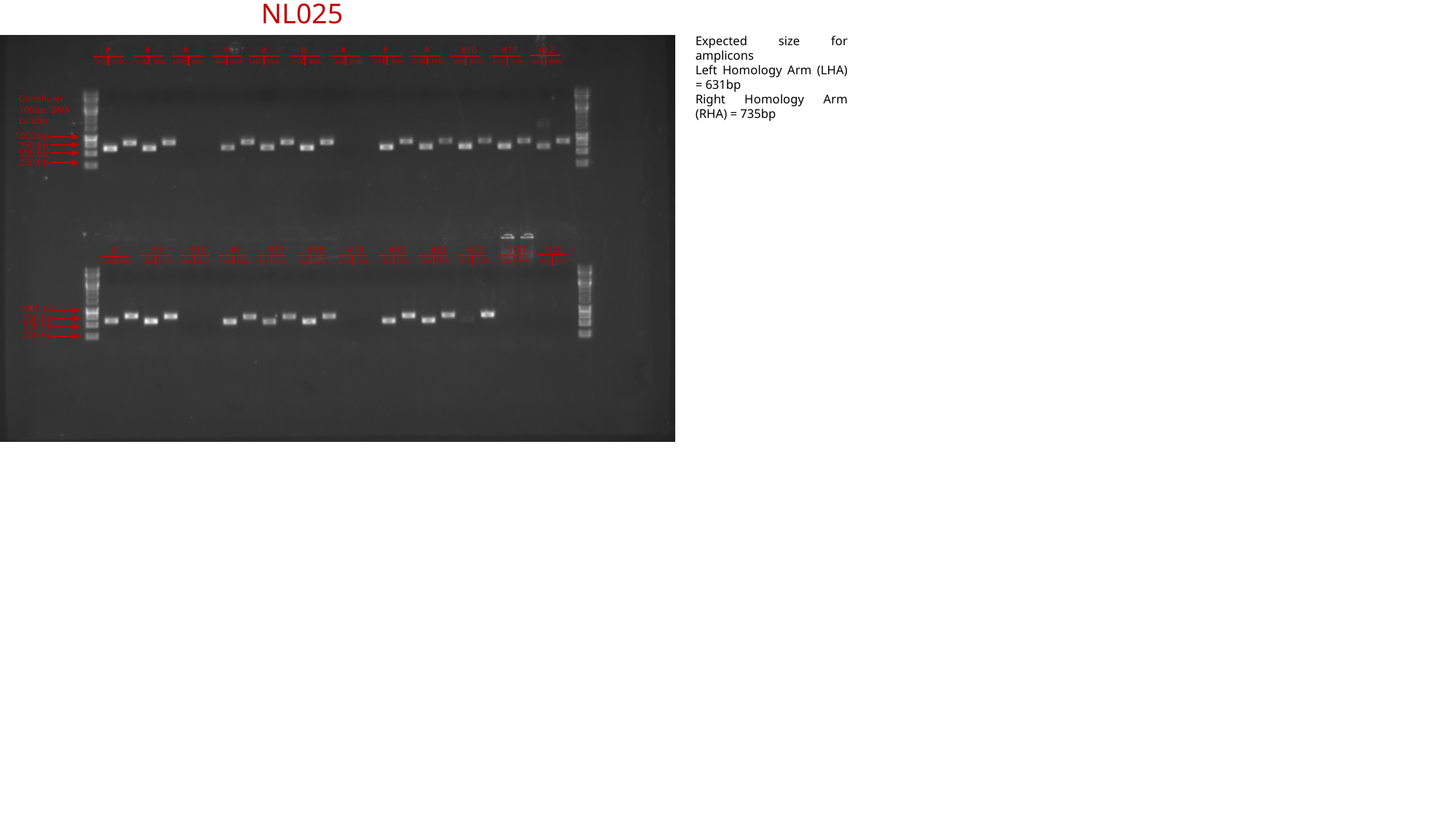

NL025
Expected size for amplicons
Left Homology Arm (LHA) = 631bp
Right Homology Arm (RHA) = 735bp
#3
#4
#5
#6
#7
#8
#9
#10
#11
#12
#1
#2
LHA RHA
LHA RHA
LHA RHA
LHA RHA
LHA RHA
LHA RHA
LHA RHA
LHA RHA
LHA RHA
LHA RHA
LHA RHA
LHA RHA
GeneRuler 100bp DNA Ladder
1000 bp
750 bp
500 bp
250 bp
#15
#16
#17
#18
#19
#20
#21
#22
10D
H20
#13
#14
LHA RHA
LHA RHA
LHA RHA
LHA RHA
LHA RHA
LHA RHA
LHA RHA
LHA RHA
LHA RHA
LHA RHA
LHA RHA
LHA RHA
1000 bp
750 bp
500 bp
250 bp

### Slide 29
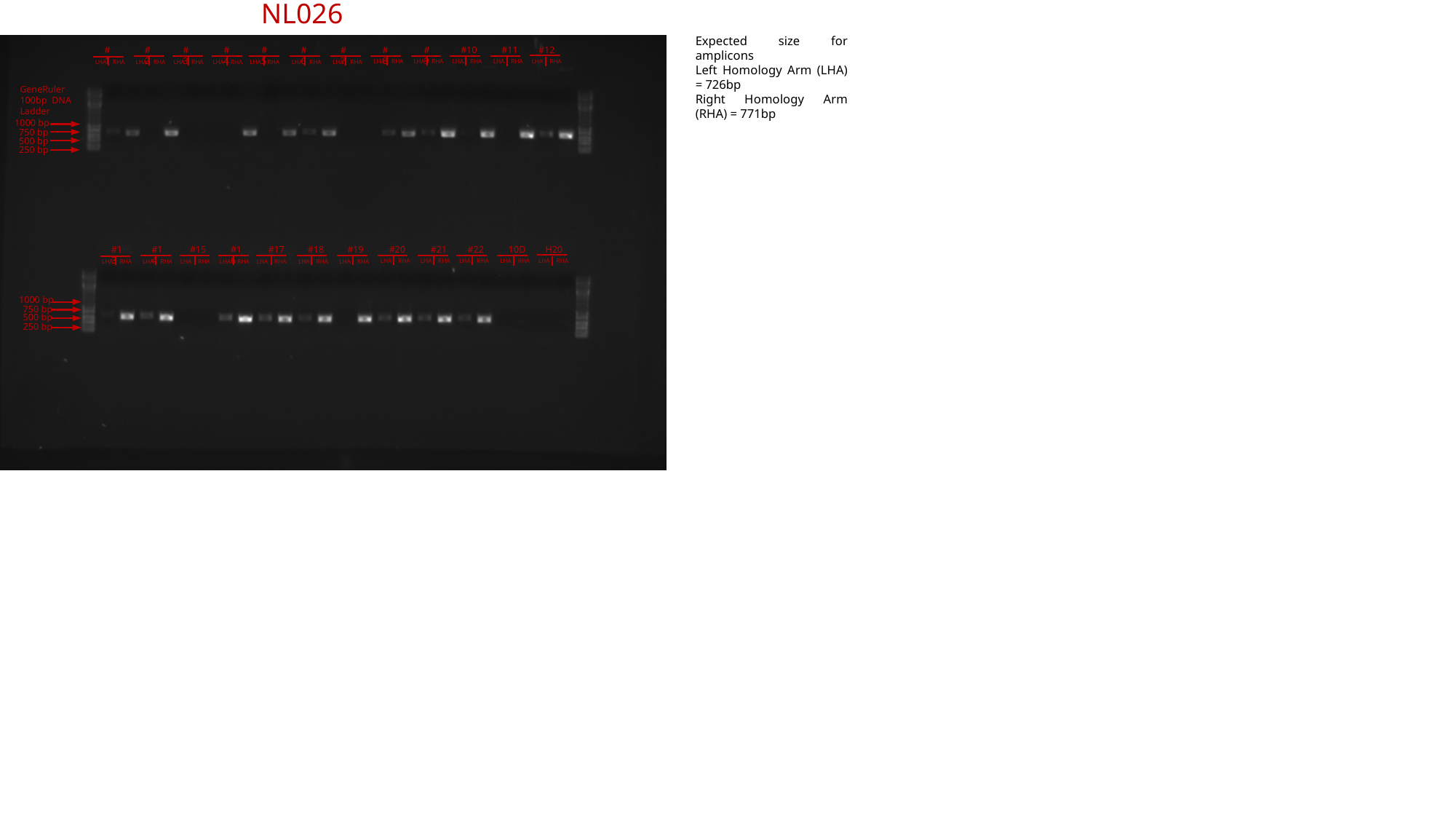

NL026
Expected size for amplicons
Left Homology Arm (LHA) = 726bp
Right Homology Arm (RHA) = 771bp
#3
#4
#5
#6
#7
#8
#9
#10
#11
#12
#1
#2
LHA RHA
LHA RHA
LHA RHA
LHA RHA
LHA RHA
LHA RHA
LHA RHA
LHA RHA
LHA RHA
LHA RHA
LHA RHA
LHA RHA
GeneRuler 100bp DNA Ladder
1000 bp
750 bp
500 bp
250 bp
#15
#16
#17
#18
#19
#20
#21
#22
10D
H20
#13
#14
LHA RHA
LHA RHA
LHA RHA
LHA RHA
LHA RHA
LHA RHA
LHA RHA
LHA RHA
LHA RHA
LHA RHA
LHA RHA
LHA RHA
1000 bp
750 bp
500 bp
250 bp

### Slide 30
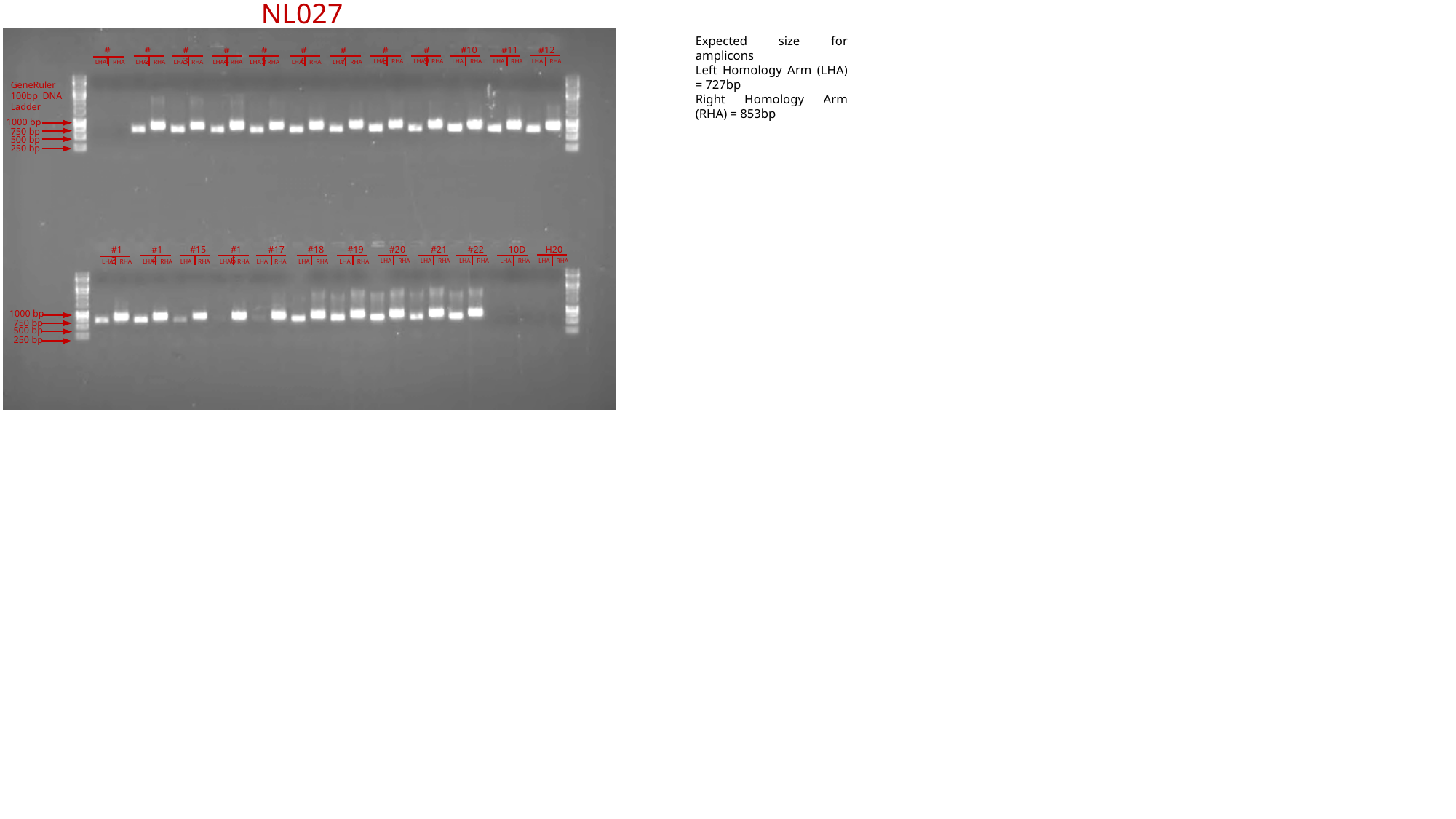

NL027
Expected size for amplicons
Left Homology Arm (LHA) = 727bp
Right Homology Arm (RHA) = 853bp
#3
#4
#5
#6
#7
#8
#9
#10
#11
#12
#1
#2
LHA RHA
LHA RHA
LHA RHA
LHA RHA
LHA RHA
LHA RHA
LHA RHA
LHA RHA
LHA RHA
LHA RHA
LHA RHA
LHA RHA
GeneRuler 100bp DNA Ladder
1000 bp
750 bp
500 bp
250 bp
#15
#16
#17
#18
#19
#20
#21
#22
10D
H20
#13
#14
LHA RHA
LHA RHA
LHA RHA
LHA RHA
LHA RHA
LHA RHA
LHA RHA
LHA RHA
LHA RHA
LHA RHA
LHA RHA
LHA RHA
1000 bp
750 bp
500 bp
250 bp

### Slide 31
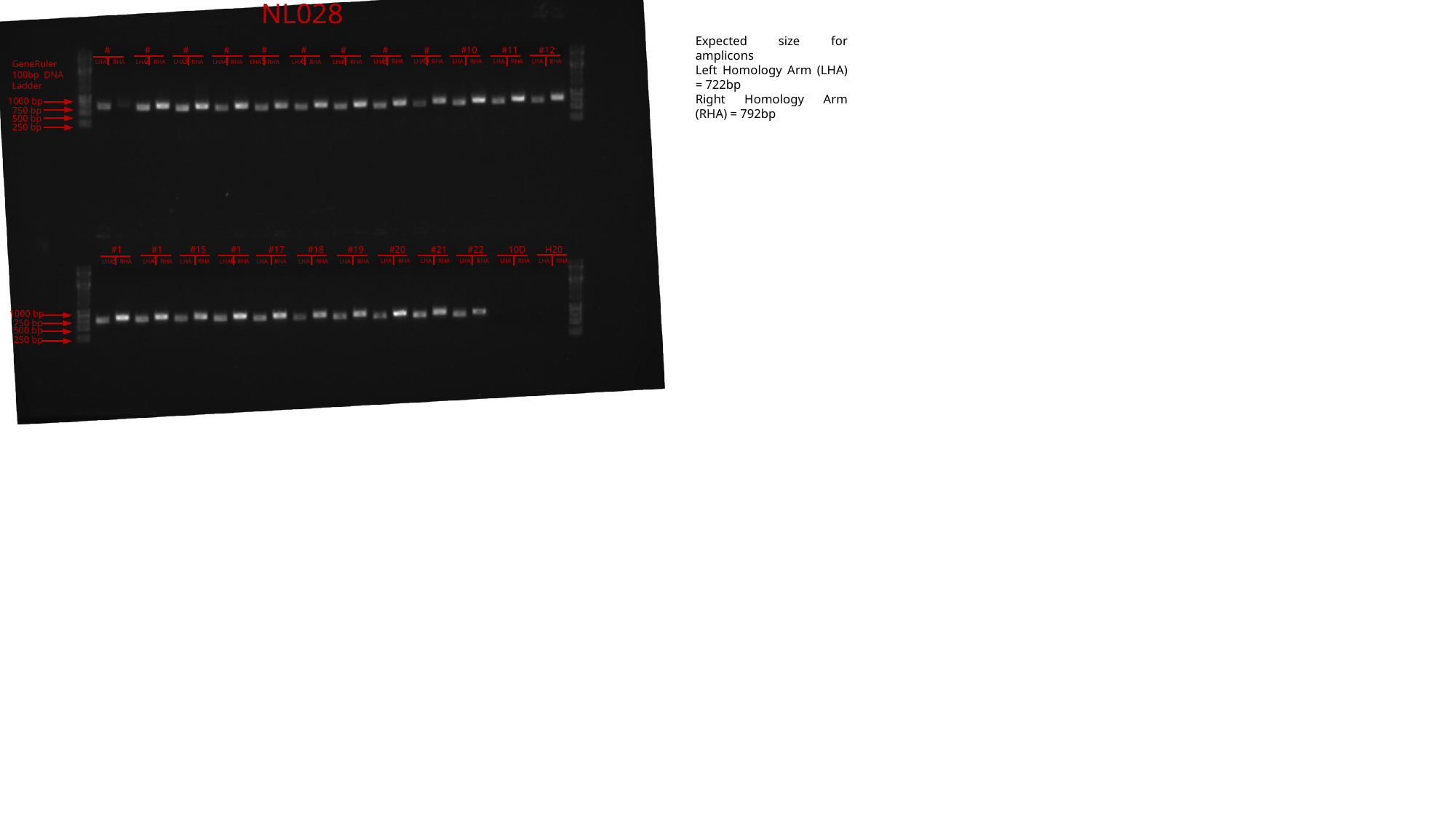

NL028
Expected size for amplicons
Left Homology Arm (LHA) = 722bp
Right Homology Arm (RHA) = 792bp
#3
#4
#5
#6
#7
#8
#9
#10
#11
#12
#1
#2
LHA RHA
LHA RHA
LHA RHA
LHA RHA
LHA RHA
LHA RHA
GeneRuler 100bp DNA Ladder
LHA RHA
LHA RHA
LHA RHA
LHA RHA
LHA RHA
LHA RHA
1000 bp
750 bp
500 bp
250 bp
#15
#16
#17
#18
#19
#20
#21
#22
10D
H20
#13
#14
LHA RHA
LHA RHA
LHA RHA
LHA RHA
LHA RHA
LHA RHA
LHA RHA
LHA RHA
LHA RHA
LHA RHA
LHA RHA
LHA RHA
1000 bp
750 bp
500 bp
250 bp

### Slide 32
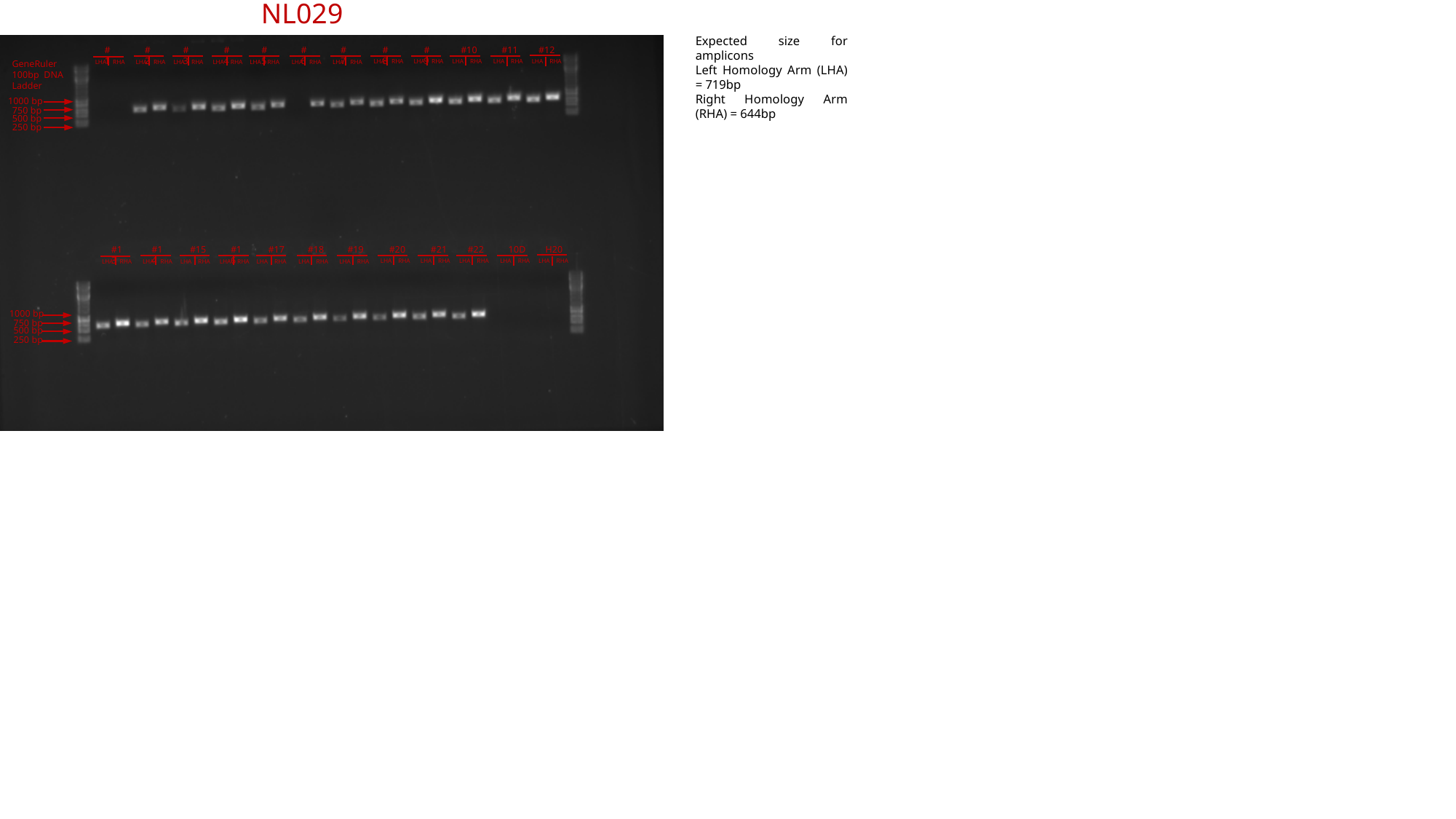

NL029
Expected size for amplicons
Left Homology Arm (LHA) = 719bp
Right Homology Arm (RHA) = 644bp
#3
#4
#5
#6
#7
#8
#9
#10
#11
#12
#1
#2
LHA RHA
LHA RHA
LHA RHA
LHA RHA
LHA RHA
LHA RHA
GeneRuler 100bp DNA Ladder
LHA RHA
LHA RHA
LHA RHA
LHA RHA
LHA RHA
LHA RHA
1000 bp
750 bp
500 bp
250 bp
#15
#16
#17
#18
#19
#20
#21
#22
10D
H20
#13
#14
LHA RHA
LHA RHA
LHA RHA
LHA RHA
LHA RHA
LHA RHA
LHA RHA
LHA RHA
LHA RHA
LHA RHA
LHA RHA
LHA RHA
1000 bp
750 bp
500 bp
250 bp

### Slide 33
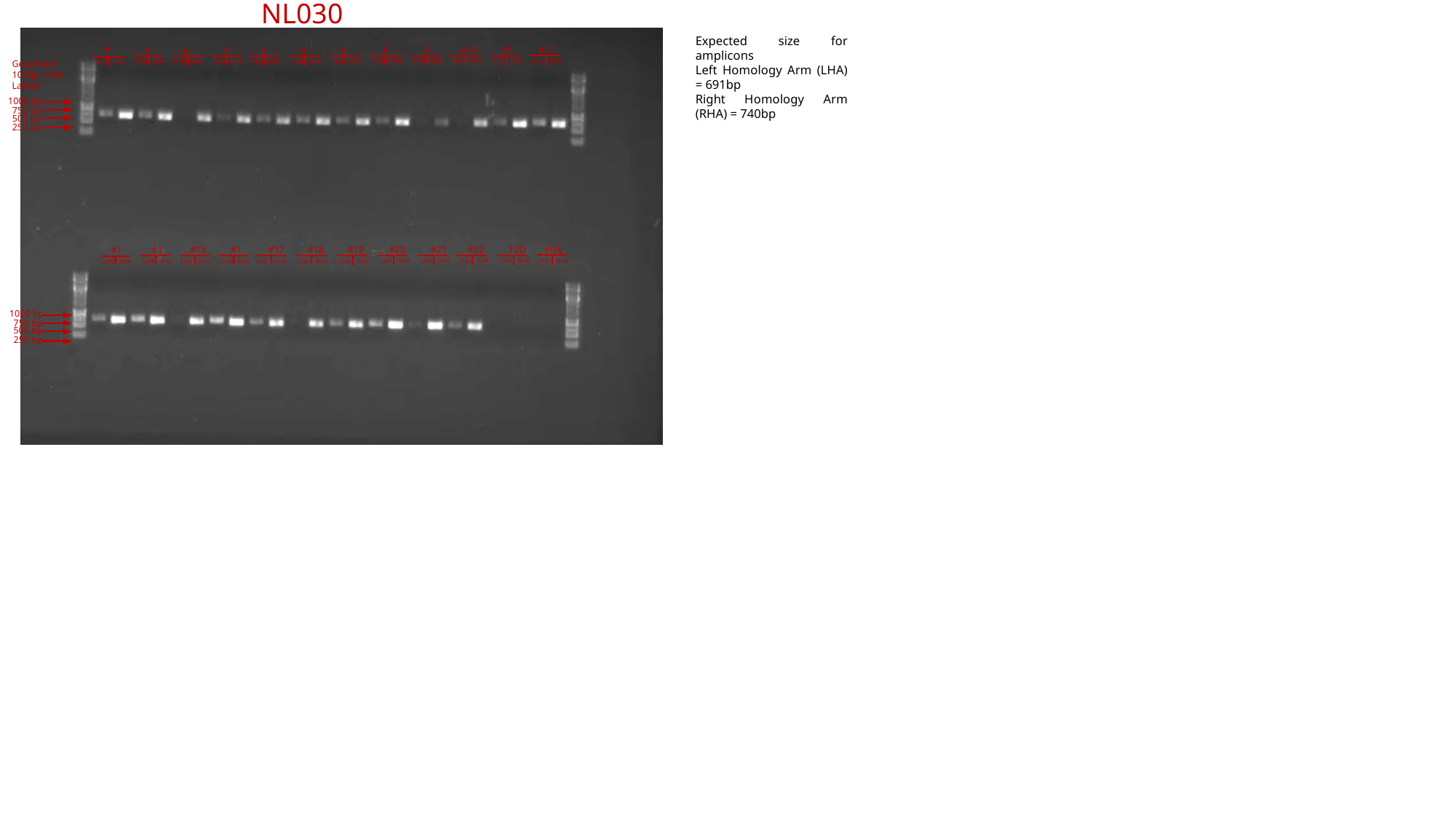

NL030
Expected size for amplicons
Left Homology Arm (LHA) = 691bp
Right Homology Arm (RHA) = 740bp
#3
#4
#5
#6
#7
#8
#9
#10
#11
#12
#1
#2
LHA RHA
LHA RHA
LHA RHA
LHA RHA
LHA RHA
LHA RHA
GeneRuler 100bp DNA Ladder
LHA RHA
LHA RHA
LHA RHA
LHA RHA
LHA RHA
LHA RHA
1000 bp
750 bp
500 bp
250 bp
#15
#16
#17
#18
#19
#20
#21
#22
10D
H20
#13
#14
LHA RHA
LHA RHA
LHA RHA
LHA RHA
LHA RHA
LHA RHA
LHA RHA
LHA RHA
LHA RHA
LHA RHA
LHA RHA
LHA RHA
1000 bp
750 bp
500 bp
250 bp

### Slide 34
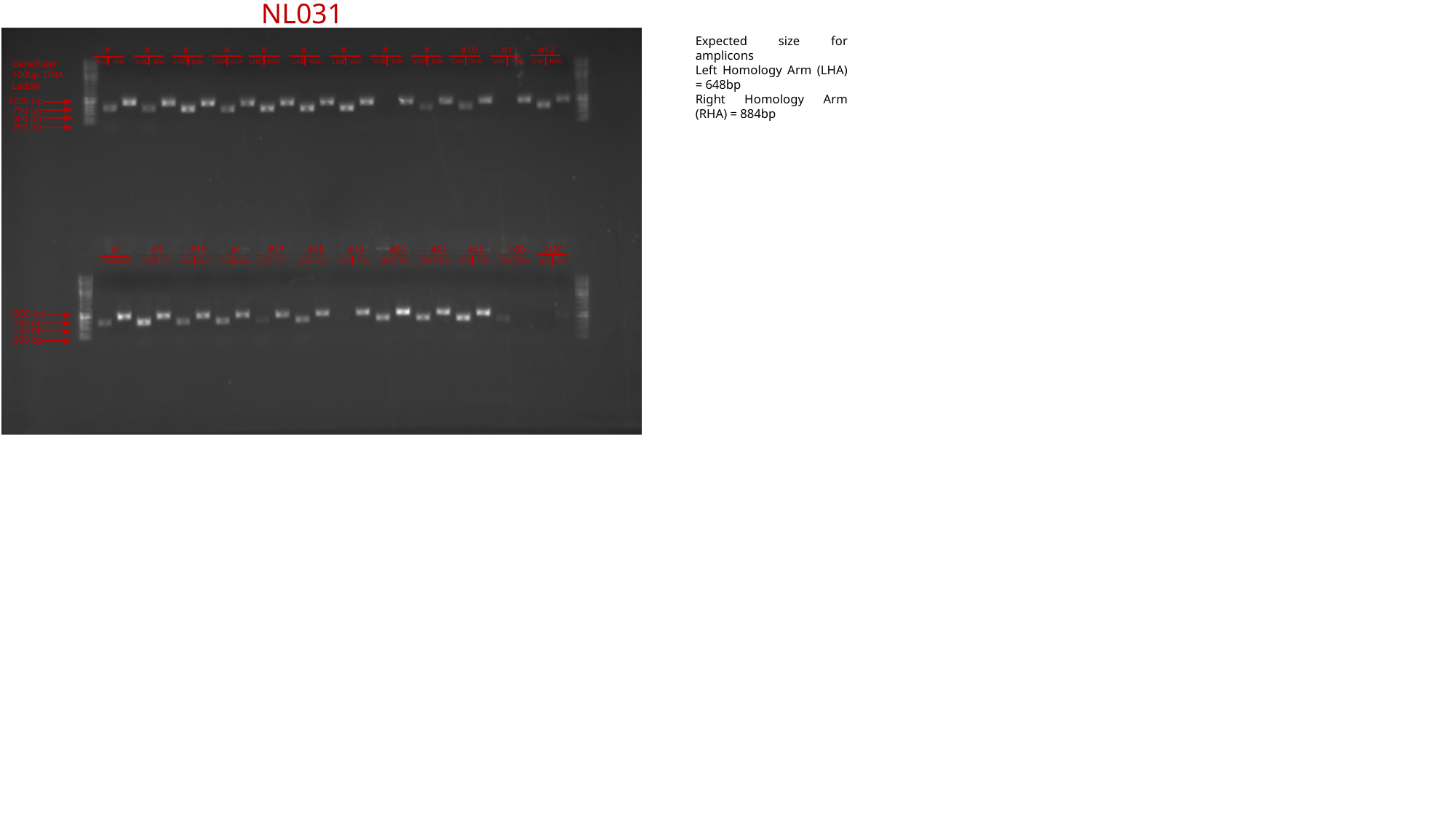

NL031
Expected size for amplicons
Left Homology Arm (LHA) = 648bp
Right Homology Arm (RHA) = 884bp
#3
#4
#5
#6
#7
#8
#9
#10
#11
#12
#1
#2
LHA RHA
LHA RHA
LHA RHA
LHA RHA
LHA RHA
LHA RHA
GeneRuler 100bp DNA Ladder
LHA RHA
LHA RHA
LHA RHA
LHA RHA
LHA RHA
LHA RHA
1000 bp
750 bp
500 bp
250 bp
#15
#16
#17
#18
#19
#20
#21
#22
10D
H20
#13
#14
LHA RHA
LHA RHA
LHA RHA
LHA RHA
LHA RHA
LHA RHA
LHA RHA
LHA RHA
LHA RHA
LHA RHA
LHA RHA
LHA RHA
1000 bp
750 bp
500 bp
250 bp

### Slide 35
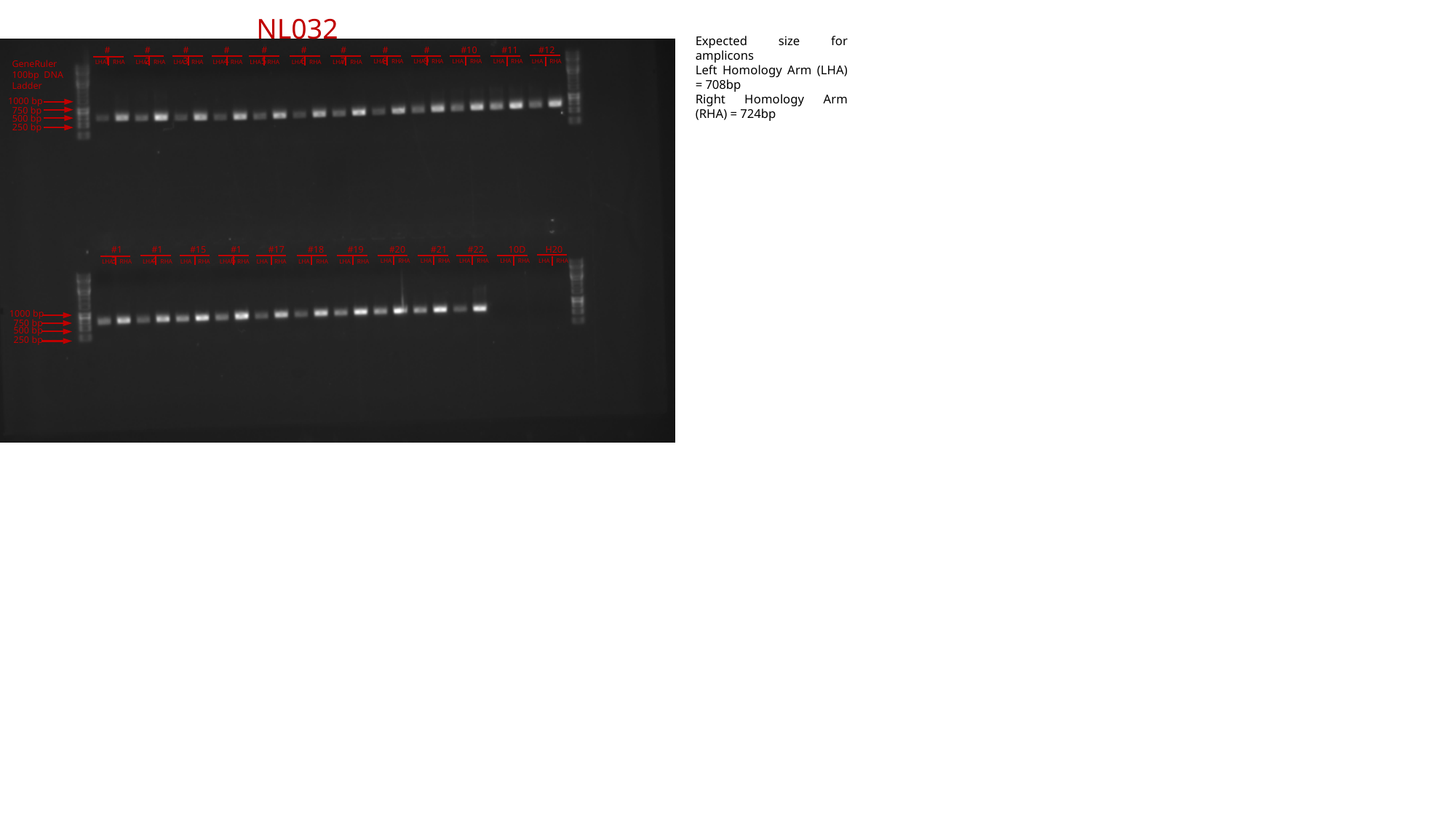

NL032
Expected size for amplicons
Left Homology Arm (LHA) = 708bp
Right Homology Arm (RHA) = 724bp
#3
#4
#5
#6
#7
#8
#9
#10
#11
#12
#1
#2
LHA RHA
LHA RHA
LHA RHA
LHA RHA
LHA RHA
LHA RHA
GeneRuler 100bp DNA Ladder
LHA RHA
LHA RHA
LHA RHA
LHA RHA
LHA RHA
LHA RHA
1000 bp
750 bp
500 bp
250 bp
#15
#16
#17
#18
#19
#20
#21
#22
10D
H20
#13
#14
LHA RHA
LHA RHA
LHA RHA
LHA RHA
LHA RHA
LHA RHA
LHA RHA
LHA RHA
LHA RHA
LHA RHA
LHA RHA
LHA RHA
1000 bp
750 bp
500 bp
250 bp

### Slide 36
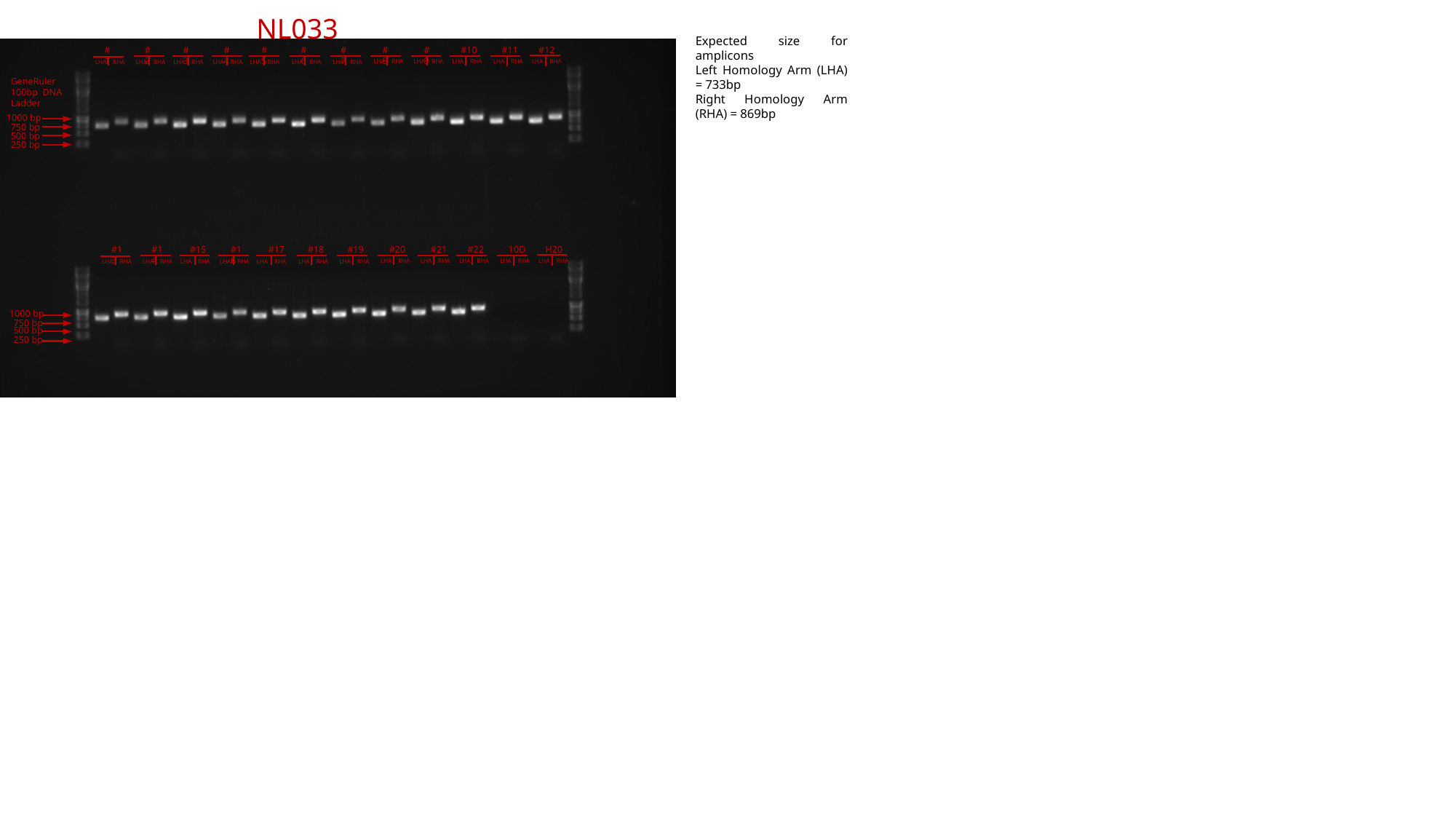

NL033
Expected size for amplicons
Left Homology Arm (LHA) = 733bp
Right Homology Arm (RHA) = 869bp
#3
#4
#5
#6
#7
#8
#9
#10
#11
#12
#1
#2
LHA RHA
LHA RHA
LHA RHA
LHA RHA
LHA RHA
LHA RHA
LHA RHA
LHA RHA
LHA RHA
LHA RHA
LHA RHA
LHA RHA
GeneRuler 100bp DNA Ladder
1000 bp
750 bp
500 bp
250 bp
#15
#16
#17
#18
#19
#20
#21
#22
10D
H20
#13
#14
LHA RHA
LHA RHA
LHA RHA
LHA RHA
LHA RHA
LHA RHA
LHA RHA
LHA RHA
LHA RHA
LHA RHA
LHA RHA
LHA RHA
1000 bp
750 bp
500 bp
250 bp

### Slide 37
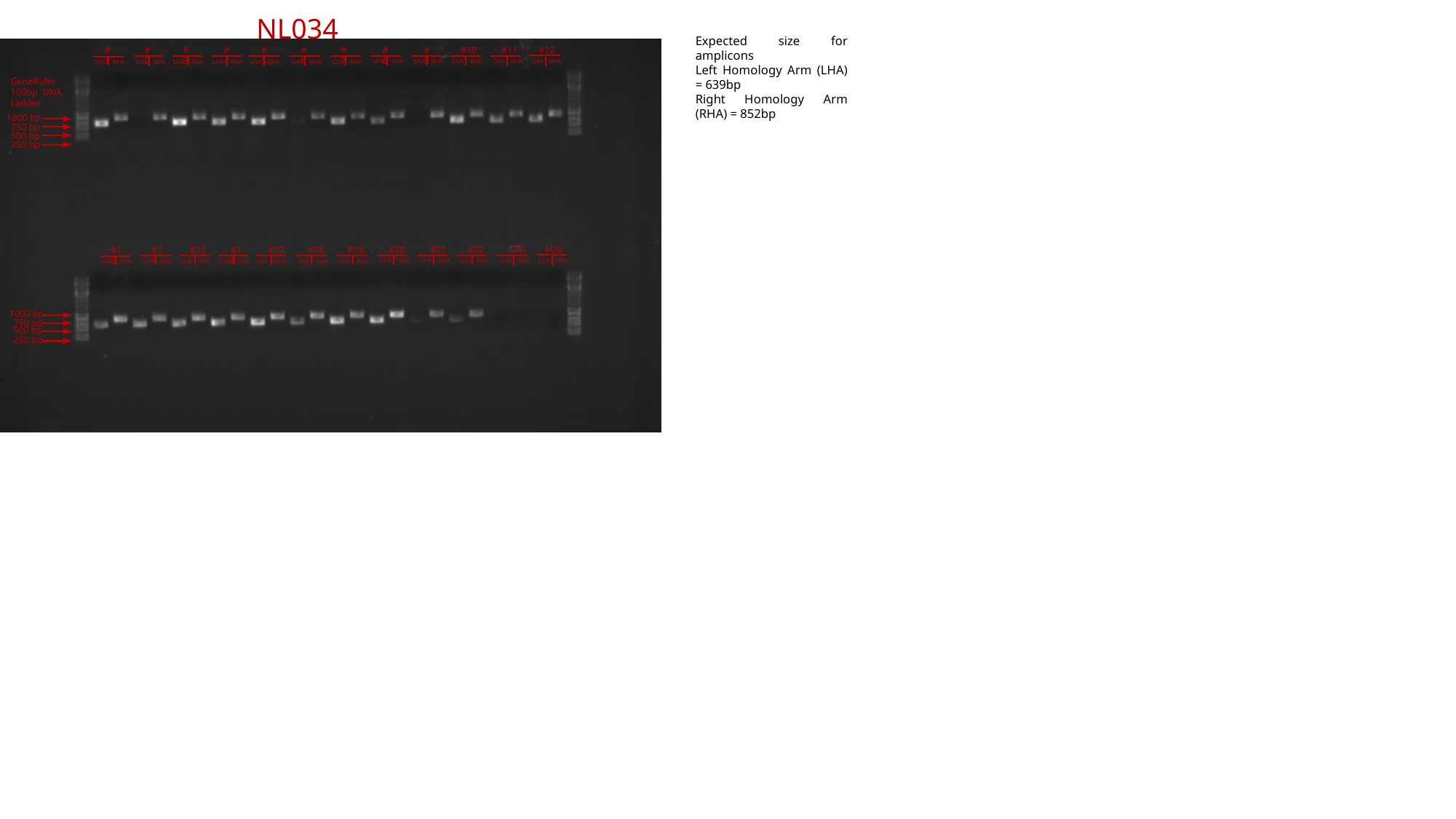

NL034
Expected size for amplicons
Left Homology Arm (LHA) = 639bp
Right Homology Arm (RHA) = 852bp
#3
#4
#5
#6
#7
#8
#9
#10
#11
#12
#1
#2
LHA RHA
LHA RHA
LHA RHA
LHA RHA
LHA RHA
LHA RHA
LHA RHA
LHA RHA
LHA RHA
LHA RHA
LHA RHA
LHA RHA
GeneRuler 100bp DNA Ladder
1000 bp
750 bp
500 bp
250 bp
#15
#16
#17
#18
#19
#20
#21
#22
10D
H20
#13
#14
LHA RHA
LHA RHA
LHA RHA
LHA RHA
LHA RHA
LHA RHA
LHA RHA
LHA RHA
LHA RHA
LHA RHA
LHA RHA
LHA RHA
1000 bp
750 bp
500 bp
250 bp

### Slide 38
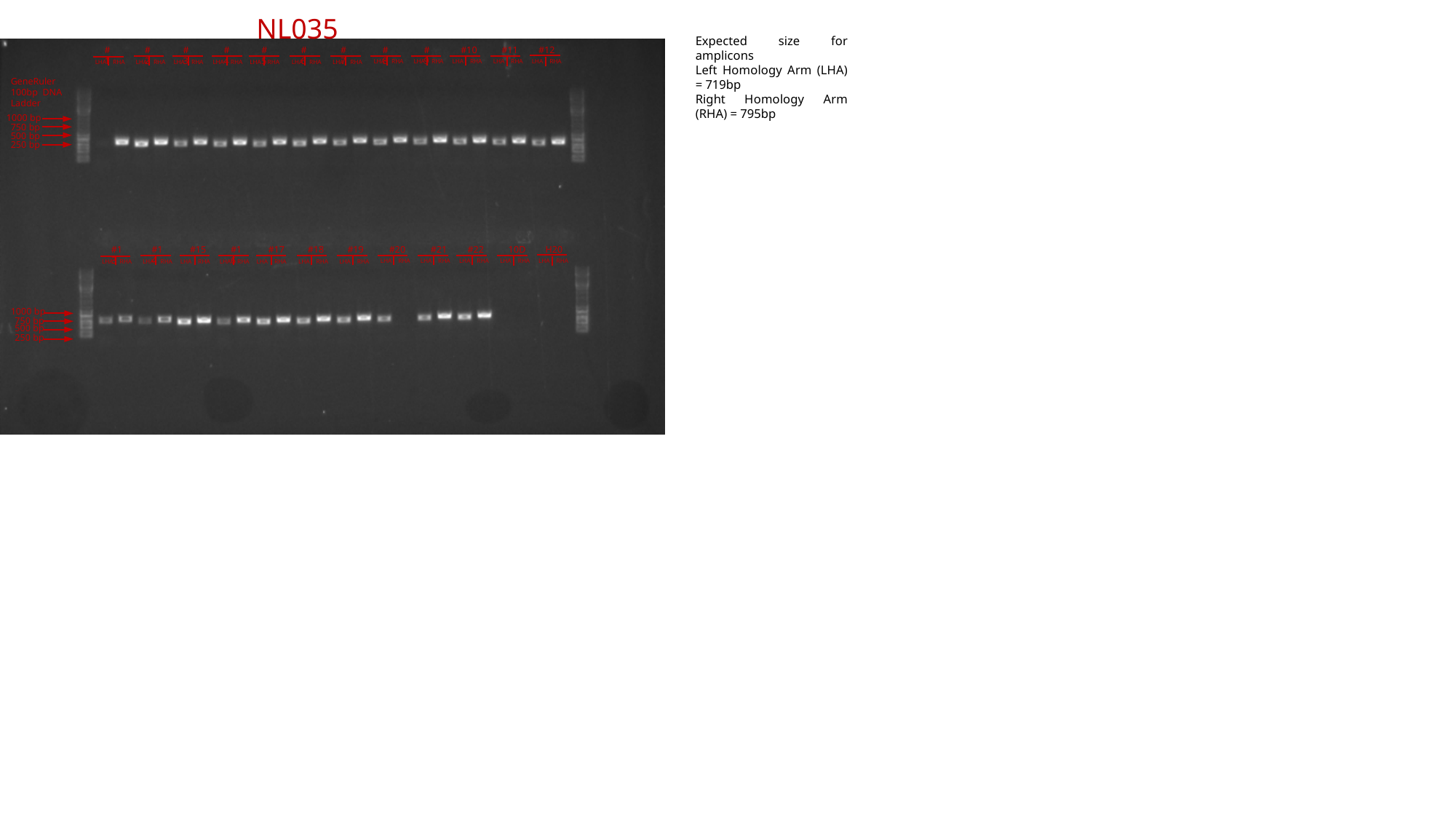

NL035
Expected size for amplicons
Left Homology Arm (LHA) = 719bp
Right Homology Arm (RHA) = 795bp
#3
#4
#5
#6
#7
#8
#9
#10
#11
#12
#1
#2
LHA RHA
LHA RHA
LHA RHA
LHA RHA
LHA RHA
LHA RHA
LHA RHA
LHA RHA
LHA RHA
LHA RHA
LHA RHA
LHA RHA
GeneRuler 100bp DNA Ladder
1000 bp
750 bp
500 bp
250 bp
#15
#16
#17
#18
#19
#20
#21
#22
10D
H20
#13
#14
LHA RHA
LHA RHA
LHA RHA
LHA RHA
LHA RHA
LHA RHA
LHA RHA
LHA RHA
LHA RHA
LHA RHA
LHA RHA
LHA RHA
1000 bp
750 bp
500 bp
250 bp

### Slide 39
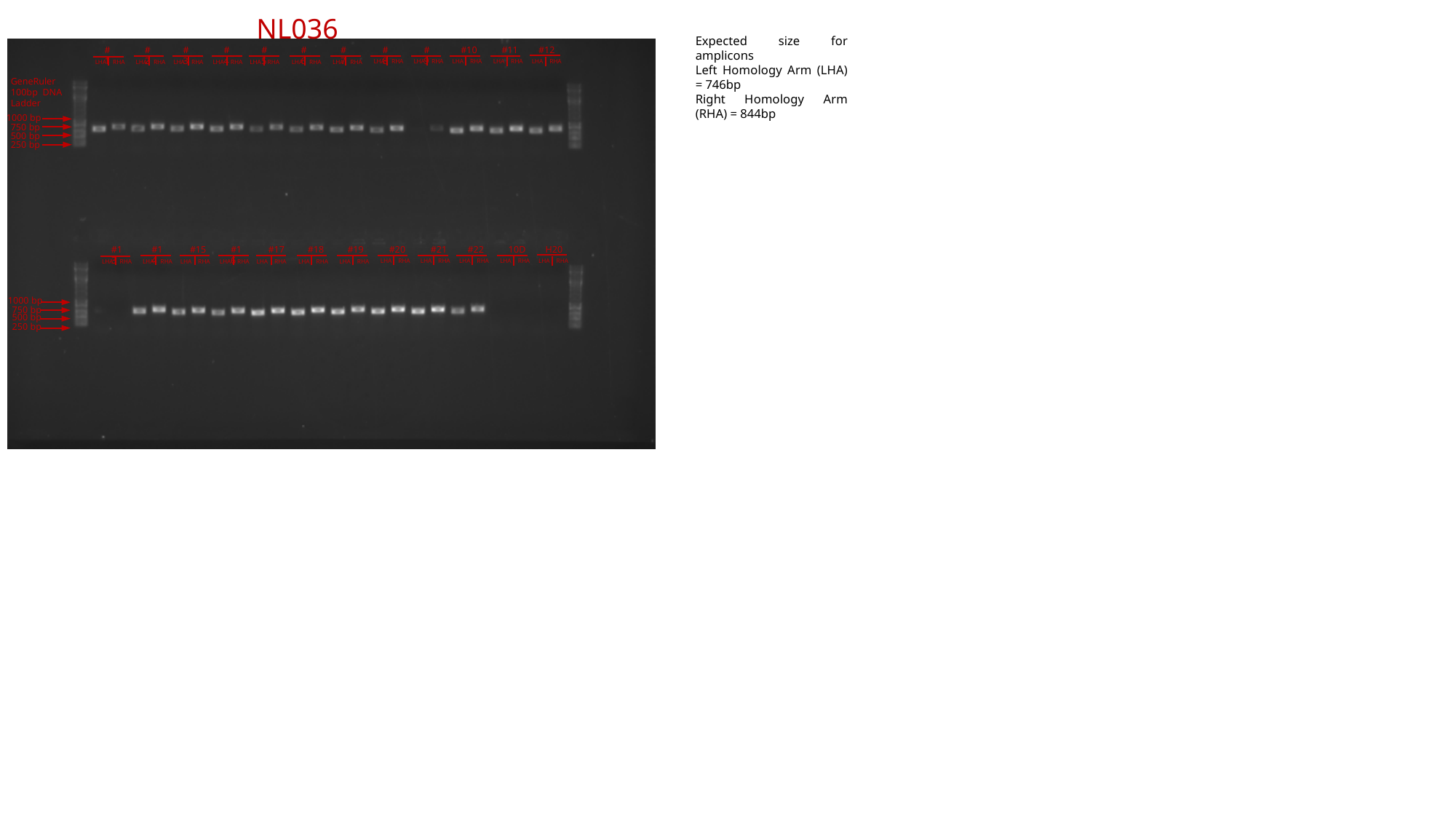

NL036
Expected size for amplicons
Left Homology Arm (LHA) = 746bp
Right Homology Arm (RHA) = 844bp
#3
#4
#5
#6
#7
#8
#9
#10
#11
#12
#1
#2
LHA RHA
LHA RHA
LHA RHA
LHA RHA
LHA RHA
LHA RHA
LHA RHA
LHA RHA
LHA RHA
LHA RHA
LHA RHA
LHA RHA
GeneRuler 100bp DNA Ladder
1000 bp
750 bp
500 bp
250 bp
#15
#16
#17
#18
#19
#20
#21
#22
10D
H20
#13
#14
LHA RHA
LHA RHA
LHA RHA
LHA RHA
LHA RHA
LHA RHA
LHA RHA
LHA RHA
LHA RHA
LHA RHA
LHA RHA
LHA RHA
1000 bp
750 bp
500 bp
250 bp

### Slide 40
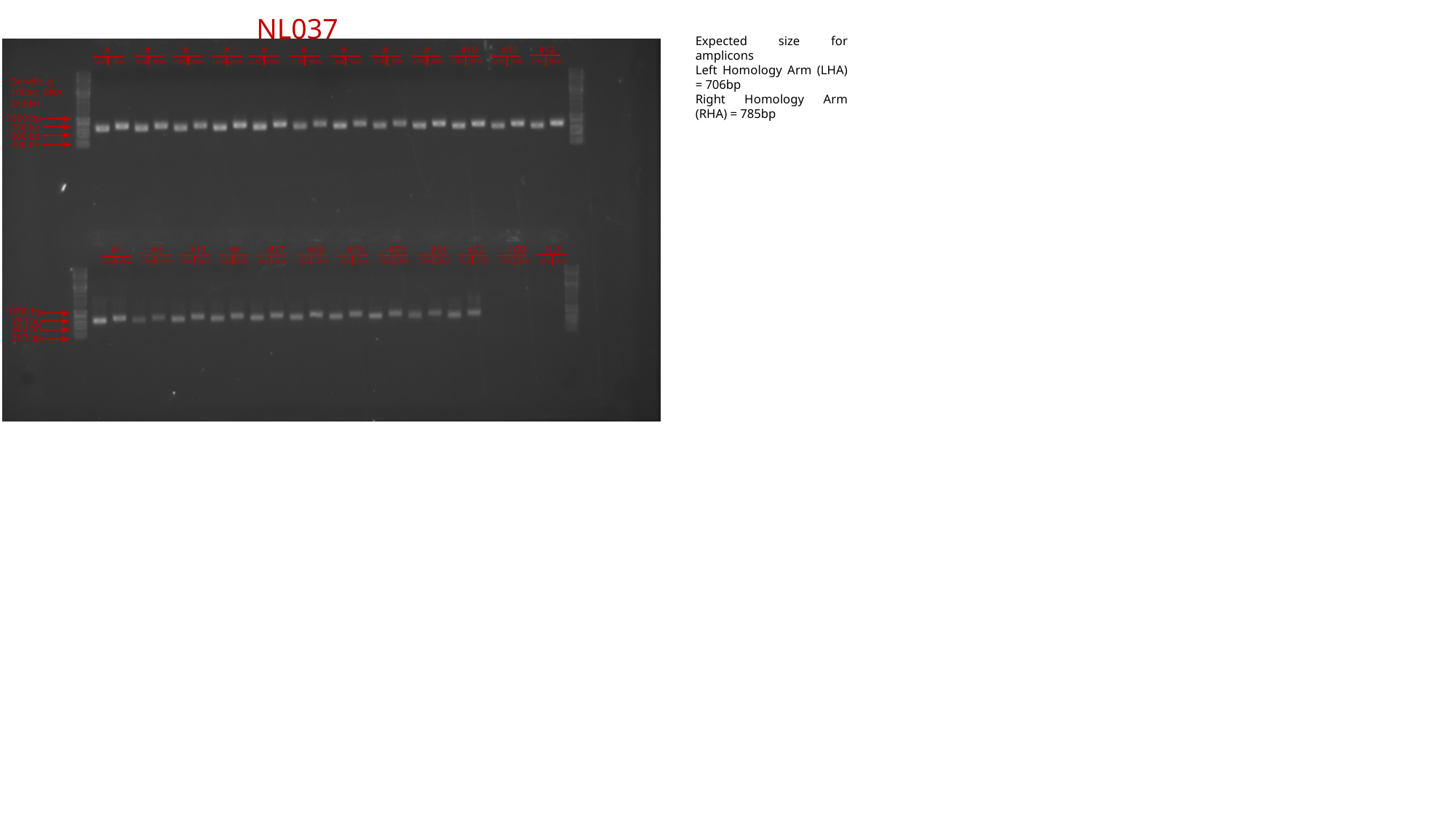

NL037
Expected size for amplicons
Left Homology Arm (LHA) = 706bp
Right Homology Arm (RHA) = 785bp
#3
#4
#5
#6
#7
#8
#9
#10
#11
#12
#1
#2
LHA RHA
LHA RHA
LHA RHA
LHA RHA
LHA RHA
LHA RHA
LHA RHA
LHA RHA
LHA RHA
LHA RHA
LHA RHA
LHA RHA
GeneRuler 100bp DNA Ladder
1000 bp
750 bp
500 bp
250 bp
#15
#16
#17
#18
#19
#20
#21
#22
10D
H20
#13
#14
LHA RHA
LHA RHA
LHA RHA
LHA RHA
LHA RHA
LHA RHA
LHA RHA
LHA RHA
LHA RHA
LHA RHA
LHA RHA
LHA RHA
1000 bp
750 bp
500 bp
250 bp

### Slide 41
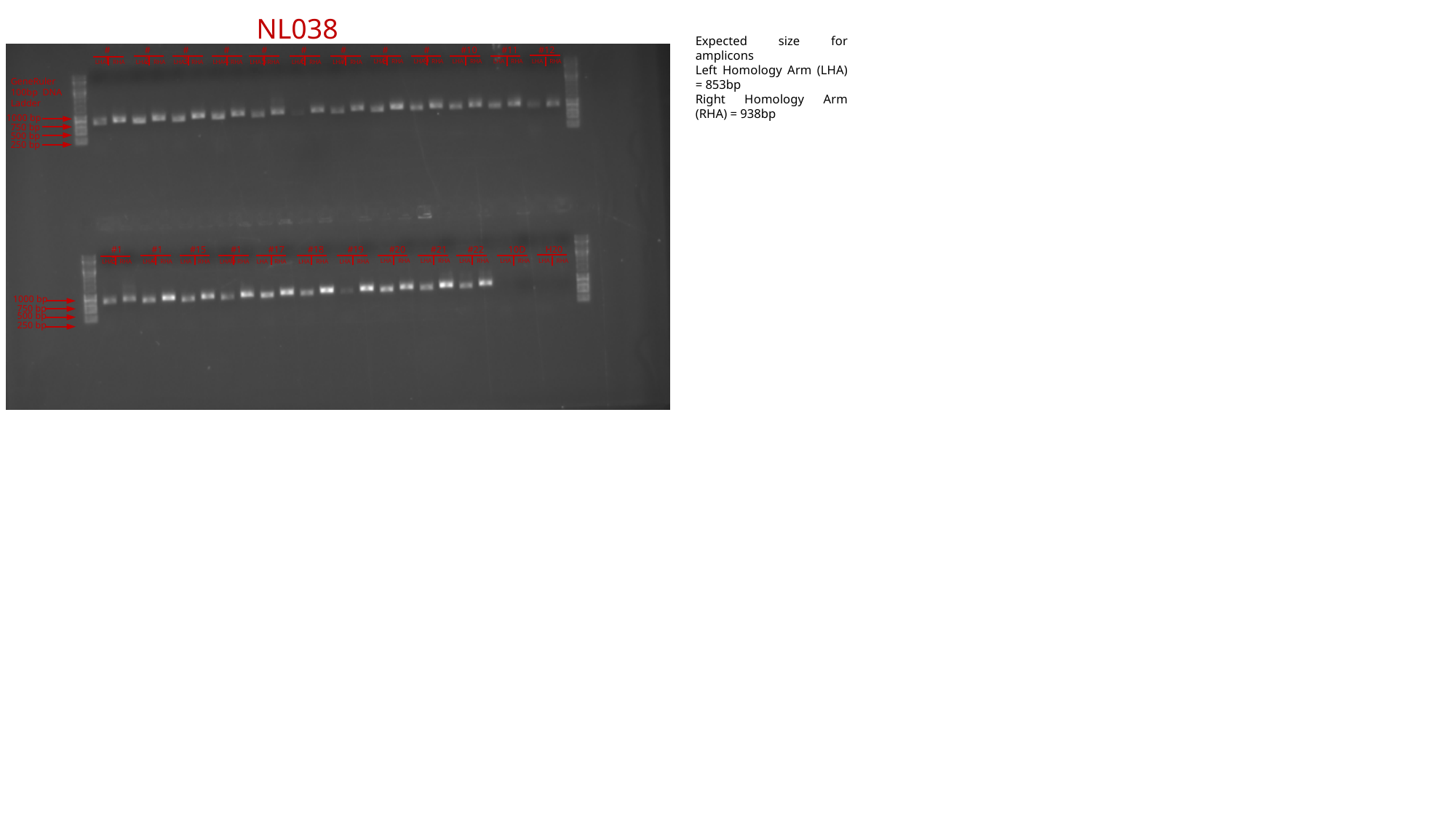

NL038
Expected size for amplicons
Left Homology Arm (LHA) = 853bp
Right Homology Arm (RHA) = 938bp
#3
#4
#5
#6
#7
#8
#9
#10
#11
#12
#1
#2
LHA RHA
LHA RHA
LHA RHA
LHA RHA
LHA RHA
LHA RHA
LHA RHA
LHA RHA
LHA RHA
LHA RHA
LHA RHA
LHA RHA
GeneRuler 100bp DNA Ladder
1000 bp
750 bp
500 bp
250 bp
#15
#16
#17
#18
#19
#20
#21
#22
10D
H20
#13
#14
LHA RHA
LHA RHA
LHA RHA
LHA RHA
LHA RHA
LHA RHA
LHA RHA
LHA RHA
LHA RHA
LHA RHA
LHA RHA
LHA RHA
1000 bp
750 bp
500 bp
250 bp

### Slide 42
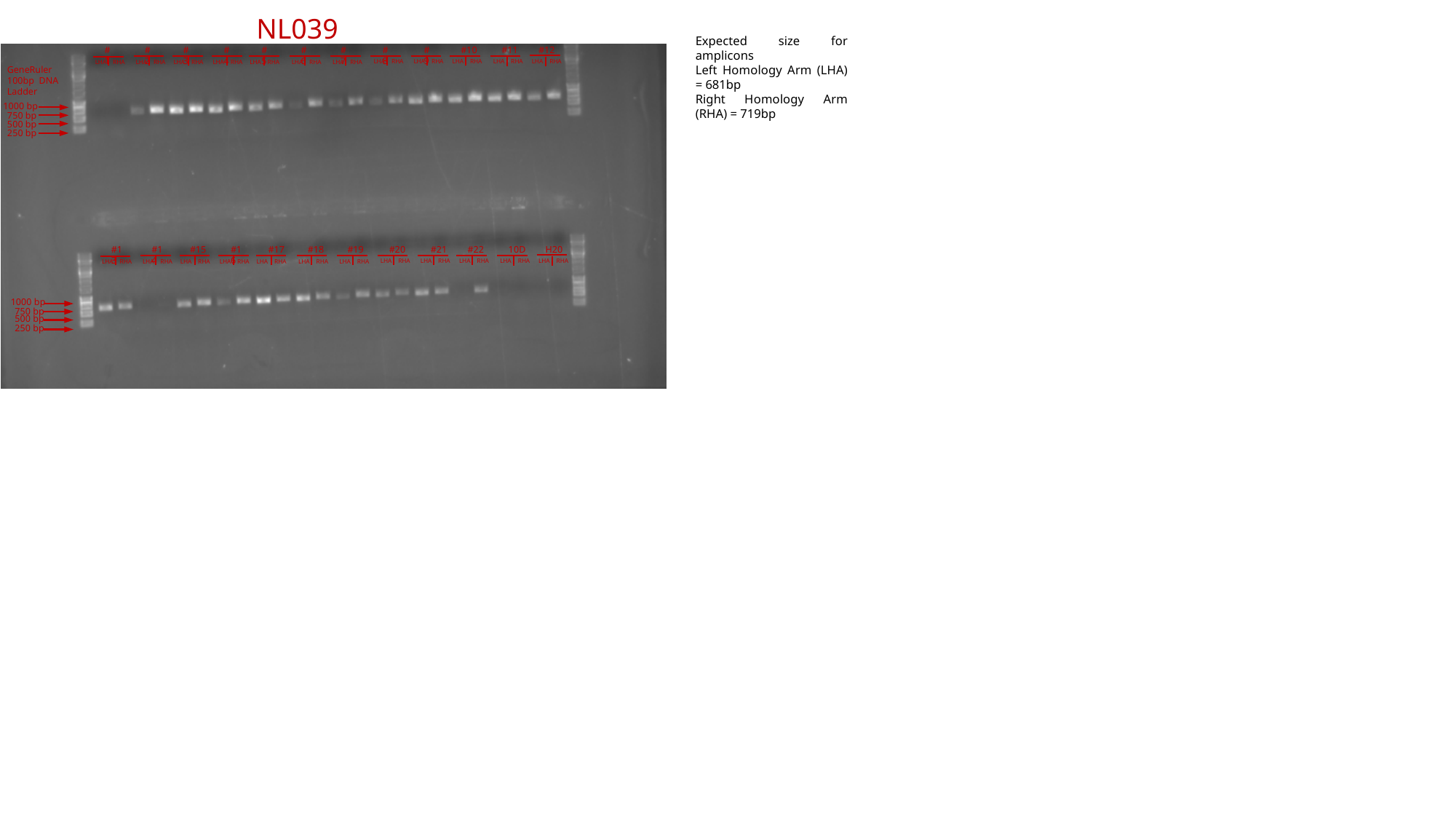

NL039
Expected size for amplicons
Left Homology Arm (LHA) = 681bp
Right Homology Arm (RHA) = 719bp
#3
#4
#5
#6
#7
#8
#9
#10
#11
#12
#1
#2
LHA RHA
LHA RHA
LHA RHA
LHA RHA
LHA RHA
LHA RHA
LHA RHA
LHA RHA
LHA RHA
LHA RHA
LHA RHA
LHA RHA
GeneRuler 100bp DNA Ladder
1000 bp
750 bp
500 bp
250 bp
#15
#16
#17
#18
#19
#20
#21
#22
10D
H20
#13
#14
LHA RHA
LHA RHA
LHA RHA
LHA RHA
LHA RHA
LHA RHA
LHA RHA
LHA RHA
LHA RHA
LHA RHA
LHA RHA
LHA RHA
1000 bp
750 bp
500 bp
250 bp

### Slide 43
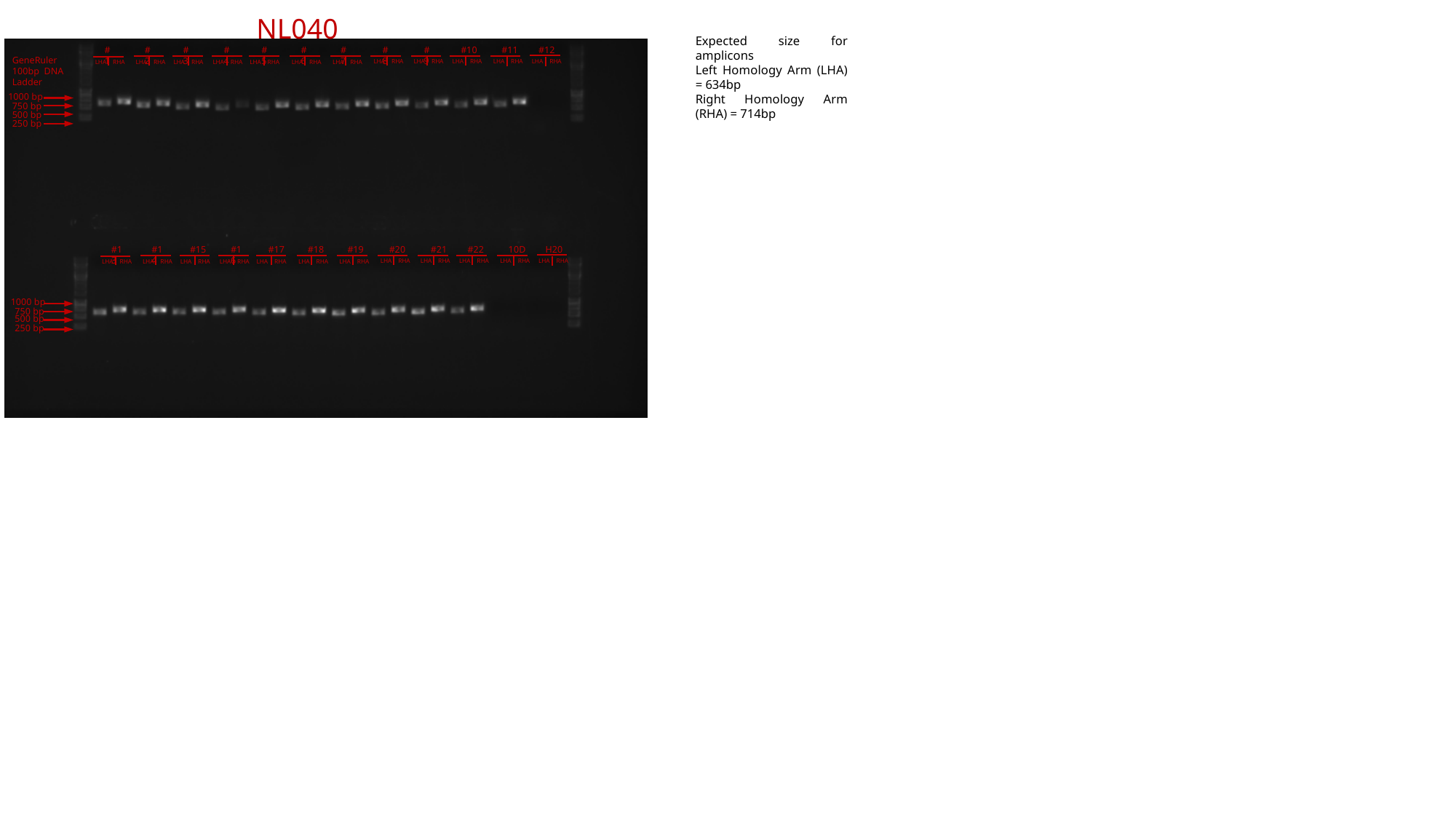

NL040
Expected size for amplicons
Left Homology Arm (LHA) = 634bp
Right Homology Arm (RHA) = 714bp
#3
#4
#5
#6
#7
#8
#9
#10
#11
#12
#1
#2
GeneRuler 100bp DNA Ladder
LHA RHA
LHA RHA
LHA RHA
LHA RHA
LHA RHA
LHA RHA
LHA RHA
LHA RHA
LHA RHA
LHA RHA
LHA RHA
LHA RHA
1000 bp
750 bp
500 bp
250 bp
#15
#16
#17
#18
#19
#20
#21
#22
10D
H20
#13
#14
LHA RHA
LHA RHA
LHA RHA
LHA RHA
LHA RHA
LHA RHA
LHA RHA
LHA RHA
LHA RHA
LHA RHA
LHA RHA
LHA RHA
1000 bp
750 bp
500 bp
250 bp

### Slide 44
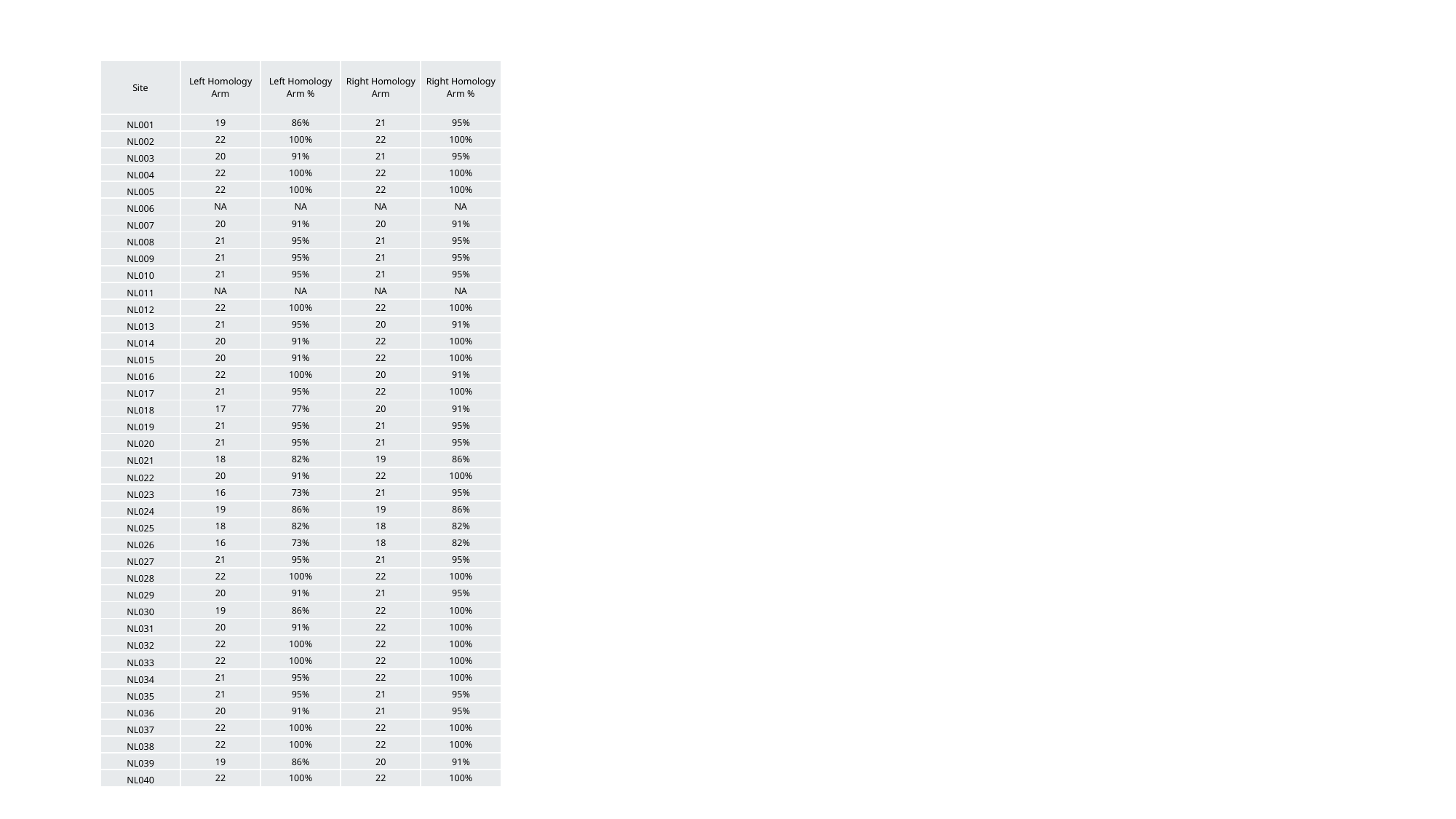

| Site | Left Homology Arm | Left Homology Arm % | Right Homology Arm | Right Homology Arm % |
| --- | --- | --- | --- | --- |
| NL001 | 19 | 86% | 21 | 95% |
| NL002 | 22 | 100% | 22 | 100% |
| NL003 | 20 | 91% | 21 | 95% |
| NL004 | 22 | 100% | 22 | 100% |
| NL005 | 22 | 100% | 22 | 100% |
| NL006 | NA | NA | NA | NA |
| NL007 | 20 | 91% | 20 | 91% |
| NL008 | 21 | 95% | 21 | 95% |
| NL009 | 21 | 95% | 21 | 95% |
| NL010 | 21 | 95% | 21 | 95% |
| NL011 | NA | NA | NA | NA |
| NL012 | 22 | 100% | 22 | 100% |
| NL013 | 21 | 95% | 20 | 91% |
| NL014 | 20 | 91% | 22 | 100% |
| NL015 | 20 | 91% | 22 | 100% |
| NL016 | 22 | 100% | 20 | 91% |
| NL017 | 21 | 95% | 22 | 100% |
| NL018 | 17 | 77% | 20 | 91% |
| NL019 | 21 | 95% | 21 | 95% |
| NL020 | 21 | 95% | 21 | 95% |
| NL021 | 18 | 82% | 19 | 86% |
| NL022 | 20 | 91% | 22 | 100% |
| NL023 | 16 | 73% | 21 | 95% |
| NL024 | 19 | 86% | 19 | 86% |
| NL025 | 18 | 82% | 18 | 82% |
| NL026 | 16 | 73% | 18 | 82% |
| NL027 | 21 | 95% | 21 | 95% |
| NL028 | 22 | 100% | 22 | 100% |
| NL029 | 20 | 91% | 21 | 95% |
| NL030 | 19 | 86% | 22 | 100% |
| NL031 | 20 | 91% | 22 | 100% |
| NL032 | 22 | 100% | 22 | 100% |
| NL033 | 22 | 100% | 22 | 100% |
| NL034 | 21 | 95% | 22 | 100% |
| NL035 | 21 | 95% | 21 | 95% |
| NL036 | 20 | 91% | 21 | 95% |
| NL037 | 22 | 100% | 22 | 100% |
| NL038 | 22 | 100% | 22 | 100% |
| NL039 | 19 | 86% | 20 | 91% |
| NL040 | 22 | 100% | 22 | 100% |

### Slide 45
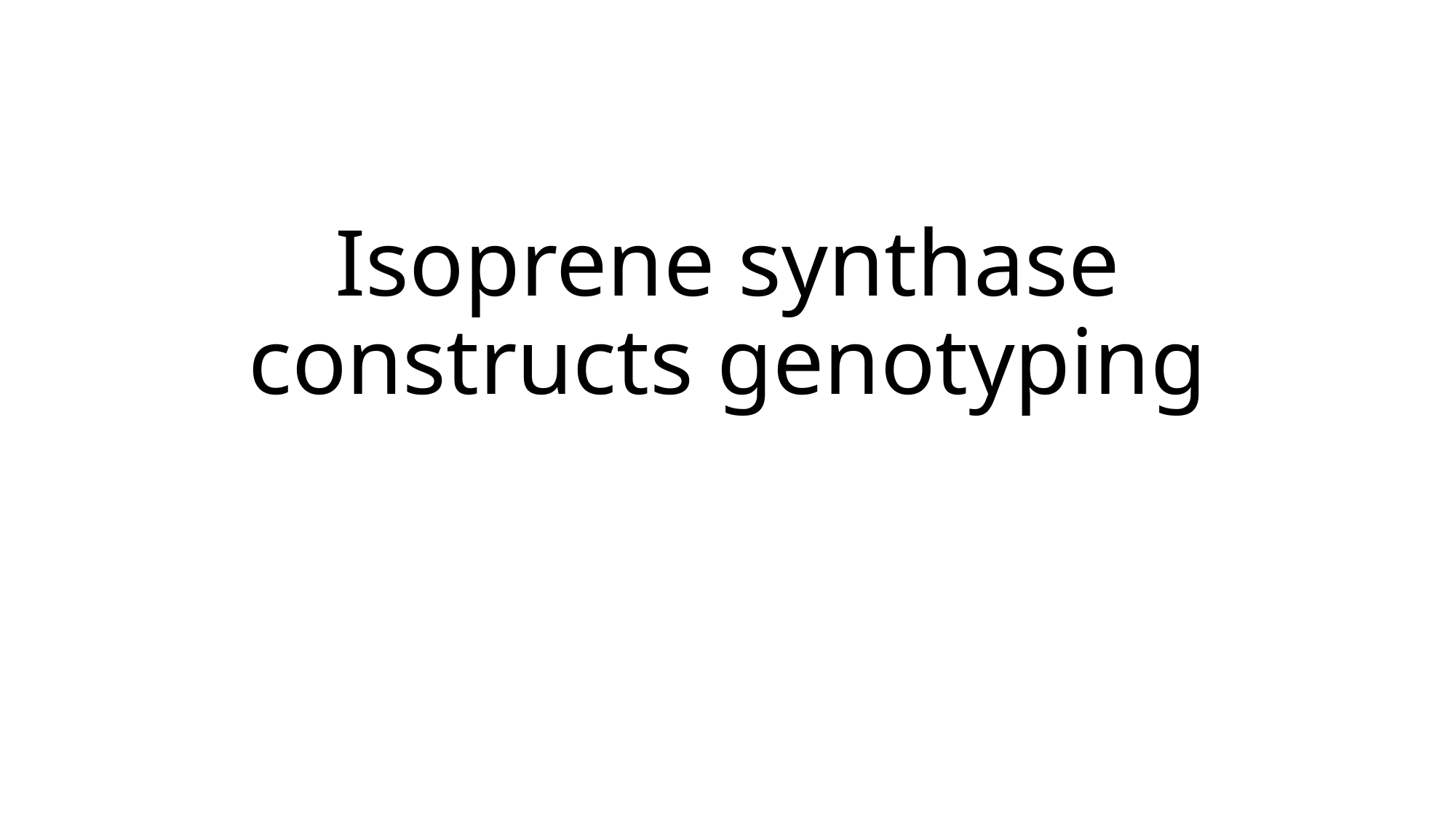

#### Isoprene synthase constructs genotyping

### Slide 46
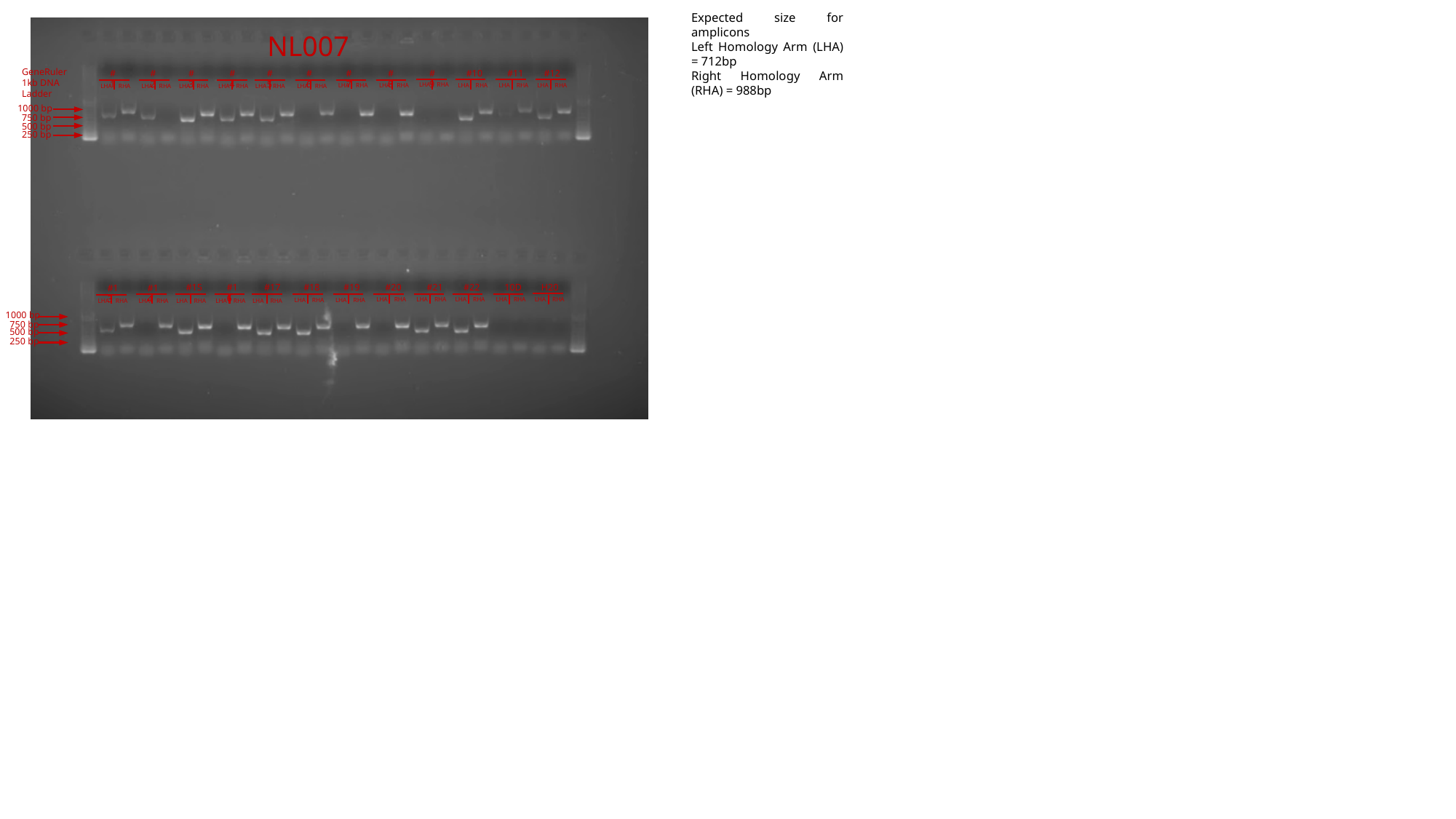

Expected size for amplicons
Left Homology Arm (LHA) = 712bp
Right Homology Arm (RHA) = 988bp
NL007
GeneRuler 1kb DNA Ladder
#3
#4
#5
#6
#7
#8
#9
#10
#11
#12
#1
#2
LHA RHA
LHA RHA
LHA RHA
LHA RHA
LHA RHA
LHA RHA
LHA RHA
LHA RHA
LHA RHA
LHA RHA
LHA RHA
LHA RHA
1000 bp
750 bp
500 bp
250 bp
#15
#16
#17
#18
#19
#20
#21
#22
10D
H20
#13
#14
LHA RHA
LHA RHA
LHA RHA
LHA RHA
LHA RHA
LHA RHA
LHA RHA
LHA RHA
LHA RHA
LHA RHA
LHA RHA
LHA RHA
1000 bp
750 bp
500 bp
250 bp

### Slide 47
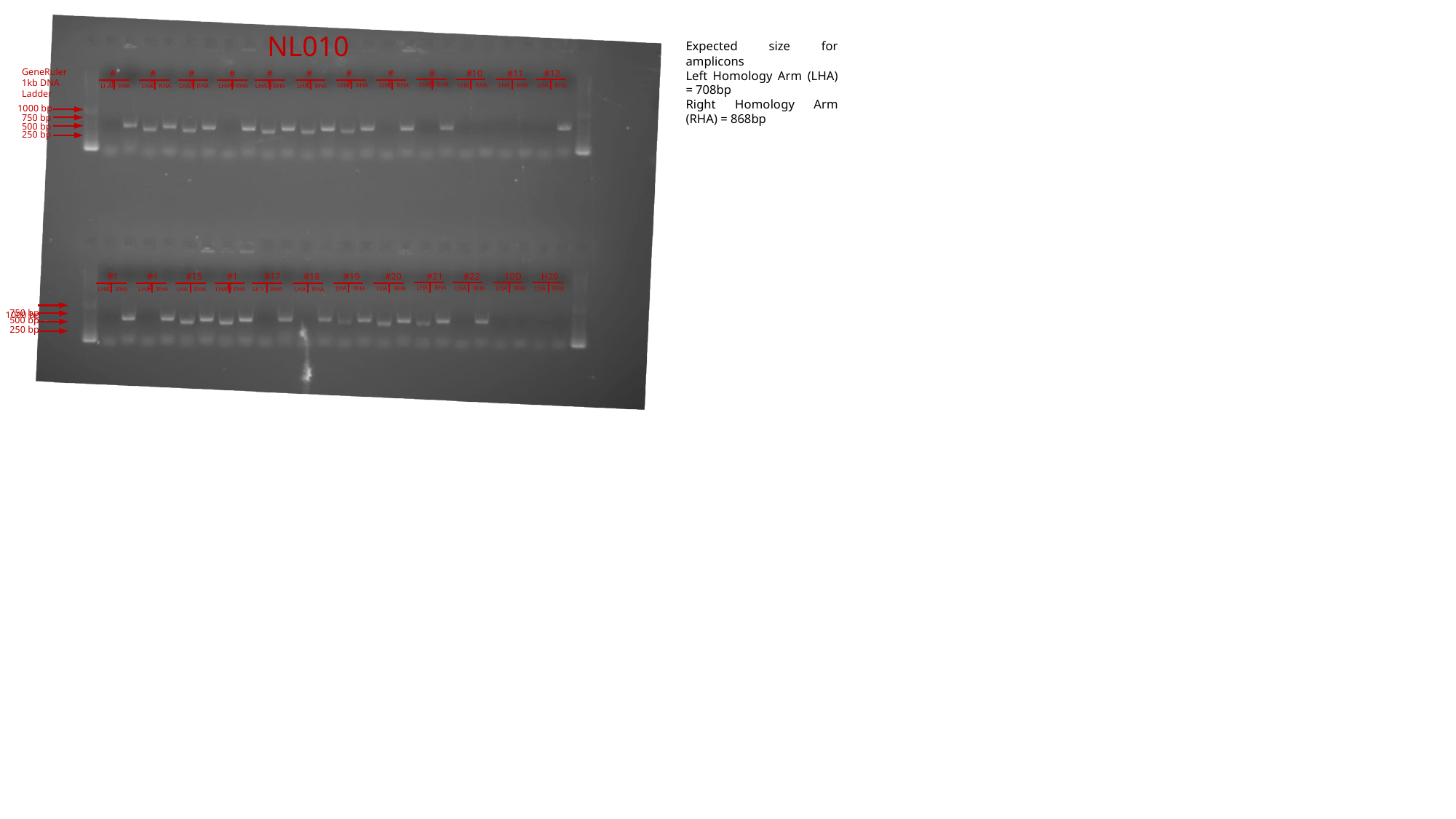

NL010
Expected size for amplicons
Left Homology Arm (LHA) = 708bp
Right Homology Arm (RHA) = 868bp
GeneRuler 1kb DNA Ladder
#3
#4
#5
#6
#7
#8
#9
#10
#11
#12
#1
#2
LHA RHA
LHA RHA
LHA RHA
LHA RHA
LHA RHA
LHA RHA
LHA RHA
LHA RHA
LHA RHA
LHA RHA
LHA RHA
LHA RHA
1000 bp
750 bp
500 bp
250 bp
#15
#16
#17
#18
#19
#20
#21
#22
10D
H20
#13
#14
LHA RHA
LHA RHA
LHA RHA
LHA RHA
LHA RHA
LHA RHA
LHA RHA
LHA RHA
LHA RHA
LHA RHA
LHA RHA
LHA RHA
750 bp
1000 bp
500 bp
250 bp

### Slide 48
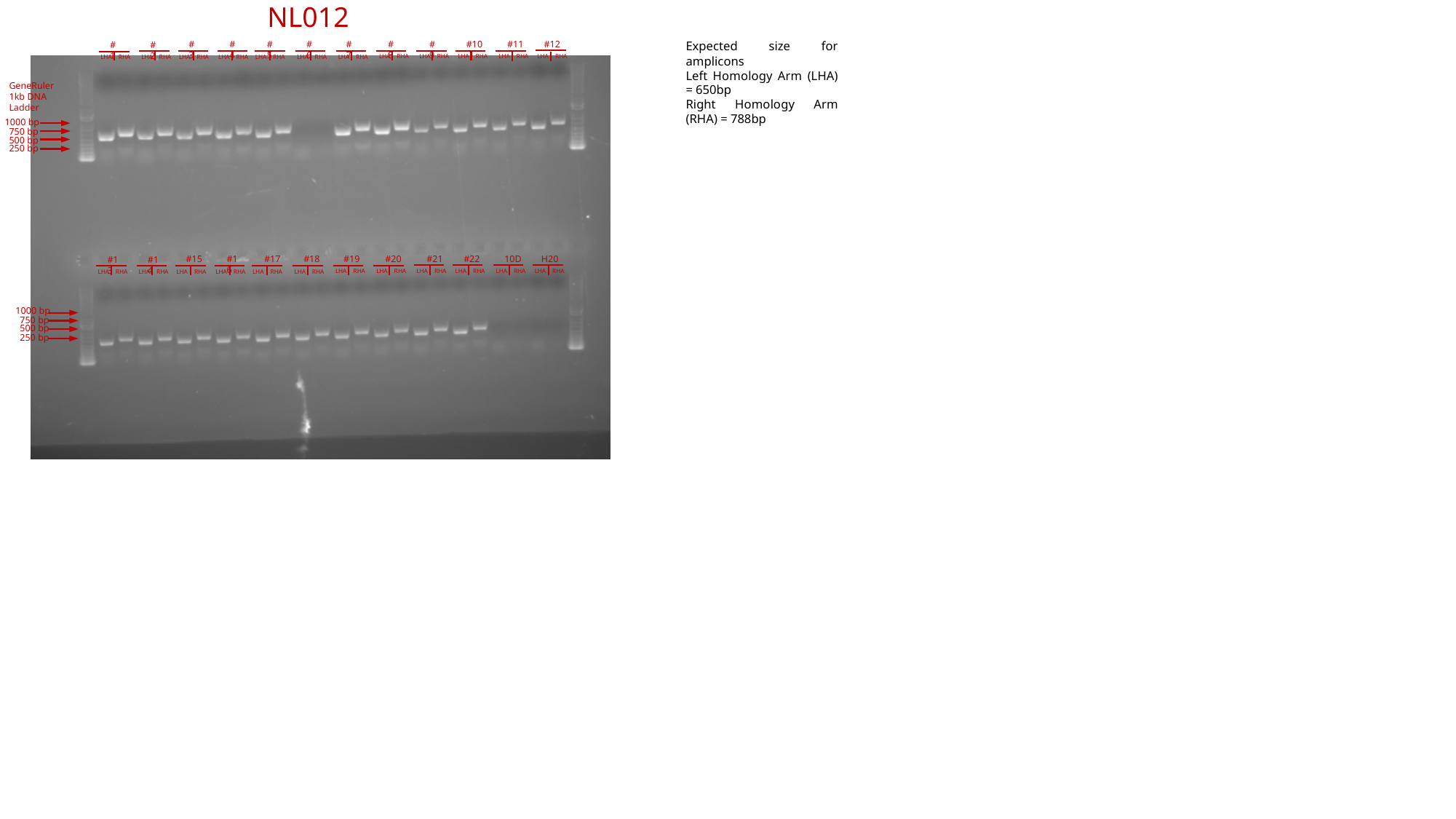

NL012
#3
#4
#5
#6
#7
#8
#9
#10
#11
#12
Expected size for amplicons
Left Homology Arm (LHA) = 650bp
Right Homology Arm (RHA) = 788bp
#1
#2
LHA RHA
LHA RHA
LHA RHA
LHA RHA
LHA RHA
LHA RHA
LHA RHA
LHA RHA
LHA RHA
LHA RHA
LHA RHA
LHA RHA
GeneRuler 1kb DNA Ladder
1000 bp
750 bp
500 bp
250 bp
#15
#16
#17
#18
#19
#20
#21
#22
10D
H20
#13
#14
LHA RHA
LHA RHA
LHA RHA
LHA RHA
LHA RHA
LHA RHA
LHA RHA
LHA RHA
LHA RHA
LHA RHA
LHA RHA
LHA RHA
1000 bp
750 bp
500 bp
250 bp

### Slide 49
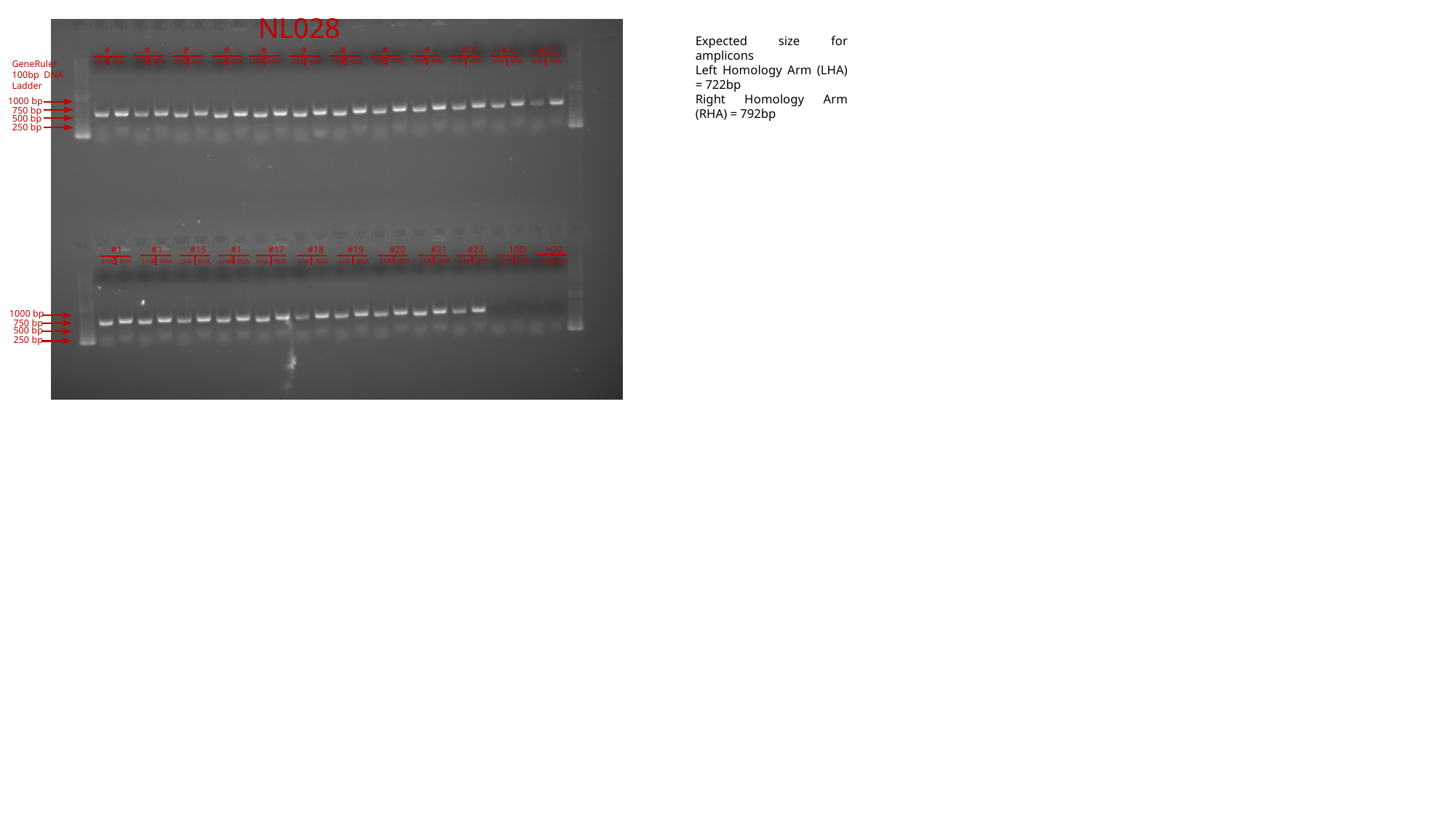

NL028
Expected size for amplicons
Left Homology Arm (LHA) = 722bp
Right Homology Arm (RHA) = 792bp
#3
#4
#5
#6
#7
#8
#9
#10
#11
#12
#1
#2
LHA RHA
LHA RHA
LHA RHA
LHA RHA
LHA RHA
LHA RHA
GeneRuler 100bp DNA Ladder
LHA RHA
LHA RHA
LHA RHA
LHA RHA
LHA RHA
LHA RHA
1000 bp
750 bp
500 bp
250 bp
#15
#16
#17
#18
#19
#20
#21
#22
10D
H20
#13
#14
LHA RHA
LHA RHA
LHA RHA
LHA RHA
LHA RHA
LHA RHA
LHA RHA
LHA RHA
LHA RHA
LHA RHA
LHA RHA
LHA RHA
1000 bp
750 bp
500 bp
250 bp

### Slide 50
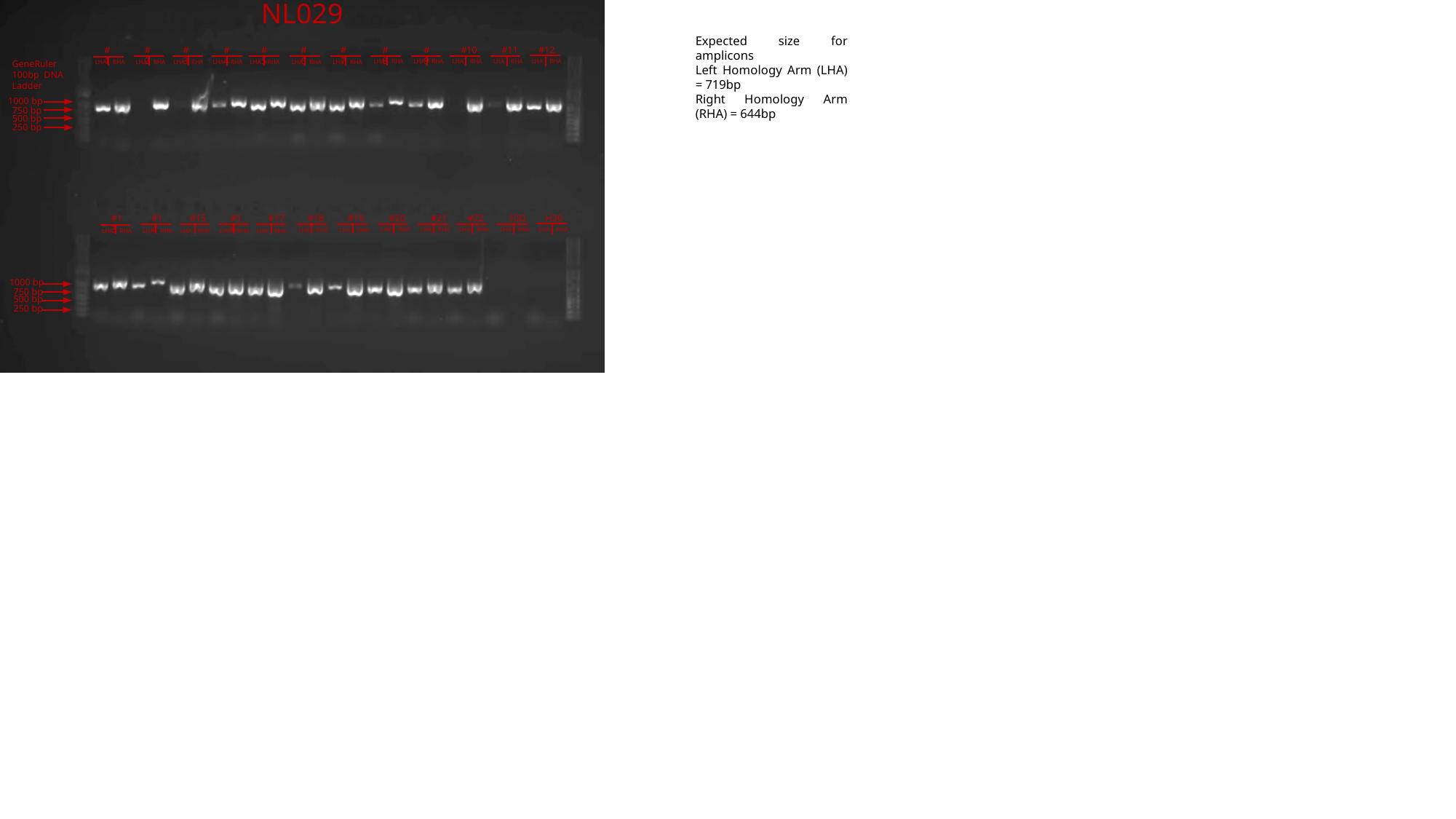

NL029
Expected size for amplicons
Left Homology Arm (LHA) = 719bp
Right Homology Arm (RHA) = 644bp
#3
#4
#5
#6
#7
#8
#9
#10
#11
#12
#1
#2
LHA RHA
LHA RHA
LHA RHA
LHA RHA
LHA RHA
LHA RHA
GeneRuler 100bp DNA Ladder
LHA RHA
LHA RHA
LHA RHA
LHA RHA
LHA RHA
LHA RHA
1000 bp
750 bp
500 bp
250 bp
#15
#16
#17
#18
#19
#20
#21
#22
10D
H20
#13
#14
LHA RHA
LHA RHA
LHA RHA
LHA RHA
LHA RHA
LHA RHA
LHA RHA
LHA RHA
LHA RHA
LHA RHA
LHA RHA
LHA RHA
1000 bp
750 bp
500 bp
250 bp

### Slide 51
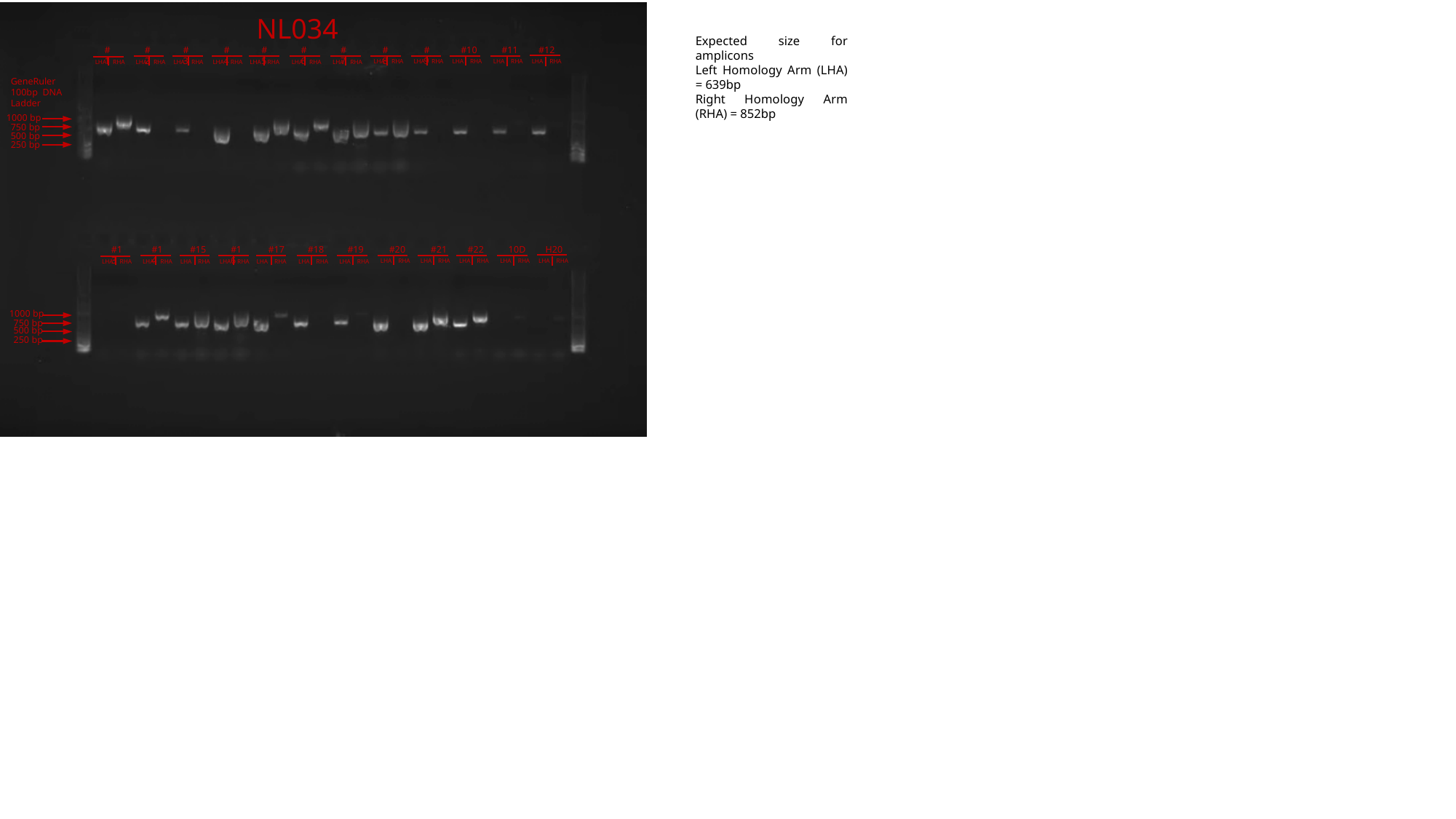

NL034
Expected size for amplicons
Left Homology Arm (LHA) = 639bp
Right Homology Arm (RHA) = 852bp
#3
#4
#5
#6
#7
#8
#9
#10
#11
#12
#1
#2
LHA RHA
LHA RHA
LHA RHA
LHA RHA
LHA RHA
LHA RHA
LHA RHA
LHA RHA
LHA RHA
LHA RHA
LHA RHA
LHA RHA
GeneRuler 100bp DNA Ladder
1000 bp
750 bp
500 bp
250 bp
#15
#16
#17
#18
#19
#20
#21
#22
10D
H20
#13
#14
LHA RHA
LHA RHA
LHA RHA
LHA RHA
LHA RHA
LHA RHA
LHA RHA
LHA RHA
LHA RHA
LHA RHA
LHA RHA
LHA RHA
1000 bp
750 bp
500 bp
250 bp
